# Enhancing HLA-DR in Cytotoxic T Lymphocytes is crucial for the development of efficient adoptive T cell Therapies for Breast Cancer

**DOI:** 10.1101/2024.09.03.610810

**Authors:** R Salvador, BF Correia, D Grosa, T Martins, SC Soares Baal, DP Saraiva, S Cristóvão-Ferreira, IL Pereira, C Rebelo de Almeida, R Fior, A Jacinto, C Mathias, S Braga, MG Cabral

## Abstract

**Background:** Despite advances in breast cancer (BC) therapies, more effective interventions are needed, especially for chemotherapy-resistant tumors. Immune checkpoint inhibitors show promise for triple-negative breast cancer, but their effectiveness across all BC subtypes remains challenging. Therefore, novel strategies, including adoptive cellular therapy, employing patients’ own T lymphocytes expanded *ex vivo*, are under investigation.

Previously, we demonstrated that cytotoxic T lymphocytes (CTLs) expressing high HLA-DR levels in the tumor microenvironment are associated with a good response to neoadjuvant chemotherapy (NACT), due to their pronounced anti-tumor properties compared to CTLs with low or no HLA-DR expression.

In this paper, we demonstrated that HLA-DR expression in CTLs is crucial for efficient T lymphocytes-based therapies.

**Methods:** To clarify the role of HLA-DR in CTLs’ anti-tumor abilities, we performed *in vitro* and *in vivo* experiments. We also improved a protocol to expand *ex vivo* HLA-DR-expressing CTLs and employed a 3D co-culture platform to test the potential of different immune agents, namely an anti-PD1, anti-OX40, anti-VEGF and anti-CD137, on CTLs cytotoxicity against BC cells. Additionally, we conducted a bioinformatic analysis of scRNA-seq data of BC patients to better understand the modulation of HLA-DR expression in CTLs.

**Results:** Our findings revealed that CTLs require HLA-DR expression to eliminate tumor cells. Additionally, we unveiled that blocking HLA-DR or depleting CD4+ T cells compromised CTLs activation and cytotoxicity, suggesting antigen presentation by CTLs through HLA-DR, and CD4+ T cells, as probable mechanisms for CTLs increased anti-tumor immune response and treatment efficacy.

We refined an *ex vivo* stimulation and cytokine supplementation protocol, observing that short-term stimulation increases HLA-DR expression while boosting CTLs functionality, unlike prolonged expansion. This result highlights the importance of prioritizing cell quality, over quantity, for therapy efficiency. Additionally, we verified that anti-PD-1 further increases HLA-DR levels in CTLs, enhancing their anti-tumor efficiency.

Notably, an *in silico* analysis revealed that PD-1 in CTLs shares 34 co-expressed genes with HLA-DR, including several non-coding RNAs, suggesting a PD-1-mediated regulation of HLA-DR expression.

**Conclusions:** Globally, our findings underscore that heightening HLA-DR expression in CTLs, by combining anti-PD-1 with short-term stimulation, offers promise for improving T lymphocyte-based therapies for BC.

**Key Message:**

- **What is already known on this topic:** While immunotherapy holds promise for breast cancer (BC), its success is still limited. Novel strategies are under investigation to improve outcomes across all BC subtypes. Previously we established a correlation between HLA-DR expression on CTLs and a positive response to neoadjuvant chemotherapy, likely due to enhanced anti-tumor properties of HLA-DR-expressing CTLs. However, the specific role of HLA-DR on CTLs and its implications for T lymphocyte-based therapies requires further investigation.
- **What this study adds**: This study revealed that HLA-DR expression is essential for CTLs to effectively eliminate tumor cells. Blocking HLA-DR or depleting CD4+ T cells impairs CTLs activation and cytotoxicity, indicating that HLA-DR-mediated antigen presentation and interaction with CD4+ T cell are crucial for CTLs function. The study also shows that combining ex vivo short-term stimulation and anti-PD-1 treatment increases HLA-DR levels in CTLs, enhancing their anti-tumor activity.
- **How this study might affect research, practice or policy**: By emphasizing the importance of optimizing CTLs quality over quantity, this approach has the potential to improve the design of more efficient T lymphocyte-based therapies for BC. This could influence future research directions, clinical practices, and treatment policies, leading to improved therapeutic outcomes.

## Introduction

Breast cancer (BC) is the most frequent type of cancer in women worldwide, accounting for up to 2 million new cases per year^1,2^.

One of the foremost challenges in addressing BC is its inherent heterogeneity, which influences the therapeutic options. Through specific biomarkers, including hormone receptor (HR) status and levels of the human epidermal growth factor receptor 2 (HER2), BC can be categorized into four subtypes: luminal A (HR+/HER2−), HER2+ (HR-/HER2+), luminal B (HR+/HER2+) and triple-negative (TNBC; HR−/HER2−). Each of these subtypes presents distinct risk factors for incidence, therapeutic response and disease progression^3,4^. Despite the advancements in surgery, chemotherapy, radiation and more recently, immunotherapy, the limitations of BC treatment persist^5, 6^.

Neoadjuvant chemotherapy (NACT) stands as a widely accepted approach for high-risk, locally advanced, or inoperable BC tumors^7^, though less than 50% of patients achieve a pathological complete response^8,9^. Previously, we proposed cytotoxic T lymphocytes (CTLs) expressing high levels of HLA-DR as a robust and independent biomarker for predicting the likelihood of BC patients successfully respond to NACT^10,11^. Nonetheless, the lack of effective options for BC patients with chemotherapy-resistant tumors poses a substantial clinical challenge, particularly for those ineligibles for prompt surgical intervention.

Recognizing the defies posed by immune escape, immunotherapies offer a promising avenue for cancer treatment, although their effectiveness can vary due to differences in patients’ immune competency and heterogeneity of tumors^12^. Questioned the prior assumption that breast tumors lack immunogenicity^13^, the scenario of BC treatment has been recently revolutionized by the introduction of immune checkpoint inhibitors (ICIs). ICIs targeting programmed death-1 (PD-1) or its ligand (PD-L1) are now established as a standard first-line of therapy for advanced or metastatic PD-L1 positive TNBC^14^. Furthermore, ICIs have also recently gained approval for high-risk early-stage TNBC^15^. Nowadays, several clinical trials are trying to expand immunotherapy to all BC subtypes, with a particular focus on early-stage cases, where it may be more effective, primary due to a less immunosuppressive tumor microenvironment^16^.

CAR-T cell therapy is also being explored as an alternative immune-based treatment for BC. This innovative approach involves engineering patient’s T-cells to express chimeric antigen receptors (CARs) that specifically target cancer cells. Recent studies have shown promising results in preclinical models, demonstrating the therapy’s ability to effectively recognize and eliminate BC cells. While CAR-T cell therapy has been primarily successful in treating hematologic malignancies, ongoing clinical trials are trying to optimize its use for solid tumors like BC. However, currently, there are no FDA-approved CAR-T cell therapies for solid tumors, including BC^17,18^.

Adoptive T cell therapy (ATC) has also emerged as a significant therapeutic strategy in the fight against cancer. This approach is continuously evolving and under evaluation, both as a standalone treatment or in combination with other immunotherapies through ongoing clinical trials^19^. Highlighting its potential in BC, a recent case demonstrated that the infusion of autologous T lymphocytes, which have been specifically designed to target neo-antigens and propagated *ex vivo*, was successfully employed in combination with ICI, in a patient with chemo-refractory metastatic BC^20^.

Despite the potential of ATC therapies in BC treatment, their integration into clinical practice poses significant challenges due to high costs, technical complexities and the toxicities related to excessive cytokine release^21^. Furthermore, while ICIs have made significant advances in BC, they are predominantly focused on addressing TNBC, leaving limited options for other cancer subtypes^16^. Nonetheless, the modulation of the immune system in BC patients appears to be an encouraging strategy for developing more personalized therapies.

Following the idea that HLA-DR-expressing CTLs are required in the tumor microenvironment to assist NACT in cancer elimination and exhibit anti-tumor properties, we suggested earlier that the upsurge of CTLs with increased levels of HLA-DR expression might be considered in novel therapeutic approaches for BC^11^. Thus, this paper further explores the therapeutic potential of HLA-DR-expressing CTLs and develops strategies to augment them. Specifically, our work underscores the relevance of HLA-DR expression levels in CTLs to monitor the quality of *ex vivo* manipulated cells, for the development of adoptive T cell transfer protocols. Furthermore, it demonstrates the synergistic effect of nivolumab, a monoclonal antibody targeting PD-1, and CTLs’ *ex vivo* short-term stimulation to elevate their HLA-DR expression and consequently enhance the cytotoxicity of these cells.

This synergy may be related to the fact that PD-1 in CTLs exhibits common correlated genes with HLA-DR, such as some known gene expression regulators, as revealed by our *in silico* analysis of BC scRNAseq databases. The combined strategy offers a promising avenue for integration into future T lymphocyte-based approaches, to amplify immune responses and achieve more favorable outcomes for BC patients.

## Methods

### Human Samples

This study was primarily performed using immune cells isolated from the buffy coats of 32 healthy donors, provided by the Instituto Português do Sangue e da Transplantação (IPST). Additionally, 209 samples were used from a cohort of patients diagnosed with breast cancer (BC) sourced from Hospital CUF Descobertas, Hospital CUF Tejo, Instituto Português de Oncologia de Lisboa Franscisco Gentil, and Hospital Prof. Doutor Fernando Fonseca. Specifically, blood samples from 14 BC patients with advanced/metastatic disease and biopsies from 195 BC patients selected for neoadjuvant chemotherapy (NACT) were utilized. Written informed consent from all individuals and approval from the Ethical Committees of the hospitals, IPST, and NOVA Medical School, were obtained. The study is in compliance with the Declaration of Helsinki. Patients’ main characteristics are detailed in supplementary table S1.

Patients’ blood samples were collected using Vacutainer tubes with ethylenediaminetetraacetic acid (EDTA, Vacutest). Fresh biopsies were collected in Transfix (Cytomark) to preserve cellular antigens, allowing flow cytometry analysis.

Peripheral blood mononuclear cells (PBMCs) were isolated from whole blood or buffy coats using a Ficoll gradient (Sigma Aldrich) and cryopreserved in a solution containing 90% fetal bovine serum (FBS, Biowest) and 10% dimethyl sulfoxide (DMSO, Sigma Aldrich) for later use.

The biopsies underwent mechanical dissociation using a BD Medicon (BD Biosciences), filtered through a 30 µm nylon mesh (Sysmex), washed with 1X Phosphate-Buffered Saline (PBS), and then stained for flow cytometry.

### Breast cancer cell lines

*In vitro* experiments were primarily carried out using the triple negative breast cancer (TNBC)-derived cell line MDA-MB-231. Additionally, some experiments were conducted with various BC cell lines, covering different BC subtypes, namely the BC cell lines BT-474 (HER-2+ subtype), HCC1806 (TNBC subtype), Hs578T (TNBC subtype) and MCF-7 (ER+ subtype). Hs578T, MCF-7, and MDA-MB-231 cell lines were cultured in Dulbecco’s Modified Eagle Medium (DMEM, Gibco) supplemented with 10% FBS (Biowest) and 1% Penicillin/Streptomycin (GE Healthcare). BT-474 and HCC1806 cell lines were cultured in RPMI 1640 (Gibco) supplemented with 10% FBS and 1% Penicillin/Streptomycin. Insulin (Sigma Aldrich) at 10µg/mL was additionally used in the culture of BT-474, Hs578T and MCF-7 cell lines. The cells were grown in monolayer cultures in T75 flasks under humidified conditions (37°C, 5% CO_2_) until they achieved 80-90% confluency. Subsequent cell passages were carried out by detaching the cells using TrypLE (Gibco).

Particularly, for *in vivo* experiments, Hs578T cancer cells were fluorescently labelled with TdTomato via lentiviral transduction. Cells were cultured in T175 flasks until 70-80% confluence and detached with EDTA 1 mM (diluted in DPBS 1X) and cell scrapers. Cell lines were tested routinely for mycoplasma contamination.

### Flow cytometry

For the flow cytometry analysis, biopsies, blood samples, or BC cells-PBMCs from co-cultures, were stained with a cocktail of monoclonal mouse anti-human conjugated antibodies (mAbs) after being processed to a single cell suspension. In the case of blood samples, the protocol included an additional step of red blood cell lysis using RBC lysis buffer (Biolegend) in the dark for 20 minutes at 4°C. Briefly, the staining protocol involved incubating cells with mAbs for 15 minutes at room temperature, followed by washing with 1 mL of PBS 1X and centrifugation at 300g for 5 minutes. Whenever intracellular markers were analyzed, cells were fixed and permeabilized with Fix/Perm kit (eBioscience) for 30 minutes at room temperature. Intracellular mAbs were added for 30 minutes, followed by a wash step with 1 mL of PBS 1X and centrifugation at 300g for 5 minutes.

The mAbs employed were: anti-CD45-PercP (clone HI30), anti-CD3-PercP (clone HIT3a), anti-CD3-APC (clone UCHT1), anti-CD25-PE (clone BC96), anti-CD4-FITC (clone OKT4), anti-CD8-PE (clone HIT8a), anti-CD8-PacificBlue (clone HIT8a), anti-HLA-DR-APC (clone L243), anti-Granzyme B-FITC (clone QA16A02), anti-CD69-PercP (clone FN50), anti-CD45-PercP (clone HI30), all from Biolegend. Cell viability was determined with BD Horizon^TM^ Fixable Viability Stain 450 (BD Biosciences) incubated with the cell suspension for 20 minutes in the dark at 4°C.

Data acquisition was conducted using the BD Fluorescence Activated Cell Sorting (FACS) Canto II with FACSDiva Software v8 (BD Biosciences), and the results were analyzed using FlowJo software v10.

In general, data are presented as a percentage of the positive population with respect to the single cells’ gate or the positive population for a given marker within the CD8+ cells.

### Fluorescence activated cell sorting (FACS)

To deplete CD4+ cells, PBMCs were stained with anti-CD45-PercP (clone HI30) and anti-CD4-FITC (clone OKT4). These cells were subsequently sorted into two distinct populations: CD45+/CD4+ and CD45+/CD4negative. For another experiment, PBMCs were sorted to isolate CD25+/HLA-DRnegative cells and CD25+/HLA-DR+ cells, after staining with anti-CD25-PE (clone BC96) and anti-HLA-DR-APC (clone L243). Sorting was performed using the FACS Aria III system from BD Biosciences with an efficiency above 90%.

### Establishment of 3D co-cultures

A 3D co-culture involving BC cell lines and PBMCs in a 1:3 ratio on agarose-coated plates was implemented, as previously established^22^. Before co-culture, PBMCs underwent stimulation, for 48h, with mouse anti-human anti-CD3 (5 μg/mL), anti-CD28 (1 μg/mL), and rat anti-mouse IgG1 (5 μg/mL) as the crosslinking antibody (Biolegend). Additionally, interleukin 2 (IL-2) at 100 UI/mL and interleukin 12 (IL-12) at 20 ng/mL (PeproTech) were added to the PBMCs monoculture during the last 24h of the stimulation protocol.

After 24h of co-culturing PBMCs with the BC cell lines, in some experiments (drug screening), therapeutic antibodies were introduced in the 3D co-culture. The spheroids were removed from the plate after 48h, dissociated by pipetting and stained with the Fixable Viability Dye to evaluate, by flow cytometry, the percentage of viable cancer cells in the co-culture. A pan-leukocyte marker, anti-CD45 (clone HI30, Biolegend) was also used to distinguish between tumor and immune cells.

### Expansion of HLA-DR+ Cytotoxic T lymphocytes

PBMCs were initially seeded in 24 multi-well plates at a density of 1×10^6^ cells/mL in 1 mL of RPMI 1640 Medium supplemented with 10% FBS and 1% Penicillin/Streptomycin. For T lymphocytes activation and expansion, 5 μg/mL of mouse anti-human anti-CD3, 1µg/mL of mouse anti-human anti-CD28 and 5μg/mL of rat anti-mouse IgG1 (Biolegend) were added to the PBMCs culture at day 0. After 24h of incubation, various cocktails of interleukins were added to the culture. IL-2 was consistently utilized at a concentration of 100 UI/mL, while IL-7, IL-12, and IL-15 were used at concentrations of 20 ng/mL (PeproTech). The cells were incubated at 37°C in a humidified atmosphere containing 5% CO_2_ for 14_days. To maintain optimal cell growth, every 2-3_days, half of the culture medium was replaced with fresh medium containing the interleukins.

Throughout the incubation, the cells were monitored and counted at seven time points (i.e., days 0, 3, 5, 7, 9, 11, 14), using trypan blue exclusion dye (GE Healthcare). When the wells became confluent, the culture was divided into new wells. To assess the final cell numbers for each condition, the cell counts from all the wells in that condition were summed after counting. For each condition, the frequency of HLA-DR+ cytotoxic T lymphocytes was monitored at the same time points, by flow cytometry.

### HLA-DR blocking Assay

Blockade of HLA-DR was performed using PBMCs obtained from healthy donors, cultured for 72h under different experimental conditions: without stimulation, with stimulation, and with stimulation along with an HLA-DR-blocking antibody. Stimulation was performed as described above, and for the HLA-DR blockade, 10µg/mL of purified anti-human HLA-DR (clone L243, Biolegend) was used. Following the 72h culture period, cells were harvested and labeled with anti-CD25 (clone BC96, Biolegend), anti-CD69 (clone FN50, Biolegend), and anti-Granzyme B (clone QA16A02, Biolegend), to assess the activation status of cytotoxic T lymphocytes.

### Fluorescence activated cell sorting (FACS)

To deplete CD4+ cells, PBMCs were stained with anti-CD45-PercP (clone HI30) and anti-CD4-FITC (clone OKT4). These cells were subsequently sorted into two distinct populations: CD45+/CD4+ and CD45+/CD4negative. For another experiment, PBMCs were sorted to isolate CD25+/HLA-DRnegative cells and CD25+/HLA-DR+ cells, after staining with anti-CD25-PE (clone BC96) and anti-HLA-DR-APC (clone L243). Sorting was performed using the FACS Aria III system from BD Biosciences with an efficiency above 90%.

### Agents Screening

Agents screening experiments were conducted using our established 3D co-culture platform^22^. Briefly, the BC cell lines BT-474, HCC1806, Hs578T, MCF-7, and MDA-MB-231 were challenged with PBMCs treated as described above, plus the addition of the following candidates: Nivolumab (Anti-PD-1, Bristol-Myers Squibb), Bevacizumab (Anti-VEGF, Roche), provided by the Champalimaud Foundation, 4-1BB (anti-137, clone 4B4-1, Biolegend), and anti-OX40 (Anti-134, clone Ber-ACT35, Biolegend). All the compounds were tested at a concentration of 50 µg/mL. The effect of the compounds on the cancer cells’ viability was assessed by flow cytometry, as referred above.

### Breast Cancer Zebrafish Xenograft Experiments

*In vivo* experiments were performed at Champalimaud foundation, using zebrafish model (*Danio rerio*) *Tg(fli1:GFP)*, which were maintained and handled in accordance with European Animal Welfare Legislation and Champalimaud Fish Platform Program.

At 48h post fertilization zebrafish larvae were anesthetized with Tricaine 1X and Hs578T TdTomato cells, alone or mixed with PBMCs at a ratio of 1:1, were microinjected into the perivitelline space (PVS)^23^. For injection, PBMCs were previously resuspended in CellTracker^(TM)^ Deep Red (1:1000 from the stock, Thermo Scientific) and incubated for 37°C for 10min, followed by 15min at 4°C.

After injection, xenografts were left on embryonic medium (E3) at 34°C. 24h post injection, successful injected zebrafish xenografts were sorted into classes according to their tumor size, being the E3 media renewed daily, as well as the removal of dead xenografts. At the end of the assay, xenografts were sacrificed and fixed in 4% (v/v) formaldehyde (FA) (Thermo Scientific) overnight at 4°C.

In the day after fixation, FA was removed, and the xenografts were permeabilized with PBS-Triton 0.1%. Nuclei were counterstaining with DAPI (0.05mg/mL) overnight 4°C. In the next day, DAPI was removed by sequential washes with PBS-Triton 0.1% and xenografts were mounted between two coverslips with MOWIOL mounting medium. All images were obtained using a Zeiss LSM710 fluorescence confocal microscope. Tumor size quantification was performed as previously described^24^.

### Bioinformatic Analysis

To investigate potential regulatory networks involving the *HLA-DR* and *PD1* genes, we used scRNA-seq data from BC patients, specifically focusing on the gene expression of CD8+ T lymphocytes. The data were extracted from the TIGER database (available at http://tiger.canceromics.org/#/), which serves as a repository for them. The analysis was performed using R version 4.4.0. The searching strategy for genes correlated with the *HLA-DR* and *PD1* was divided into three stages: a) search for genes positively/negatively correlated with the *HLA-DR gene*; b) search for genes positively/negatively correlated with the *PD1 gene*; c) and finally, search for commonly observed genes. We adopted the correlation value cutoff |>0.5|, p-value < 0.05 as a search criterion. To narrow down the search for shared targets between *HLA-DR* and *PD1*, only genes detected in the largest datasets were considered. Thus, for the genes correlated with *HLA-DR* and *PD1*, we used the genes that were significant in at least 3 datasets.

### Statistical Analysis

Statistical analysis was conducted using GraphPad Prism v8, and statistical significance was considered for *p-value <*0.05. Comparisons between samples were performed using a nonparametric Mann-Whitney test, paired t-test, or a two-way ANOVA with multiple comparisons.

## Results

### HLA-DR expression on CTLs allows breast cancer cells elimination

Previously, we reported that HLA-DR levels in CTLs, assessed in biopsies is a robust biomarker for predicting the response of BC to neoadjuvant chemotherapy (NACT)^10,11^ and developed a predictive probability of response algorithm^11^.

In this paper, we validated our predictive model in a new cohort of BC patients selected to NACT based on clinically established criteria, as the model accurately predicts most responses to this treatment (Supplementary Figure S1). Additionally, we demonstrated a positive correlation between HLA-DR levels in tumor infiltrating CTLs and HLA-DR levels in systemic CTLs^10^.

Then, to further investigate the anti-tumor effect of HLA-DR-expressing CTLs from the blood of BC patients, we conducted *ex vivo and in vivo* experiments (Figure 1A). Specifically, when stimulated PBMCs isolated from BC patients were added to a spheroid of the BC cell line MDA-MB-231^22^, it was observed an increase in tumor cell mortality, which was dependent on the frequency of HLA-DR-expressing CTLs within the PBMCs (Figure 1B). This establishes a positive correlation between the percentage of HLA-DR-expressing CTLs in the blood of BC patients and the ability of PBMCs to kill tumor cells (Spearman r=0.5429, p=0.0479, Figure 1B). Moreover, the addition of sorted HLA-DR-expressing CTLs to the BC spheroid, showed an enhanced cytotoxicity capacity in comparison to the addition of sorted CTLs without HLA-DR (p=0.0323, Figure 1C). Notably, both sorted populations exhibited equivalent levels of the classical activation marker CD25, highlighting the pivotal role of HLA-DR in augmenting CTLs-mediated tumor cell elimination.

**Figure 1.**
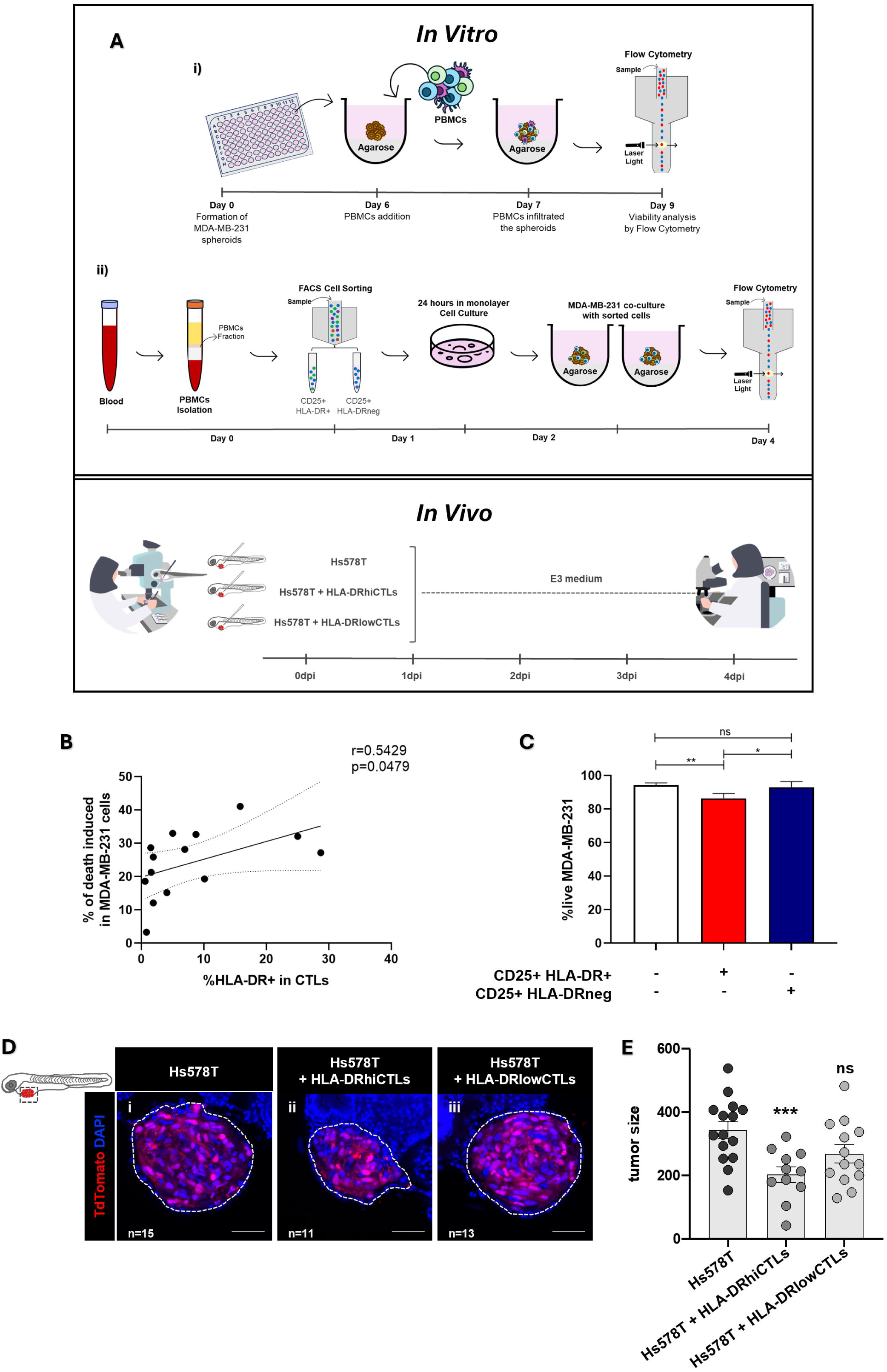
Breast Cancer cells elimination requires HLA-DR expression on CTLs. **(A)** Schematic representation of the experimental methodology employed. In the *in vitro* experiments: (i) the viability of breast cancer (BC) cell lines was analyzed via flow cytometry after the addition of peripheral blood mononuclear cells (PBMCs) derived from BC patients, which were previously stimulated. (ii) Fluorescence-activated cell sorting (FACS) was performed to separate two populations of activated immune cells (CD25+), one expressing HLA-DR (HLA-DR+) and the other lacking this molecule (HLA-DRneg). The cytotoxic capacity of both populations was assessed via flow cytometry after co-culturing with spheroids of BC cells^22^. In the *in vivo* experiments, zebrafish xenograft models were injected with a BC cell line alone or with a BC cell line plus immune cells, either from donors with high frequency of HLA-DRhiCTLs or with CTLs expressing low HLA-DR levels. Four days post-injection, the cytotoxic capacity of the immune cells was evaluated by analyzing the tumor size in the zebrafish, through confocal microscopy. **(B)** A positive correlation was observed between the percentage of CTLs expressing HLA-DR in BC patients’ blood and the efficacy of their PBMCs in eliminating tumor cells (Spearman r=0.5429, p=0.0479, n=14). **(C)** The viability of the MDA-MB-231 BC cell line (white bars, n=9) was significantly reduced upon the addition of sorted CD25+HLA-DR+CTLs, to the spheroids (red bar, p=0.0031, n=9), while the addition of CD25+HLA-DRnegHLA-DR did not lead to a decrease (blue bar, ns, n=9) of the cell line’s viability. **(D)** Representative confocal microscopy images showing a reduction in tumor size when xenografted zebrafish models were injected with BC patients’ PBMCs enriched in HLA-DRhiCTLs compared to those injected with PBMCs whose CTLs exhibited low HLA-DR levels (HLA-DRlowCTLs). Scale bars represent 50 µm. **(E)** Analysis of similar images to the one represented in D demonstrated that PBMCs from patients with increased frequency of HLA-DRhiCTLs exhibited a significant capacity to reduce tumor size (p<0.001, n=11), while PBMCs from patients with a higher frequency of HLA-DRlowCTLs did not affect tumor size (ns, n=13). Data are represented as mean ± SD*, *p < 0.05, **p < 0.01, ***p < 0.001, ns= non-statistical*.

These observations were further corroborated *in vivo,* through experiments with a zebrafish model that yielded comparable results to the spheroids. Specifically, xenografted zebrafish bearing the human BC cell line Hs578T, injected with PBMCs from BC patients with a high frequency of CTLs expressing high levels of HLA-DR showed a significant reduction in tumor size (p<0.001) compared to those injected with PBMCs from donors with low frequency of these cells (Figure 1D and 1E). These results showed that HLA-DR-expressing CTLs not only are a biomarker for NACT efficiency but also are more competent in promoting tumor cell death in vitro and in vivo.

### HLA-DR in CTLs and CD4+T cells are essential for effective CTLs activation and immune response

To explore the therapeutic potential of HLA-DR in CTLs, we investigated the activation and cytotoxic profile of stimulated CTLs when HLA-DR was blocked by a specific antibody.

Notably, blocking HLA-DR led to a reduction in the capacity of CTLs to become activated and cytotoxic, evidenced by decreased frequency of CD25+, CD69+ and Granzyme B+ cells within these population (Figure 2A). This underscores the pivotal role of HLA-DR in CTLs activation. However, since total PBMCs were used in our experiments, it is important to note that the blockade may affect HLA-DR expressed by other cells besides CTLs. Therefore, the observed results should be interpreted as reflecting this general blockade.

**Figure 2.**
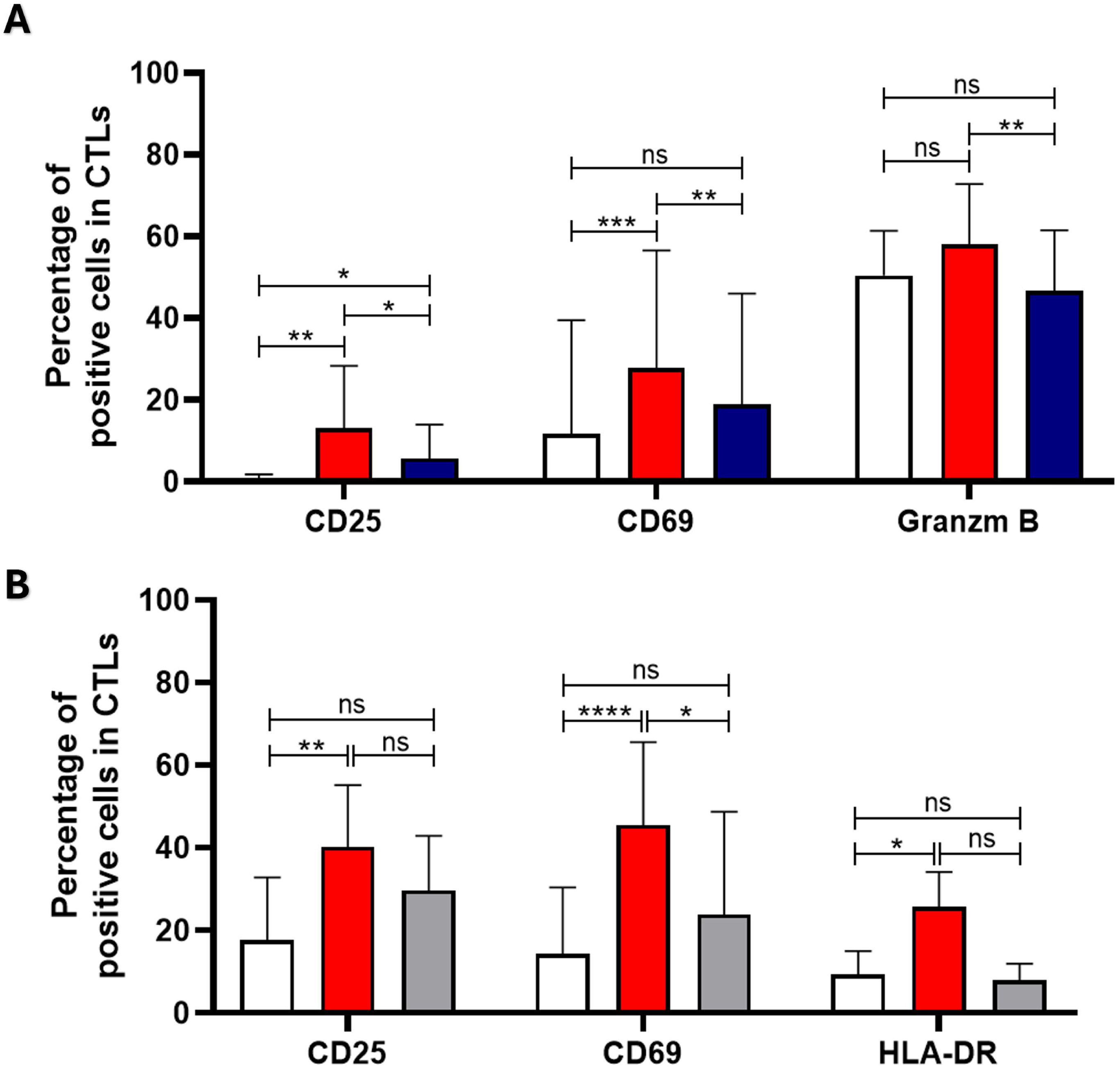
HLA-DR in CTLs and CD4+T cells are essential for effective CTLs activation and Immune Response. **(A)** The activation of CTLs following stimulation was confirmed by assessing the percentage of CD69+, CD25+ and Granzyme B+ cells within this population. Stimulation of PBMCs increases CTLs activation and cytotoxicity (red bars), compared to non-stimulatory conditions (white bars). HLA-DR blockade (blue bars) reduces the activation and cytotoxicity levels of CTLs, shown by decreased percentages of CTLs expressing CD25 (n=19, p=0.0262), CD69 (n=19, p=0.0050), and Granzyme B (n=14, p=0.0029). **(B)** PBMCs, depleted (grey bars) or not depleted (red bars) from CD4+ T cells, were stimulated, and the CTLs activation levels were assessed as previously. Following stimulation, the frequency of CTLs expressing the activation/cytotoxic markers was heightened compared to the basal state (white bars), in the condition with CD4+ T cells (red bars, n=9, p < 0.05). Nonetheless, stimulatory conditions barely affected the percentage of positive cells for the activation markers when CD4+ T cells were depleted (grey bars, n=6). Data are represented as mean ± SD, *\*p < 0.05, **p < 0.01, ***p < 0.001, ****p < 0.0001, ns= non-statistical*.

Additionally, considering that HLA-DR is the antigen presenting-molecule that enables antigen presentation to CD4+ T cells, we conducted experiments with PBMCs depleted of CD4+ T cells. Interestingly, stimulation of these PBMCs depleted of CD4+T cells had no significant effect on CTLs (Figure 2B), as indicated by the unchanged percentage of cells positive for various activation markers (grey bars), contrary to the conditions where CD4+ T cells were present (red bars, p < 0.05, Figure 2B). This observation highlights the importance of CD4+ T cells for robust CTLs activation.

Disrupting the interaction between CTLs and CD4+ T cells through HLA-DR blockade or CD4+ T cells depletion may compromise anti-tumor immune responses, suggesting a critical role for both HLA-DR and CD4+ T cells in CTLs activation and cytotoxic capabilities.

### Therapeutic Potential of CTLs could be boosted through Short-Term Expansion

Recent efforts have focused on refining the *ex vivo* expansion of T cells for therapeutic purposes. This pursuit aims to bolster their quantity for adoptive T cell transfer protocols and enhance their ability to recognize and destroy malignant cells, thereby improving treatment efficacy^25,26^.

In this context, we explored the dynamics of T cell expansion with a specific focus on HLA-DR-expressing CTLs. We evaluated various expansion protocols for immune cells, aiming to optimize their quantity and functionality. Our findings revealed that immune cells can be successfully expanded *ex vivo* for up to 14 days using T cell receptor stimulation combined with cytokine cocktails (Figure 3A). The most effective expansion protocol utilized anti-CD3 (5 μg/mL) and anti-CD28 (1 μg/mL) antibodies, along with interleukin (IL)-2 and IL-12 (Figure 3A). Therefore, we opted for this method in the subsequent experiments.

**Figure 3.**
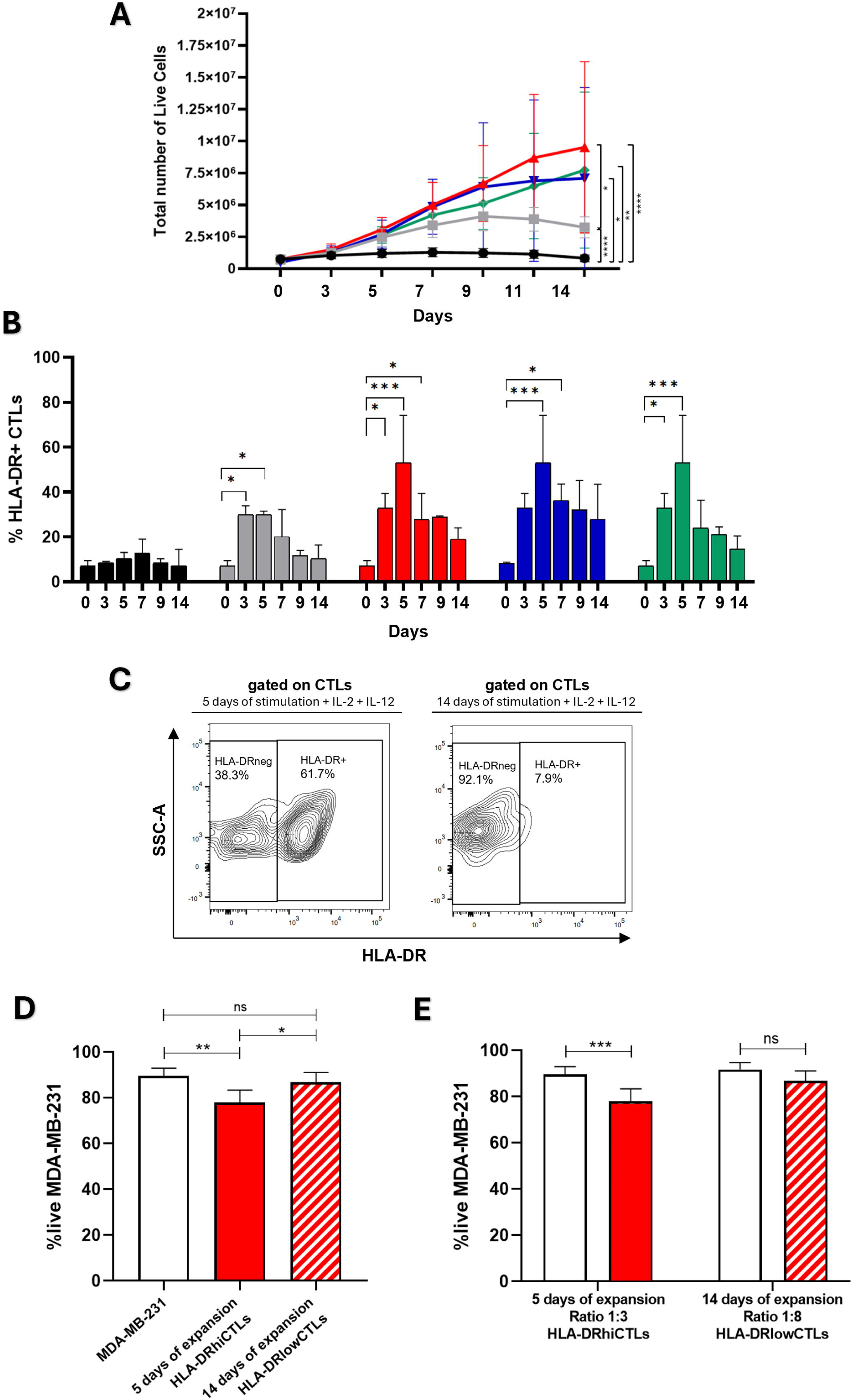
Therapeutic Potential of CTLs could be boosted through Short-Term Expansion. **(A)** Total number of live PBMCs during *ex vivo* expansion for up to 14 days. The cells survived for 14 days but did not proliferate without stimulation (black line, n=6). PBMCs stimulation with anti-CD3 (5 μg/mL) and anti-CD28 (1 μg/mL) slightly increased the number of viable cells for at least 10 days (grey line, n=6). The number of viable cells increased even more when the expansion protocol included the previous stimulation plus different cytokine cocktails, namely interleukin (IL)-2, IL-7, IL-12, and IL-15 (green line, n=4) or IL-2, IL-12, and IL-15 (blue line, n=4). The best expansion protocol was obtained with only IL-2 and IL-12 in addition to the anti-CD3 and anti-CD28 (red line, n=6). **(B)** Percentage of HLA-DR-expressing CTLs under different stimulating conditions. This percentage increased in the initial 5 days of *ex vivo* expansion and decreased after this period across all the conditions (n=4). Grey bars refer to the unstimulated condition. Black bars refer to the condition where PBMCs were stimulated with anti-CD3 (5 μg/mL) and anti-CD28 (1 μg/mL), green bars refer to the condition of stimulation with anti-CD3 and anti-CD28 in combination with IL-2, IL-7, IL-12, and IL-15. Blue bars refer to the condition of stimulation with anti-CD3 and anti-CD28 plus IL-2, IL-12, and IL-15. Finally, red bars represent PBMCs stimulated with IL-2 and IL-12 in addition to the two stimulatory antibodies. **(C)** Representative image of a contour plot illustrating that a short-term stimulation protocol with anti-CD3 and anti-CD28 plus IL-2 and IL-12 (5 days - left side) led to a significant increase in HLA-DR expressing CTLs, contrary to prolonged incubation (14 days - right side) with the same reagents. **(D)** *In vitro* 3D co-cultures composed of BC cell line MDA-MB-231 in a spheroid structure alone (white bar, n=4), with PBMCs short-term stimulated (5 days) (red bar, n=9), or with PBMCs long-period stimulated (14 days) (red striped bar, n=9). Short-term stimulation intensifies the cytotoxic activity of the PBMCs, as the viability of MDA-MB-231 cells was significantly decreased when co-cultured with PBMCs expanded using short-term stimulation protocol (p=0.0036), but barely affected when co-cultured with PBMCs stimulated for a superior period. **(E)** *In vitro* 3D co-cultures composed of BC cell line MDA-MB-231 in a spheroid structure alone (white bars), with PBMCs short-term stimulated (5 days), added to the culture at a ratio of 3 PBMCs:1BC cell (red bars, n=5) and the BC cell line MDA-MB-231 in a spheroid structure alone (white bars), with PBMCs long-period stimulated (14 days), added to the culture at a ratio of 8 PBMCs:1BC cell (red striped bar, n=5). The viability of the BC cell line was significantly reduced when co-cultured with PBMCs stimulated for a shorter period, even at a lower ratio, while the viability of the BC cell line was not influenced by higher numbers of PBMCs stimulated during more days. Data are represented as mean ± SD, *\*p < 0.05, **p < 0.01, ***p < 0.001, ****p < 0.0001, ns= non-statistical*.

However, while total numbers of PBMCs increased during the expansion protocol, we observed that specifically HLA-DR-expressing CTLs increased during the first days but suffered a decline after 4 to 5 days, regardless of the cytokine cocktail employed (Figure 3B). Interestingly, our results also demonstrated that a short-term stimulation protocol (5 days) led to an upregulation of HLA-DR expression on CTLs compared to prolonged expansion method (14 days) (Figure 3C). Additionally, experiments employing 3D co-culture models unveiled that cells expanded for longer periods (14 days) exhibited diminished cytotoxic capacity compared to cells submitted to short-term stimulation (5 days) (Figure 3D). Notably, we also demonstrated that cells expanded for a longer period exhibited diminished cytotoxic capacity even when added to BC cells in a higher ratio (8 PBMCs to 1 BC cell), compared to cells expanded for a shorter duration, which were added to BC cells only at a ratio of 3 PBMCs to 1 BC cell. Indeed, PBMCs expanded for 14 days, despite their superior number (8000), did not lead to a significant decrease in the viability of MDA-MB-231 cells (Figure 3E), contrary to the less expanded PBMCs, which were present in reduced numbers (3000) in the co-culture.

This higher cytotoxic capacity exhibited by less expanded PBMCs is most likely because short stimulation led to more HLA-DR-expressing CTLs than long stimulation, as shown in Figure 3C.

These results underscore the importance of prioritizing expansion protocols that guarantee the effective cytotoxic capabilities of CTLs, which we demonstrated to be strongly dependent on HLA-DR expression, achieved through short-term simulation.

### Anti-PD-1 treatment contributes to increment HLA-DR in CTLs and amplifies their Anti-Tumor activity

Using the 3D co-culture model we established ^22^ as a screening platform, we tested potential agents aimed at augmenting CTLs’ HLA-DR expression and consequently their cytotoxicity against BC cells, in line with the results previously shown.

Our screening efforts encompassed several promising agents and different BC cell lines (Table S1). Following a previous characterization of the immune profile of HLA-DR+ CTLs compared to HLA-DRneg CTLs revealing that several co-receptors are more expressed in HLA-DR+ CTLs (Figure S2), we thought that using agonists targeting these molecules could be an interesting way to turn HLA-DRneg CTLs into HLA-DR+ CTLs. Therefore, in our screening, we included two of these candidates, namely an anti-CD137 and an anti-CD134/OX40, which have been under evaluation in several clinical trials.

Additionally, we included Nivolumab, an immune checkpoint inhibitor drug that targets PD-1, boosting the immune response against cancer cells, via PD-1/PDL-1 axis blockade, and Bevacizumab, a monoclonal antibody that targets and neutralizes vascular endothelial growth factor (VEGF), but which has also been reported as having immunostimulatory properties.^27,28^ Both have been used extensively in clinical practice for different solid cancers.

Nivolumab significantly potentiated the anti-tumor properties of activated CTLs against MDA-MB-231 and HCC-1806, while the other tested drugs did not show any effect on improving cytotoxicity against these BC cancer lines (p=0.0163, Figure 4A; Figure S3 and Table S1). We observed consistent results with PBMCs sourced from healthy donors and PBMCs obtained from patients with metastatic BC (p=0.0487, Figure 4C), both of which have low or null basal levels of HLA-DR in CTLs.

**Figure 4.**
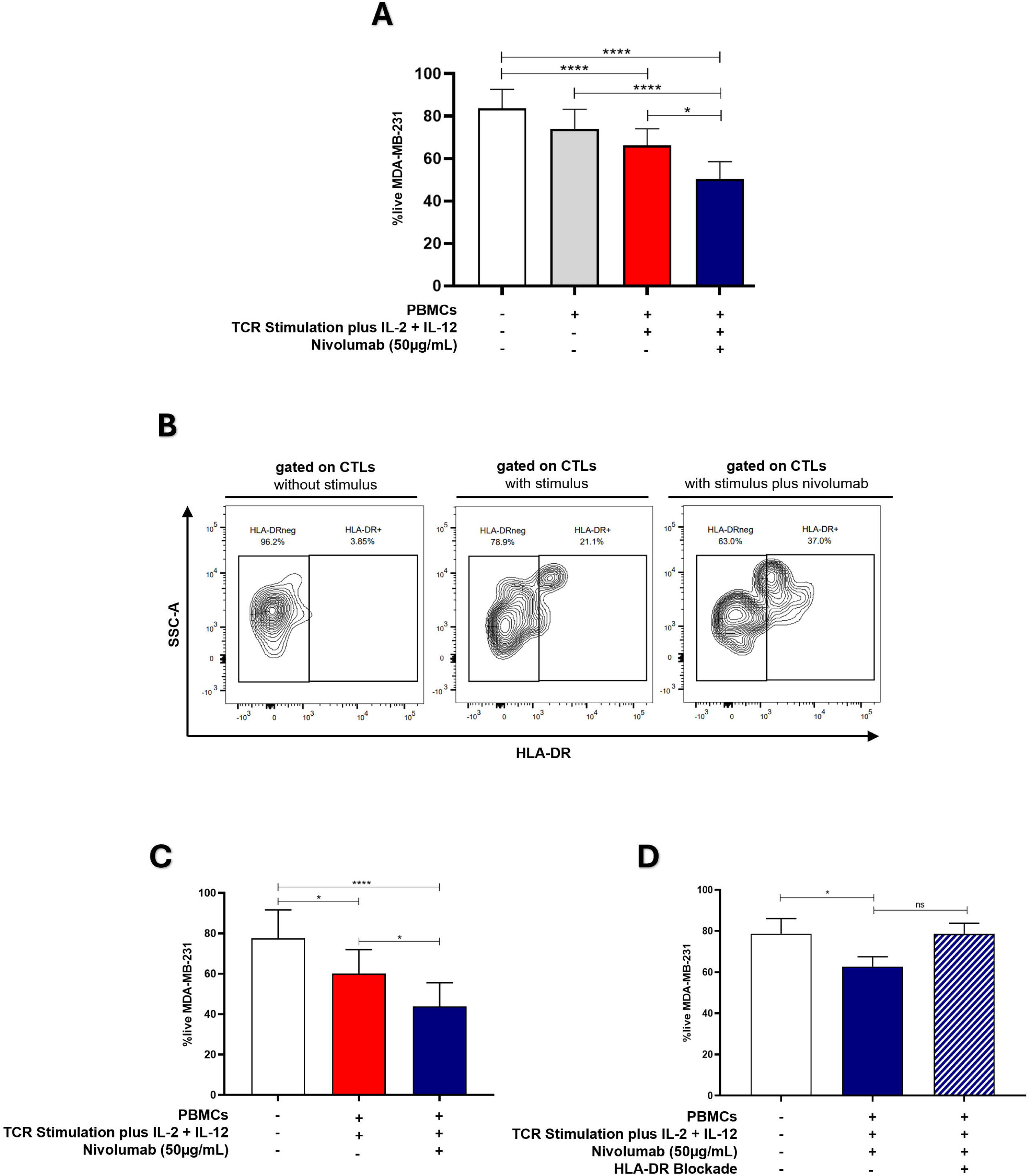
Anti-PD-1 treatment potentiates the anti-tumor capacity of cytotoxic T lymphocytes (CTLs) against Breast Cancer (BC). **(A)** The viability of the MDA-MD-231 BC cell line in the 3D co-culture alone (white bar, n=27), incubated in the presence of unstimulated PBMCs (grey bar, n=27), incubated in the presence of stimulated PBMCs (red bar, n=27) and incubated in the presence of stimulated PBMCs plus the addition of the anti-PD-1, Nivolumab (blue bar, n=22). The results demonstrated that the cytotoxic capacity against cancer cells was synergistically enhanced with the stimulation protocol previously established and the addition of Nivolumab to the co-culture (p=0.0163). **(B)** Representative contour plot showing that Nivolumab enhances the frequency of CTLs that express HLA-DR. **(C)** The viability of the MDA-MD-231 BC cell line alone (white bar, n=27), incubated in the presence of stimulated PBMCs which have been isolated from metastatic BC patients (red bar, n=14) and incubated in the presence of those stimulated PBMCs plus Nivolumab (blue bar, n=14, p=0.0487). These results were consistent with the results observed in (A) using PBMCs obtained from healthy donors. **(D)** The viability of the MDA-MD-231 BC cell line alone (white bar, n=5), incubated in the presence of stimulated PBMCs plus Nivolumab (blue bar, n=5, p=0.0159), and incubated in the presence of stimulated PBMCs plus Nivolumab and an anti-HLA-DR blocking antibody (blue striped bar, n=5, non-statistical). This demonstrates that blocking HLA-DR impairs cytotoxicity against BC cells, even in the presence of Nivolumab. Data are represented as mean ± SD, *\*p < 0.05, ****p < 0.0001, ns= non-statistical*.

Notably, our results also indicated that Nivolumab treatment *in vitro*, increases the percentage of HLA-DR+CTLs among PBMCs (Figure 4B) and that BC patients submitted to anti-PD-1 therapy also exhibited an elevated frequency of CTLs expressing HLA-DR in their blood, 4 months after initiating this treatment (Figure S4).

Supporting our finding that Nivolumab boosts CTLs cytotoxicity, through HLA-DR increment, HLA-DR blockade in our 3D co-cultures dampened the effect of Nivolumab, resulting in a compromised cytotoxic capacity of CTLs (Figure 4D).

Overall, our findings suggest that anti-PD-1 treatment holds considerable potential in boosting CTLs cytotoxicity through HLA-DR, beyond mere stimulation, especially against BC subtypes with elevated PD-L1 expression (Figure 4A, Figure S3, Table S1).

Although immunotherapy with anti-PD-1 in BC has primarily been applied for TNBC cases, considered the most immunogenetic subtype, our analysis of immune features in patients’ biopsies from TNBC, ER+ and HER2+ BC subtypes, revealed similar frequencies of infiltrating CTLs and comparable PD-L1 expression across all subtypes (Figure S5). This suggests that the proposed approach could potentially be applied to all BC subtypes. Interestingly, since ER+ and HER2+ exhibited fewer HLA-DR-expressing infiltrating CTLs, these two subtypes may benefit even more from this type of treatment.

### Several ncRNAs are correlated with the expression of HLA-DR and PD1 genes in CTLs

To investigate potential regulatory networks involving the *HLA-DR* and *PD1* genes, we used scRNA-seq data from BC patients extracted from the TIGER database. We focused on gene expression data from CD8+ T lymphocytes. Detailed gene lists for each dataset can be found in the supplementary material. We did not find significant correlation values regarding negatively correlated genes for both *HLA-DR* and *PD1*, therefore, our analysis only considered positively correlated genes with *both* genes.

For the *HLA-DR* gene, we found five scRNA-seq datasets from CD8+ T cells, named BRCA_GSE150660, BRCA_GSE110686, BRCA_GSE114727_10X, BRCA_GSE161529 and BRCA_EMTAB8107 (Figure 5A). Considering these datasets, we identified genes significantly correlated with HLA-DR (r > 0.5; p < 0.05) across the largest number of datasets, highlighting those with potential biological significance within the regulatory network of interest. The same analysis logic was applied to the *PD1 (PDCD1)* gene, based on four scRNA-Seq datasets, namely: BRCA_GSE110686, BRCA_GSE114727_10X, BRCA_GSE161529 and BRCA_EMTAB8107 (Figure 5B). The information about genes and their correlation value is present in supplementary tables S3 to S11. Our analysis revealed 34 genes positively correlated with both HLA-DR and PD1 (Figure 5C).

**Figure 5.**
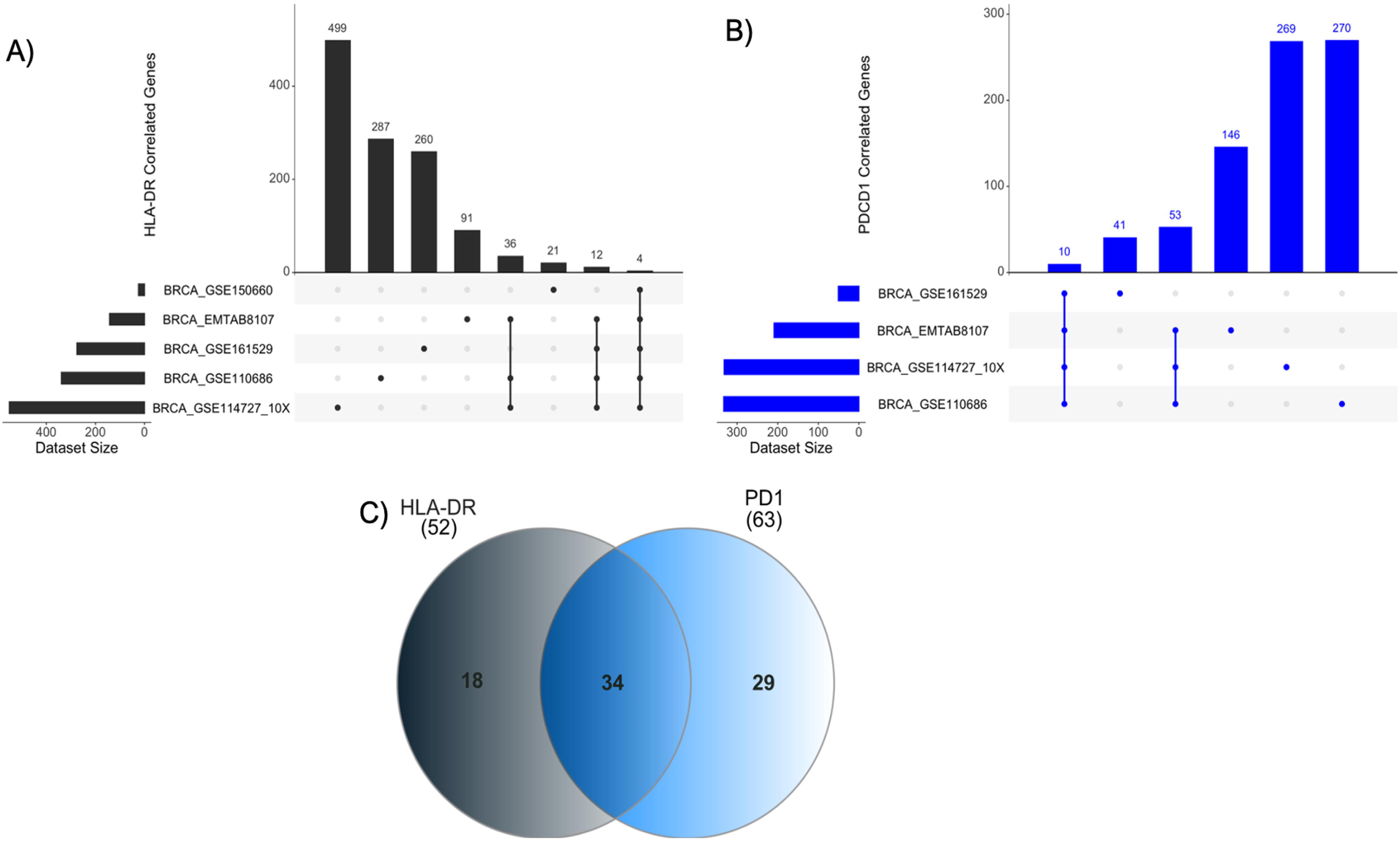
Upset plots representing the number of genes positively correlated with HLA-DR and PD1 in the analyzed datasets. **(A)** For HLA-DR, the dataset BRCA_GSE114727_10x was the one with the highest number of genes correlated with HLA-DRA (n=499), and the dataset BRCA_GSE150660 the one with the lowest number (n=21). It is also possible to highlight that four genes (CD74, CCL4, CCR5, and COTL1) were present in all analyzed datasets. **(B)** In the same way, for PD1 (PDCD1), the datasets BRCA_GSE110686 and BRCA_GSE114727_10X were the ones with the highest number of genes (n=270 and n=269, respectively. Among all datasets, 10 genes were shared between them: TIGIT, TNFRSF9, IFNG, CXCR6, CTLA4, ZBED2, CD2, CLEC2D, CD3D and PTPN7. **(C)** Venn diagram representing exclusive and shared genes among HLA-DR and PD1. 34 positively correlated genes are shared among these genes.

Notably, several non-coding RNAs (ncRNAs) were among these correlated genes. Specifically, four of the five datasets that include 52 genes for HLA-DR, found a positive correlation with ncRNAs, specifically with three long non-coding RNA (lncRNAs). For PD1, considering the 63 genes shared at least in three datasets, we also detected a positive correlation with lncRNAs. In Table 1 we present all lncRNAs that have a positive correlation with HLA-DR or PD1. In bold we highlight two lncRNAs, MIR155HG and KCNQ1OT1, that were consistently shared, in the different datasets, which may have a role in the regulatory axis of HLA-DR expression in response to anti-PD1 treatment.

**Table 1.** Long non-coding RNAs (lncRNAs) positively correlate with HLA-DR and PD1 in each dataset. The lncRNAs MIR155HG and KCNQ1OT1 are highlighted in bold since they are genes shared between HLA-DR and PD1.

| **Dataset** | **Gene 1** | **Gene 2** | **Correlation value** |
| --- | --- | --- | --- |
| BRCA_GSE110686 | HLA-DR | **MIR155HG** | 0.79 |
|  |  | **KCNQ1OT1** | 0.81 |
|  |  | LINC01480 | 0.54 |
|  | PD1 | **MIR155HG** | 0.79 |
|  |  | **KCNQ1OT1** | 0.79 |
|  |  | AC133644.2 | 0.79 |
|  |  | LINC01480 | 0.60 |
| BRCA_GSE114727_10X | HLA-DR | **MIR155HG** | 0.68 |
|  |  | **KCNQ1OT1** | 0.63 |
|  | PD1 | **MIR155HG** | 0.85 |
|  |  | **KCNQ1OT1** | 0.78 |
|  |  | AC133644.2 | 0.74 |
|  |  | AC017002.1 | 0.67 |
|  |  | LINC01480 | 0.61 |
| BRCA_GSE161529 | HLA-DR | **MIR155HG** | 0.52 |
|  |  | LINC01857 | 0.55 |
|  | PD1 | LINC01943 | 0.56 |
| BRCA_EMTAB8107 | HLA-DR | **KCNQ1OT1** | 0.67 |
|  |  | **MIR155HG** | 0.59 |
|  | PD1 | **MIR155HG** | 0.76 |
|  |  | **KCNQ1OT1** | 0.74 |
|  |  | LINC01480 | 0.70 |
|  |  | AC133644.2 | 0.66 |

## Discussion

Projections indicate a substantial increase in Breast Cancer (BC) incidence and mortality rates in the coming years, underscoring the urgent need for collective efforts to mitigate this growing burden^29^.

Immunotherapy emerged as a promising approach to treat BC. Though, challenges such as tumor heterogeneity and immune evasion persist, hindering the advancement of this therapeutic avenue^30–32^. The success of the KEYNOTE-355 trial^33^ and the subsequent regulatory approval of pembrolizumab for triple-negative breast cancer (TNBC) have spurred additional clinical trials exploring alternative immunotherapy modalities, such as adoptive T cell therapies, across all BC subtypes^34,35^.

Despite recent advancements, a significant portion of BC patients, especially those with advanced disease, fail to respond to conventional and immunotherapy treatments, often due to the suboptimal effectiveness of cytotoxic T lymphocytes (CTLs).

Previously, we demonstrated that the presence of HLA-DR-expressing CTLs in the tumor microenvironment is crucial for a favorable response to neoadjuvant chemotherapy (NACT) in BC patients^10,11^, likely because this molecule enhances the anti-tumor activity of these cells. Notably, our findings are consistent with other studies indicating the importance of HLA-DR expression for effective response to NACT in BC patients, associated with increased IL-12 and IFN-γ plasma levels^10,11,36^.

In the current study, we further validated, in a new cohort of BC patients, our predictive model of the probability of response to NACT, based on HLA-DR levels in tumor infiltrating CTLs^11^, reinforcing the accuracy of this biomarker for the clinical setting.

Additionally, employing 3D co-culture and zebrafish xenograft models, we established that PBMCs enriched with HLA-DR-expressing CTLs exhibited superior cytotoxicity against BC cells compared to PBMCs with few or no HLA-DR-expressing CTLs. These results further support that the development of strategies aimed at increasing HLA-DR expression in CTLs should be of therapeutic importance to enhance the anti-tumor immune response against BC.

Our experiments revealed that blocking HLA-DR or depleting CD4+ T cells reduced CTLs activation and cytotoxicity. Given that HLA-DR+CTLs possess the machinery for antigen processing and loading onto HLA-DR molecules^37^ and that T cell-T cell synapsis were described^38^, we hypothesize that HLA-DR-mediated antigen presentation not only primes CTLs for efficient recognition and elimination of target cells but also reinforces their activation through interactions with CD4+ T cells. These CD4+ T cells then release cytokines, such as IFN-γ, ensuring a more robust and sustained CTLs activity (Figure 6).

**Figure 6.**
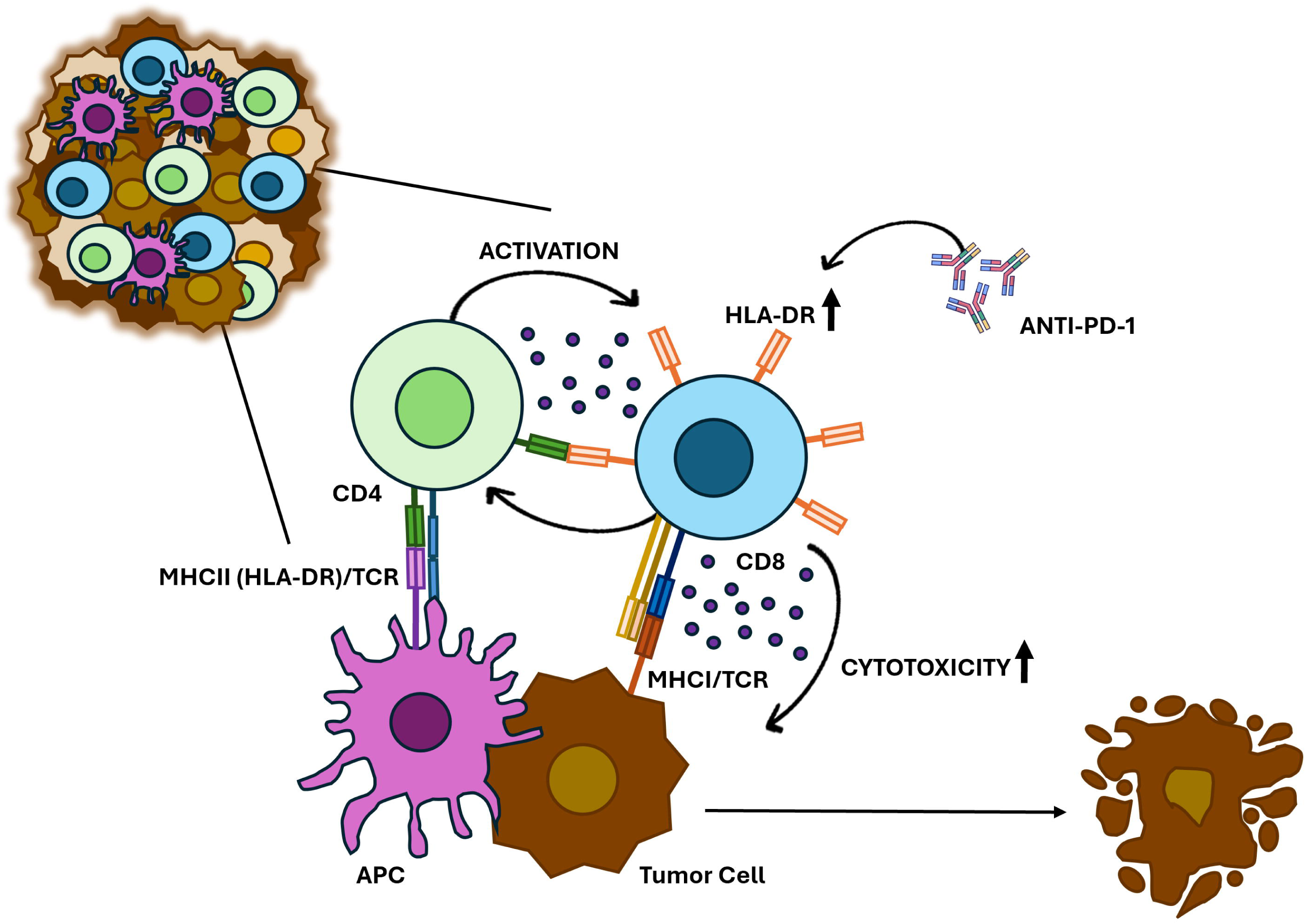
Proposed model for HLA-DR-expressing CTLs in anti-cancer immune response. HLA-DR-mediated antigen presentation primes CTLs to recognize and eliminate breast cancer cells. HLA-DR-expressing CTLs may also interact with CD4+ T cells, which release cytokines such as IFN-γ, reinforcing CTL activation and sustaining a robust immune response against tumors. Additionally, anti-PD-1 treatment increases the HLA-DR expression in CTLs and significantly boosts their cytotoxicity.

Recent studies have been exploring strategies to enhance tumor immunity against cancer cells. For instance, it was demonstrated in pre-clinical models that targeting IL-2 to CD8+ T cells promotes robust effector T-cell responses and potent antitumor immunity^39^. In this work we investigated ways to expand the population of HLA-DR-expressing CTLs.

We concluded that PBMCs short-term stimulation, together with IL-2 and IL-12 supplementation, significantly enhances the frequency of these cells, which correlates with their improved cytotoxic capacity against BC cells. In contrast, prolonged PBMCs stimulation increases the total number of cells but results in diminished HLA-DR-expressing CTLs and reduced cytotoxicity.

Notably, the addition of Nivolumab, an anti-PD-1 antibody, synergistically increased the percentage of HLA-DR expression in CTLs and boosted the cytotoxicity of PBMCs population. This suggests that treating PBMCs with anti-PD-1 could enhance CTL-mediated anti-tumor responses, not only because this antibody blocks inhibitory signals from PD1/PD-L1 interactions, but also because it upregulates HLA-DR expression, probably promoting a positive feedback loop of cytokine release and sustained activation (Figure 6).

Both HLA-DR and PD-1 have been extensively studied in tumor biology contexts^36–40;^ however, little is known about HLA-DR role in tumor-infiltrating CTLs, or regarding the links between both molecules. For instance, a study by Heng Yu et al (2023)^45^ on laryngeal squamous cell carcinoma found that high expression of tumor HLA-DR is associated with improved response to anti-PD-1 therapy^45^, but this research focused on HLA-DR expressed by tumor cells and not specifically on CTLs.

Although it is known that upon anti-PD-1 treatment, PD-1 expression typically increases in lymphocytes as part of the normal activation process^46,47^, the interplay between this molecule and HLA-DR, which also rises in lymphocytes under stimulatory conditions, remains unclear.

Therefore, aiming at investigate shared regulatory pathways between HLA-DR and PD1 genes, we conducted a data analysis using scRNA-Seq data. Thirty-four genes were identified that showed a positive correlation with both genes. A noteworthy finding was the presence of long non-coding RNAs (lncRNAs) in this list. lncRNAs are transcripts longer than 500 nucleotides^48^, which are now recognized for their potential to encode small functional peptides^49, 50^. Specifically, the lncRNAs MIR155HG and KCNQ1OT1 were positively correlated with both HLA-DR and PD1, exhibiting high correlation values. These data might suggest that these genes are involved in the regulatory axis involving the expression of HLA-DR in response to modulations in PD-1 expression, highlighting the relevance of more in-depth studies in the future.

Overall, our study underscores the crucial role of HLA-DR expression in CTLs for the elimination of BC cells. We hypothesize that manipulating HLA-DR expression, through combined approaches, could significantly augment CTLs-mediated tumor killing, particularly for BC patients with lower HLA-DR expression in CTLs, potentially applied across all the BC subtypes.

These findings pave the way for the development of improved personalized therapeutic interventions with patients’ own immune cells. Ultimately, the proposed approach holds the potential to reshape the landscape of BC treatments, thereby contributing to increased BC-free survival rates.

## Supporting information

https://eur01.safelinks.protection.outlook.com/?url=https%3A%2F%2Fdrive.google.com%2Fdrive%2Ffolders%2F1GQdUyL_CFZbh8kN3N2J6MufQbc_j8r3z%3Fusp%3Ddrive

## Ethics approval and consent to participate

Written informed consent was obtained from all individuals involved in the study. The study received approval from the Ethical Committees of the participating hospitals, Instituto Português do Sangue e da Transplantação and NOVA Medical School.

## Consent for publication

Consent for publication was obtained from all participants and authors included in the study.

## Availability of data and material

Not applicable.

## Author contributions

RS conducted the entirety of the experiments, including analysis and interpretation of data, statistical analysis, figure assembly, and manuscript writing. DPS, BFC, and DG helped in conducting the flow cytometry experiments. RF and CRA developed experiments using the zebrafish model. AJ actively contributed to scientific discussion. SB, SCF, ILP and TM played a key role in obtaining patients’ samples and clinical data and scientific discussion. CM and SCSB developed the bioinformatics analysis. MGC designed and supervised the study, contributed to data interpretation, and participated in manuscript writing. All authors contributed to manuscript revisions. The final manuscript received approval from all authors.

## Acknowledgements

We would like to thank to all the breast cancer patients that agreed to participate in this study. Our sincere appreciation goes to the dedicated professionals—Radiologists, Pathologists, Oncologists, Surgeons, and Nurses—at Hospital CUF Descobertas, Hospital CUF Tejo, Instituto Português de Oncologia de Lisboa Franscisco Gentil and Hospital Prof Doutor Fernando Fonseca, whose invaluable assistance played a crucial role in the collection of samples. We also thank the Instituto Português do Sangue e da Transplantação (IPST) and all healthy donors. Additionally, we extend our thanks to the Flow Cytometry Facility and Cell Culture Facility of NOVA Medical School.

## References

1. Sung H, Ferlay J, Siegel RL, et al. Global Cancer Statistics 2020: GLOBOCAN Estimates of Incidence and Mortality Worldwide for 36 Cancers in 185 Countries. CA Cancer J Clin. 2021;71(3):209–249. doi:10.3322/caac.21660

2. Giaquinto AN, Sung H, Miller KD, et al. Breast Cancer Statistics, 2022. CA Cancer J Clin. 2022;72(6):524–541. doi:10.3322/caac.21754

3. Orrantia-Borunda E, Anchondo-Nuñez P, Acuña-Aguilar LE, Gómez-Valles FO, Ramírez-Valdespino CA. Subtypes of Breast Cancer. In: Breast Cancer. Exon Publications; 2022:31–42. doi:10.36255/exon-publications-breast-cancer-subtypes

4. Tong CWS, Wu M, Cho WCS, To KKW. Recent Advances in the Treatment of Breast Cancer. Front Oncol. 2018;8. doi:10.3389/fonc.2018.00227

5. Burguin A, Diorio C, Durocher F. Breast Cancer Treatments: Updates and New Challenges. J Pers Med. 2021;11(8):808. doi:10.3390/jpm11080808

6. Tufail M, Cui J, Wu C. Breast cancer: molecular mechanisms of underlying resistance and therapeutic approaches. Am J Cancer Res. 2022;12(7):2920–2949. http://www.ncbi.nlm.nih.gov/pubmed/35968356

7. Thompson AM, Moulder-Thompson SL. Neoadjuvant treatment of breast cancer. Ann Oncol. 2012;23:x231–x236. doi:10.1093/annonc/mds324

8. DeMichele A, Yee D, Esserman L. Mechanisms of Resistance to Neoadjuvant Chemotherapy in Breast Cancer. Phimister EG, ed. N Engl J Med. 2017;377(23):2287–2289. doi:10.1056/NEJMcibr1711545

9. Cortazar P, Zhang L, Untch M, et al. Pathological complete response and long-term clinical benefit in breast cancer: the CTNeoBC pooled analysis. Lancet. 2014;384(9938):164–172. doi:10.1016/S0140-6736(13)62422-8

10. Saraiva DP, Jacinto A, Borralho P, Braga S, Cabral MG. HLA-DR in Cytotoxic T Lymphocytes Predicts Breast Cancer Patients’ Response to Neoadjuvant Chemotherapy. Front Immunol. 2018;9. doi:10.3389/fimmu.2018.02605

11. Saraiva DP, Azeredo-Lopes S, Antunes A, et al. Expression of HLA-DR in Cytotoxic T Lymphocytes: A Validated Predictive Biomarker and a Potential Therapeutic Strategy in Breast Cancer. Cancers (Basel). 2021;13(15):3841. doi:10.3390/cancers13153841

12. Wang S, Xie K, Liu T. Cancer Immunotherapies: From Efficacy to Resistance Mechanisms – Not Only Checkpoint Matters. Front Immunol. 2021;12. doi:10.3389/fimmu.2021.690112

13. Jacob SL, Huppert LA, Rugo HS. Role of Immunotherapy in Breast Cancer. JCO Oncol Pract. 2023;19(4):167–179. doi:10.1200/OP.22.00483

14. Howard FM, Villamar D, He G, Pearson AT, Nanda R. The emerging role of immune checkpoint inhibitors for the treatment of breast cancer. Expert Opin Investig Drugs. 2022;31(6):531–548. doi:10.1080/13543784.2022.1986002

15. Dixon-Douglas J, Loi S. Immunotherapy in Early-Stage Triple-Negative Breast Cancer: Where Are We Now and Where Are We Headed? Curr Treat Options Oncol. 2023;24(8):1004–1020. doi:10.1007/s11864-023-01087-y

16. Debien V, De Caluwé A, Wang X, et al. Immunotherapy in breast cancer: an overview of current strategies and perspectives. npj Breast Cancer. 2023;9(1):7. doi:10.1038/s41523-023-00508-3

17. Shafer P, Kelly LM, Hoyos V. Cancer Therapy With TCR-Engineered T Cells: Current Strategies, Challenges, and Prospects. Front Immunol. 2022;13. doi:10.3389/fimmu.2022.835762

18. Chamorro DF, Somes LK, Hoyos V. Engineered Adoptive T-Cell Therapies for Breast Cancer: Current Progress, Challenges, and Potential. Cancers (Basel). 2023;16(1):124. doi:10.3390/cancers16010124

19. Du S, Yan J, Xue Y, Zhong Y, Dong Y. Adoptive cell therapy for cancer treatment. Exploration. 2023;3(4). doi:10.1002/EXP.20210058

20. Zacharakis N, Chinnasamy H, Black M, et al. Immune recognition of somatic mutations leading to complete durable regression in metastatic breast cancer. Nat Med. 2018;24(6):724–730. doi:10.1038/s41591-018-0040-8

21. Vasileiou M, Papageorgiou S, Nguyen NP. Current Advancements and Future Perspectives of Immunotherapy in Breast Cancer Treatment. Immuno. 2023;3(2):195–216. doi:10.3390/immuno3020013

22. Saraiva DP, Matias AT, Braga S, Jacinto A, Cabral MG. Establishment of a 3D Co-culture With MDA-MB-231 Breast Cancer Cell Line and Patient-Derived Immune Cells for Application in the Development of Immunotherapies. Front Oncol. 2020;10. doi:10.3389/fonc.2020.01543

23. Martinez-Lopez M, Póvoa V, Fior R. Generation of Zebrafish Larval Xenografts and Tumor Behavior Analysis. J Vis Exp. 2021;(172). doi:10.3791/62373-v

24. Fior R, Póvoa V, Mendes R V., et al. Single-cell functional and chemosensitive profiling of combinatorial colorectal therapy in zebrafish xenografts. Proc Natl Acad Sci. 2017;114(39). doi:10.1073/pnas.1618389114

25. Rasmussen A-M, Borelli G, Hoel HJ, et al. Ex vivo expansion protocol for human tumor specific T cells for adoptive T cell therapy. J Immunol Methods. 2010;355(1-2):52–60. doi:10.1016/j.jim.2010.02.004

26. Smith C, Økern G, Rehan S, et al. Ex vivo expansion of human T cells for adoptive immunotherapy using the novel Xeno-free CTS Immune Cell Serum Replacement. Clin Transl Immunol. 2015;4(1). doi:10.1038/cti.2014.31

27. Martino E, Misso G, Pastina P, et al. Immune-modulating effects of bevacizumab in metastatic non-small-cell lung cancer patients. Cell Death Discov. 2016;2(1):16025. doi:10.1038/cddiscovery.2016.25

28. Napoletano C, Ruscito I, Bellati F, et al. Bevacizumab-Based Chemotherapy Triggers Immunological Effects in Responding Multi-Treated Recurrent Ovarian Cancer Patients by Favoring the Recruitment of Effector T Cell Subsets. J Clin Med. 2019;8(3):380. doi:10.3390/jcm8030380

29. Arnold M, Morgan E, Rumgay H, et al. Current and future burden of breast cancer: Global statistics for 2020 and 2040. The Breast. 2022;66:15–23. doi:10.1016/j.breast.2022.08.010

30. Li X, Zhu Y, Yi J, Deng Y, Lei B, Ren H. Adoptive cell immunotherapy for breast cancer: harnessing the power of immune cells. J Leukoc Biol. Published online November 9, 2023. doi:10.1093/jleuko/qiad144

31. Dey A, Ghosh S, Jha S, et al. Recent advancement in breast cancer treatment using CAR T cell therapy:-A review. Adv Cancer Biol - Metastasis. 2023;7:100090. doi:10.1016/j.adcanc.2023.100090

32. Morad G, Helmink BA, Sharma P, Wargo JA. Hallmarks of response, resistance, and toxicity to immune checkpoint blockade. Cell. 2021;184(21):5309–5337. doi:10.1016/j.cell.2021.09.020

33. Cortes J, Rugo HS, Cescon DW, et al. Pembrolizumab plus Chemotherapy in Advanced Triple-Negative Breast Cancer. N Engl J Med. 2022;387(3):217–226. doi:10.1056/NEJMoa2202809

34. Loi S, Giobbie-Hurder A, Gombos A, et al. Pembrolizumab plus trastuzumab in trastuzumab-resistant, advanced, HER2-positive breast cancer (PANACEA): a single-arm, multicentre, phase 1b–2 trial. Lancet Oncol. 2019;20(3):371–382. doi:10.1016/S1470-2045(18)30812-X

35. Agostinetto E, Montemurro F, Puglisi F, et al. Immunotherapy for HER2-Positive Breast Cancer: Clinical Evidence and Future Perspectives. Cancers (Basel). 2022;14(9). doi:10.3390/cancers14092136

36. Osuna-Gómez R, Arqueros C, Galano C, et al. Effector Mechanisms of CD8+ HLA-DR+ T Cells in Breast Cancer Patients Who Respond to Neoadjuvant Chemotherapy. Cancers (Basel). 2021;13(24):6167. doi:10.3390/cancers13246167

37. Holling TM, Schooten E, van Den Elsen PJ. Function and regulation of MHC class II molecules in T-lymphocytes: of mice and men. Hum Immunol. 2004;65(4):282–290. doi:10.1016/j.humimm.2004.01.005

38. Gérard A, Khan O, Beemiller P, et al. Secondary T cell–T cell synaptic interactions drive the differentiation of protective CD8+ T cells. Nat Immunol. 2013;14(4):356–363. doi:10.1038/ni.2547

39. Moynihan KD, Kumar MP, Sultan H, et al. IL2 Targeted to CD8+ T Cells Promotes Robust Effector T-cell Responses and Potent Antitumor Immunity. Cancer Discov. 2024;14(7):1206–1225. doi:10.1158/2159-8290.CD-23-1266

40. Tam W. Identification and characterization of human BIC, a gene on chromosome 21 that encodes a noncoding RNA. Gene. 2001;274(1-2):157–167. doi:10.1016/S0378-1119(01)00612-6

41. Tam W, Ben-Yehuda D, Hayward WS. bic, a Novel Gene Activated by Proviral Insertions in Avian Leukosis Virus-Induced Lymphomas, Is Likely To Function through Its Noncoding RNA. Mol Cell Biol. 1997;17(3):1490–1502. doi:10.1128/MCB.17.3.1490

42. Lin H, Ni R, Li D, et al. LncRNA MIR155HG Overexpression Promotes Proliferation, Migration, and Chemoresistance in Gastric Cancer Cells. Int J Med Sci. 2023;20(7):933–942. doi:10.7150/ijms.82216

43. Peng L, Chen Z, Chen Y, Wang X, Tang N. MIR155HG is a prognostic biomarker and associated with immune infiltration and immune checkpoint molecules expression in multiple cancers. Cancer Med. 2019;8(17):7161–7173. doi:10.1002/cam4.2583

44. Xia F, Wang Y, Xue M, et al. LncRNA KCNQ1OT1: Molecular mechanisms and pathogenic roles in human diseases. Genes Dis. 2022;9(6):1556–1565. doi:10.1016/j.gendis.2021.07.003

45. Heng Y, Zhu X, Wu Q, et al. High Expression of Tumor HLA-DR Predicts Better Prognosis and Response to Anti-PD-1 Therapy in Laryngeal Squamous Cell Carcinoma. Transl Oncol. 2023;33:101678. doi:10.1016/j.tranon.2023.101678

46. Kamphorst AO, Pillai RN, Yang S, et al. Proliferation of PD-1+ CD8 T cells in peripheral blood after PD-1–targeted therapy in lung cancer patients. Proc Natl Acad Sci. 2017;114(19):4993–4998. doi:10.1073/pnas.1705327114

47. Simon S, Labarriere N. PD-1 expression on tumor-specific T cells: Friend or foe for immunotherapy? Oncoimmunology. 2018;7(1):e1364828. doi:10.1080/2162402X.2017.1364828

48. Mattick JS, Amaral PP, Carninci P, et al. Long non-coding RNAs: definitions, functions, challenges and recommendations. Nat Rev Mol Cell Biol. 2023;24(6):430–447. doi:10.1038/s41580-022-00566-8

49. Mattick JS. Non-coding RNAs: the architects of eukaryotic complexity. EMBO Rep. 2001;2(11):986–991. doi:10.1093/embo-reports/kve230

50. Cabili MN, Trapnell C, Goff L, et al. Integrative annotation of human large intergenic noncoding RNAs reveals global properties and specific subclasses. Genes Dev. 2011;25(18):1915–1927. doi:10.1101/gad.17446611

