## Supplementary material for "Enhancing HLA-DR in Cytotoxic T Lymphocytes is crucial for the development of efficient adoptive T cell Therapies for Breast Cancer": https://eur01.safelinks.protection.outlook.com/?url=https%3A%2F%2Fdrive.google.com%2Fdrive%2Ffolders%2F1GQdUyL_CFZbh8kN3N2J6MufQbc_j8r3z%3Fusp%3Ddrive

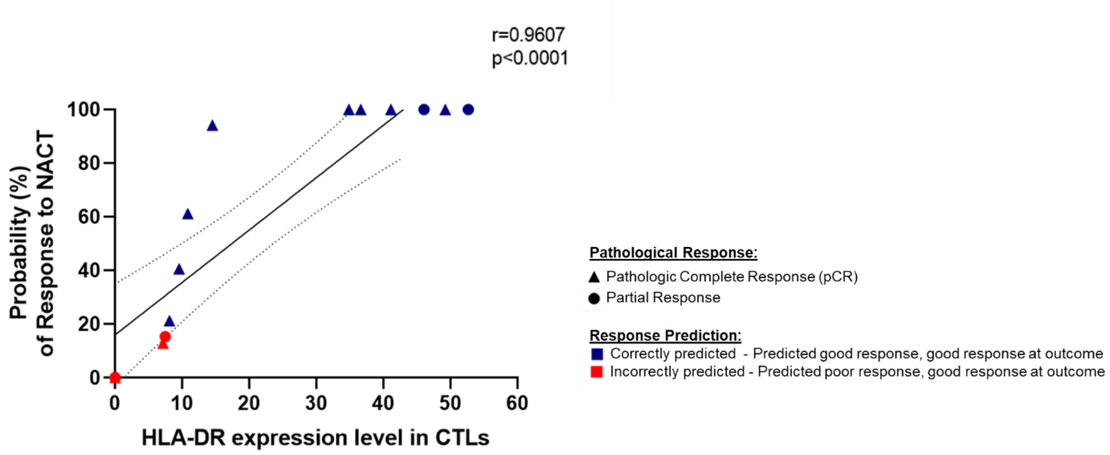


**Figure S1 – Validation of the predictive probability model of response to Neoadjuvant Chemotherapy (NACT) based on the HLA-DR level in Cytotoxic T Lymphocytes (CTLs).** Validation of our predictive probability model of BC patients' response to NACT, based on the HLA-DR level in infiltrating CTLs, assessed in patients' biopsies[12]. This validation was conducted in a cohort of 14 patients who underwent NACT. All patients responded to the treatment, with their responses categorized as either pathologic complete response (triangles) or partial response (circles). Our model accurately predicted the response to NACT for the majority of these patients (blue dots), with only four patients' responses not correctly predicted (red dots). This validation confirms the model's utility in predicting which BC patients will benefit from NACT, with a Spearman correlation of r=0.9607 (p<0.0001, n=14).

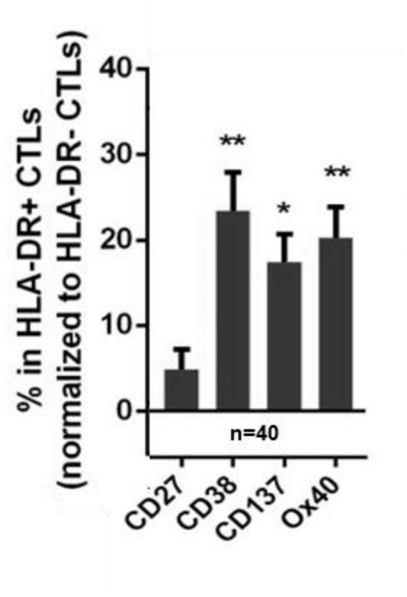


**Figure S2 - Immune profile of HLA-DR+ CTLs compared to HLA-DR- CTLs.** % of HLA-DR+CTLs that also express the stimulatory co-receptors CD27, CD38, CD137 and Ox40, related to the % of HLA-DR-CTLs that also express those markers. Results reveal that in general all the markers analyzed are more expressed in HLA-DR+ CTLs compared to HLA-DR- CTLs (n=40). Data are represented as ± SEM, $*p < 0.05$, $**p < 0.01$.

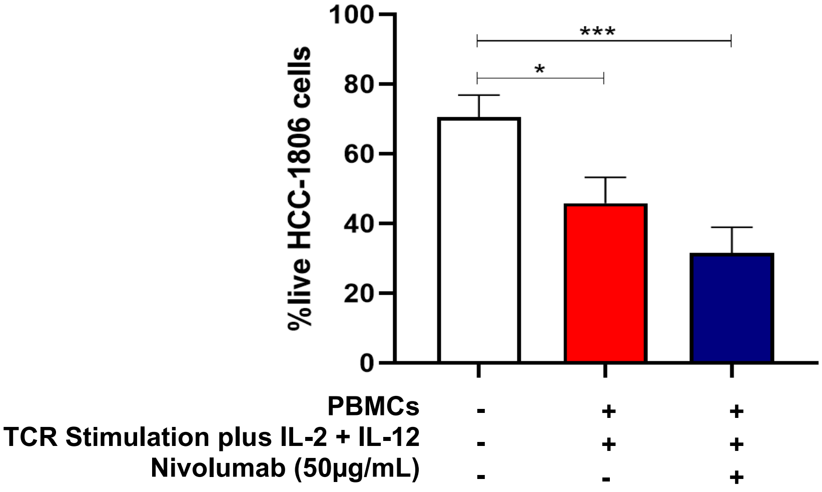


**Figure S3 – Anti-PD-1 treatment potentiates the anti-tumor capacity of cytotoxic T lymphocytes (CTLs) against the HCC-1806 Breast Cancer cell line.** The viability of the HCC-1806 BC cell line in the 3D co-culture alone (white bar, n=8), incubated in the presence of stimulated PBMCs (red bar, n=8, p=0.0486) and incubated in the presence of stimulated PBMCs plus the addition of the anti-PD-1, Nivolumab (blue bar, n=8, p<0.0001), demonstrating that cytotoxicity enhanced synergistically with the stimulation protocol previously established and the addition of Nivolumab to the co-culture. Data are represented as mean ± SD, **p < 0.05, ***p < 0.001*.

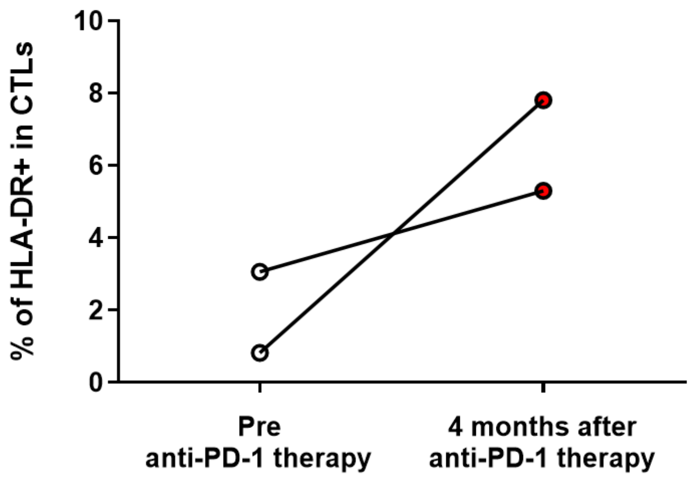


**Figure S4 – Breast Cancer (BC) patients submitted to anti-PD-1 therapy exhibited an elevated frequency of Cytotoxic T Lymphocytes (CTLs) expressing HLA-DR in their blood.** The BC patients undergoing anti-PD-1 therapy showed a higher percentage of CTLs expressing HLA-DR in their blood 4 months after starting this treatment. Each line represents one patient under anti-PD-1 therapy (n=2).

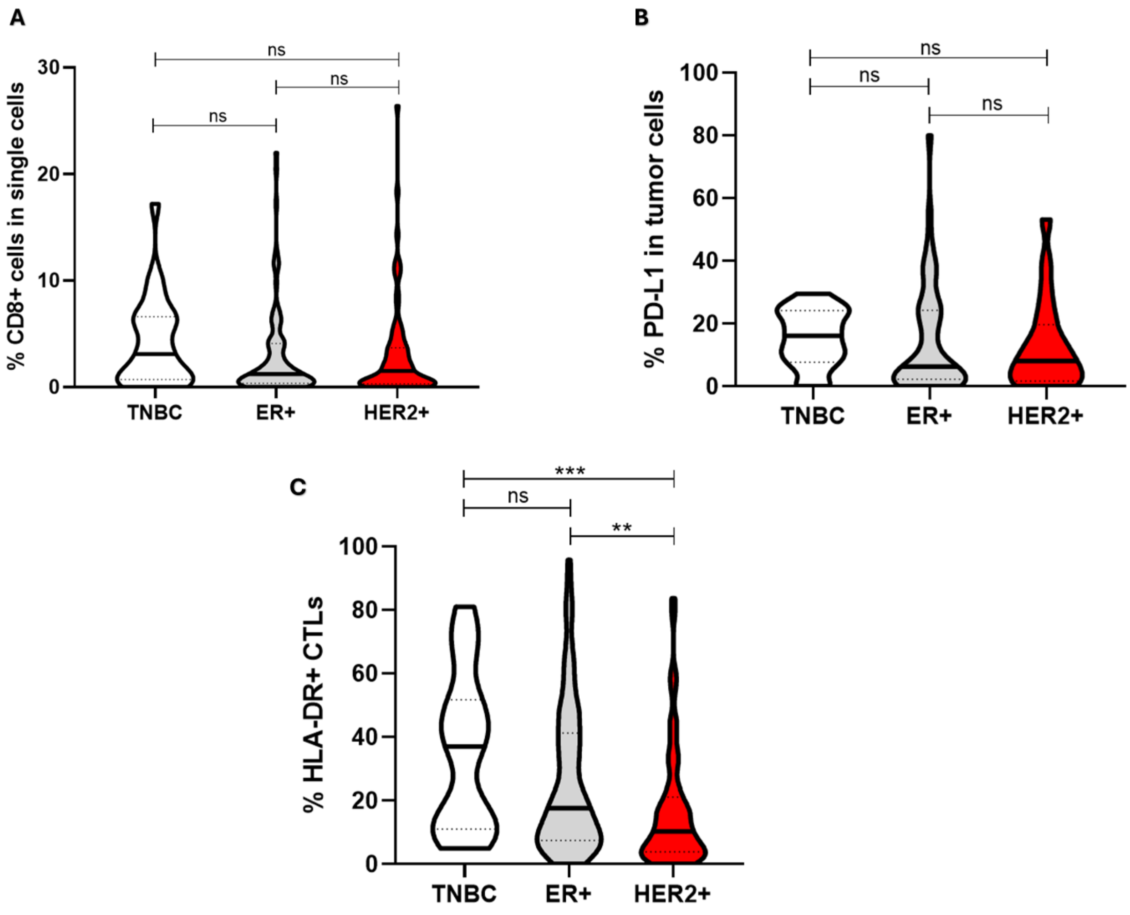


**Figure S5 – Phenotyping of patients' biopsies across the three main Breast Cancer subtypes.** Analysis of the immune profile by flow cytometry of biopsies from a cohort of 195 breast cancer (BC) patients. Only three relevant parameters were shown. (**A)** Percentage of infiltrating CTLs, showing no significant differences between the analyzed subtypes: TNBC (n=26), HER2+ (n=74), and Luminal A and B (ER+, n=95). **(B)** Percentage of tumor cells expressing PD-L1, showing consistency of this marker expression across the analyzed subtypes: TNBC (n=16), HER2+ (n=20) and ER+ (n=47). **(C)** Percentage of HLA-DR-expressing infiltrating CTLs, demonstrating a significant difference between this parameter in TNBC cases (n=25, p= 0.0003) in comparison with HER2+ (n=69) and ER+ (n=91). Data are represented as mean ± SD, ***p < 0.01, ***p < 0.001, ns= non-statistical*.

**Table S1 - Overview of participant characteristics enrolled in the study – biopsy and blood donors.** Median values for age and body mass index (BMI) are included, along with key tumor characteristics such as dimension and Ki67 levels (related to tumor proliferation rate). Additionally, other clinical data including breast cancer subtype, node status, and grade are summarized.

| | Patients | | Healthy Donors |
|---|---|---|---|
| | **Neoadjuvant Breast Cancer** | **Advanced Breast Cancer** | |
| **Number of subjects** | 195 | 14 | 32 |
| **Sample type** | Biopsies | Blood | Blood |
| **Age** | Median: 57<br>(Range: 29 - 90) | Median: 58<br>(range: 47 - 84) | range: 18–65 |
| **Body Mass Index (BMI)** | Median: 26.45<br>(range: 19.14 - 46.64) | Median: 26.06<br>(range: 18.37 - 35.56) | NA |
| **Subtype:**<br>• **ER+ (PR –/+)**<br>• **HER2+**<br>• **TNBC** | <br>48.72%<br>37.95%<br>13.33% | <br>71.42%<br>14.29%<br>14.29% | <br>NA<br>NA<br>NA |
| **Tumor dimension (mm)** | Median: 32.91<br>(range: 6 - 200) | Median: 57.92<br>(range: 2.5 - 160) | NA |
| **Ki67** | Median: 35<br>(range: 1 - 97) | Median: 40<br>(range: 19 - 90) | NA |
| **Axillary lymph node invasion status** | Negative – 62.57%<br>Positive – 37.43% | Negative – 50.00%<br>Positive – 50.00% | NA |
| **Grade** | G1 - 20.29%<br>G2 - 51.49%<br>G3 - 28.22% | G1 – 0.00%<br>G2 – 35.72%<br>G3 - 28.57%<br>Not known – 35.71% | NA |
| **Treatment:**<br>• **NACT**<br>• **Other chemotherapeutic treatments** | 100%<br>NA | 35.71%<br>64.29% | NA |

*NA: non-applicable.

**Table S2 –** Summary of the drugs tested, and the BC cell lines used in a small-size screening developed in the 3D platform established by our team[22], aiming to identify candidates that enhance HLA-DR expression in cytotoxic T lymphocytes (CTLs), and therefore their cytotoxic capacity against the BC cells. Each candidate entry corresponds to a specific agent assessed in the study. The "BC Cell Line" column specifies the particular BC cell line against which agents were tested, evaluating their promise and effectiveness. The “Characteristics of BC cell line” column recaps the representativeness of BC subtype and the expression of PD-L1 in the respective cell lines used in the study.

| **BC cell line** | **Characteristics of BC cell line[38]** | **Candidates:** | | | |
|---|---|---|---|---|---|
| | | **Nivolumab (Anti PD-1)** | **Bevacizumab (Anti VEGF)** | **Anti CD137** | **Anti CD134** |
| **MDA MB 231** | • Triple-negative BC<br>• High Expression of PD-L1 | ↑ | ↓ | ↓ | ↓ |
| **MCF-7** | • Luminal A (ER+ PR+)<br>• Low Expression of PD-L1 | ↓ | ↓ | ⦸ | ⦸ |
| **Hs578T** | • Triple-negative BC<br>• Middle Expression of PD-L1 | × | ⦸ | ⦸ | ⦸ |
| **BT-474** | • HER2+<br>• Low Expression of PD-L1 | × | ⦸ | ⦸ | ⦸ |
| **HCC 1806** | • Triple-negative BC<br>• High Expression of PD-L1 | ↑ | ⦸ | ⦸ | ⦸ |

↑ Candidate exhibits anti-tumor capabilities.
↓ Candidate fails to demonstrate anti-tumor properties.
⦸ Candidate's anti-tumor capabilities remain untested.
× Low spheroid viability or inadequate immune cell penetration into spheroids hampered the assessment of candidate anti-tumor efficacy.

**Table S3 –** List of genes significantly correlated with HLA-DR from BRCA_GSE150660 database

| Gene | Gene 2 | Cell Type | Correlation value |
|---|---|---|---|
| **HLA-DRA** | CD74 | CD8T | 0.778 |
| **HLA-DRA** | GZMH | CD8T | 0.714 |
| **HLA-DRA** | CCL4 | CD8T | 0.71 |
| **HLA-DRA** | CST7 | CD8T | 0.685 |
| **HLA-DRA** | CCR5 | CD8T | 0.646 |
| **HLA-DRA** | TMSB4X | CD8T | 0.643 |
| **HLA-DRA** | CCL5 | CD8T | 0.623 |
| **HLA-DRA** | EOMES | CD8T | 0.603 |
| **HLA-DRA** | GZMA | CD8T | 0.597 |
| **HLA-DRA** | HIGD2A | CD8T | 0.58 |
| **HLA-DRA** | CD81 | CD8T | 0.574 |
| **HLA-DRA** | SERF2 | CD8T | 0.567 |
| **HLA-DRA** | ITGB2 | CD8T | 0.554 |
| **HLA-DRA** | GZMK | CD8T | 0.549 |
| **HLA-DRA** | OAZ1 | CD8T | 0.549 |
| **HLA-DRA** | COTL1 | CD8T | 0.549 |
| **HLA-DRA** | CD8B | CD8T | 0.542 |
| **HLA-DRA** | NKG7 | CD8T | 0.53 |
| **HLA-DRA** | EIF1 | CD8T | 0.521 |
| **HLA-DRA** | CD8A | CD8T | 0.515 |
| **HLA-DRA** | PLEK | CD8T | 0.512 |

**Table S4 –** List of genes significantly correlated with HLA-DR from BRCA_GSE110686 database

| Gene | Gene 2 | Cell Type | Correlation value |
|---|---|---|---|
| **HLA-DRA** | CD74 | CD8T | 0.884 |
| **HLA-DRA** | CXCL13 | CD8T | 0.878 |
| **HLA-DRA** | LSP1 | CD8T | 0.872 |
| **HLA-DRA** | CCL3 | CD8T | 0.871 |
| **HLA-DRA** | GZMB | CD8T | 0.866 |
| **HLA-DRA** | HAVCR2 | CD8T | 0.864 |
| **HLA-DRA** | CCL4 | CD8T | 0.861 |
| **HLA-DRA** | TIGIT | CD8T | 0.859 |
| **HLA-DRA** | FABP5 | CD8T | 0.857 |
| **HLA-DRA** | DUSP4 | CD8T | 0.854 |
| **HLA-DRA** | PTMS | CD8T | 0.852 |
| **HLA-DRA** | GZMH | CD8T | 0.85 |
| **HLA-DRA** | LAG3 | CD8T | 0.839 |
| **HLA-DRA** | ANXA5 | CD8T | 0.835 |
| **HLA-DRA** | PRF1 | CD8T | 0.832 |
| **HLA-DRA** | ID2 | CD8T | 0.832 |
| **HLA-DRA** | GAPDH | CD8T | 0.829 |
| **HLA-DRA** | CXCR6 | CD8T | 0.816 |
| **HLA-DRA** | AC069363.1 | CD8T | 0.813 |
| **HLA-DRA** | VCAM1 | CD8T | 0.812 |
| **HLA-DRA** | FKBP1A | CD8T | 0.806 |
| **HLA-DRA** | GZMA | CD8T | 0.806 |
| **HLA-DRA** | PDCD1 | CD8T | 0.801 |
| **HLA-DRA** | PTTG1 | CD8T | 0.8 |
| **HLA-DRA** | SIRPG | CD8T | 0.799 |
| **HLA-DRA** | CD27 | CD8T | 0.798 |
| **HLA-DRA** | ACP5 | CD8T | 0.797 |
| **HLA-DRA** | COTL1 | CD8T | 0.796 |
| **HLA-DRA** | GNLY | CD8T | 0.795 |
| **HLA-DRA** | DYNLL1 | CD8T | 0.795 |
| **HLA-DRA** | PARK7 | CD8T | 0.794 |
| **HLA-DRA** | MIR155HG | CD8T | 0.792 |
| **HLA-DRA** | NKG7 | CD8T | 0.791 |
| **HLA-DRA** | YWHAH | CD8T | 0.789 |
| **HLA-DRA** | FASLG | CD8T | 0.788 |

| Gene | Gene 2 | Cell Type | Correlation value |
|---|---|---|---|
| **HLA-DRA** | RABAC1 | CD8T | 0.786 |
| **HLA-DRA** | IFNG | CD8T | 0.78 |
| **HLA-DRA** | APOBEC3G | CD8T | 0.778 |
| **HLA-DRA** | CLEC2B | CD8T | 0.771 |
| **HLA-DRA** | CCR5 | CD8T | 0.767 |
| **HLA-DRA** | CST7 | CD8T | 0.767 |
| **HLA-DRA** | CD8A | CD8T | 0.764 |
| **HLA-DRA** | RHOA | CD8T | 0.761 |
| **HLA-DRA** | CSF1 | CD8T | 0.759 |
| **HLA-DRA** | APOBEC3C | CD8T | 0.758 |
| **HLA-DRA** | RGS2 | CD8T | 0.757 |
| **HLA-DRA** | SH2D1A | CD8T | 0.757 |
| **HLA-DRA** | SIT1 | CD8T | 0.756 |
| **HLA-DRA** | SUMO2 | CD8T | 0.755 |
| **HLA-DRA** | PHLDA1 | CD8T | 0.753 |
| **HLA-DRA** | TNFRSF9 | CD8T | 0.749 |
| **HLA-DRA** | TRAC | CD8T | 0.749 |
| **HLA-DRA** | SYNGR2 | CD8T | 0.748 |
| **HLA-DRA** | ABI3 | CD8T | 0.748 |
| **HLA-DRA** | AC133644.2 | CD8T | 0.745 |
| **HLA-DRA** | IDH2 | CD8T | 0.741 |
| **HLA-DRA** | CLIC1 | CD8T | 0.739 |
| **HLA-DRA** | IFI27L2 | CD8T | 0.737 |
| **HLA-DRA** | ALOX5AP | CD8T | 0.73 |
| **HLA-DRA** | PSMB3 | CD8T | 0.729 |
| **HLA-DRA** | CCR1 | CD8T | 0.725 |
| **HLA-DRA** | ITM2A | CD8T | 0.719 |
| **HLA-DRA** | SCAMP2 | CD8T | 0.718 |
| **HLA-DRA** | KLRC1 | CD8T | 0.711 |
| **HLA-DRA** | CD2 | CD8T | 0.709 |
| **HLA-DRA** | PTPN6 | CD8T | 0.709 |
| **HLA-DRA** | LITAF | CD8T | 0.709 |
| **HLA-DRA** | ITGB1 | CD8T | 0.708 |
| **HLA-DRA** | CD8B | CD8T | 0.707 |
| **HLA-DRA** | BHLHE40 | CD8T | 0.706 |
| **HLA-DRA** | UCP2 | CD8T | 0.705 |
| **HLA-DRA** | IVNS1ABP | CD8T | 0.705 |

| Gene | Gene 2 | Cell Type | Correlation value |
|---|---|---|---|
| **HLA-DRA** | CHST12 | CD8T | 0.705 |
| **HLA-DRA** | DBI | CD8T | 0.704 |
| **HLA-DRA** | SRI | CD8T | 0.704 |
| **HLA-DRA** | MT1E | CD8T | 0.699 |
| **HLA-DRA** | LGALS1 | CD8T | 0.698 |
| **HLA-DRA** | LAYN | CD8T | 0.694 |
| **HLA-DRA** | COPZ1 | CD8T | 0.693 |
| **HLA-DRA** | SUB1 | CD8T | 0.69 |
| **HLA-DRA** | CAP1 | CD8T | 0.687 |
| **HLA-DRA** | SH2D2A | CD8T | 0.686 |
| **HLA-DRA** | EZR | CD8T | 0.682 |
| **HLA-DRA** | TXNDC17 | CD8T | 0.682 |
| **HLA-DRA** | CCND2 | CD8T | 0.678 |
| **HLA-DRA** | PGAM1 | CD8T | 0.678 |
| **HLA-DRA** | BATF | CD8T | 0.677 |
| **HLA-DRA** | IFI27 | CD8T | 0.675 |
| **HLA-DRA** | MYL6 | CD8T | 0.675 |
| **HLA-DRA** | HERPUD1 | CD8T | 0.674 |
| **HLA-DRA** | CD63 | CD8T | 0.673 |
| **HLA-DRA** | NRBP1 | CD8T | 0.672 |
| **HLA-DRA** | CAPZB | CD8T | 0.67 |
| **HLA-DRA** | PKM | CD8T | 0.669 |
| **HLA-DRA** | CKS2 | CD8T | 0.669 |
| **HLA-DRA** | CMC1 | CD8T | 0.668 |
| **HLA-DRA** | HNRNPLL | CD8T | 0.668 |
| **HLA-DRA** | TRBC1 | CD8T | 0.666 |
| **HLA-DRA** | CTSC | CD8T | 0.666 |
| **HLA-DRA** | HNRNPA3 | CD8T | 0.664 |
| **HLA-DRA** | SOX4 | CD8T | 0.664 |
| **HLA-DRA** | RHOC | CD8T | 0.663 |
| **HLA-DRA** | ID3 | CD8T | 0.663 |
| **HLA-DRA** | IGFLR1 | CD8T | 0.661 |
| **HLA-DRA** | CDK2AP2 | CD8T | 0.659 |
| **HLA-DRA** | CD3D | CD8T | 0.659 |
| **HLA-DRA** | TRAPPC1 | CD8T | 0.658 |
| **HLA-DRA** | ARPC2 | CD8T | 0.658 |
| **HLA-DRA** | SH3BGRL3 | CD8T | 0.657 |

| Gene | Gene 2 | Cell Type | Correlation value |
|---|---|---|---|
| **HLA-DRA** | CARD16 | CD8T | 0.656 |
| **HLA-DRA** | AIF1 | CD8T | 0.656 |
| **HLA-DRA** | BUB3 | CD8T | 0.655 |
| **HLA-DRA** | PRDX6 | CD8T | 0.652 |
| **HLA-DRA** | COMMD7 | CD8T | 0.649 |
| **HLA-DRA** | CD82 | CD8T | 0.646 |
| **HLA-DRA** | HMGB1 | CD8T | 0.641 |
| **HLA-DRA** | ZBED2 | CD8T | 0.64 |
| **HLA-DRA** | CD79A | CD8T | 0.64 |
| **HLA-DRA** | MT1X | CD8T | 0.638 |
| **HLA-DRA** | SRGN | CD8T | 0.637 |
| **HLA-DRA** | SERF2 | CD8T | 0.634 |
| **HLA-DRA** | CCL5 | CD8T | 0.631 |
| **HLA-DRA** | COX5A | CD8T | 0.631 |
| **HLA-DRA** | SRP14 | CD8T | 0.63 |
| **HLA-DRA** | RBX1 | CD8T | 0.629 |
| **HLA-DRA** | MCM5 | CD8T | 0.629 |
| **HLA-DRA** | SLA | CD8T | 0.626 |
| **HLA-DRA** | CD2BP2 | CD8T | 0.626 |
| **HLA-DRA** | HSPB1 | CD8T | 0.625 |
| **HLA-DRA** | ENSA | CD8T | 0.623 |
| **HLA-DRA** | SCP2 | CD8T | 0.623 |
| **HLA-DRA** | ADGRE5 | CD8T | 0.623 |
| **HLA-DRA** | GALM | CD8T | 0.622 |
| **HLA-DRA** | HMGN3 | CD8T | 0.621 |
| **HLA-DRA** | TPI1 | CD8T | 0.618 |
| **HLA-DRA** | HSBP1 | CD8T | 0.618 |
| **HLA-DRA** | HNRNPK | CD8T | 0.616 |
| **HLA-DRA** | NDUFC1 | CD8T | 0.615 |
| **HLA-DRA** | DOK2 | CD8T | 0.613 |
| **HLA-DRA** | BSG | CD8T | 0.613 |
| **HLA-DRA** | KLRC2 | CD8T | 0.612 |
| **HLA-DRA** | VCP | CD8T | 0.612 |
| **HLA-DRA** | TMED9 | CD8T | 0.609 |
| **HLA-DRA** | PSMB2 | CD8T | 0.608 |
| **HLA-DRA** | CTLA4 | CD8T | 0.608 |
| **HLA-DRA** | POMP | CD8T | 0.608 |

| Gene | Gene 2 | Cell Type | Correlation value |
|---|---|---|---|
| **HLA-DRA** | UBB | CD8T | 0.607 |
| **HLA-DRA** | CFL1 | CD8T | 0.607 |
| **HLA-DRA** | SOD1 | CD8T | 0.606 |
| **HLA-DRA** | NDUFB3 | CD8T | 0.606 |
| **HLA-DRA** | LCK | CD8T | 0.605 |
| **HLA-DRA** | CALM3 | CD8T | 0.603 |
| **HLA-DRA** | KIF20B | CD8T | 0.602 |
| **HLA-DRA** | CTSW | CD8T | 0.602 |
| **HLA-DRA** | KRT86 | CD8T | 0.601 |
| **HLA-DRA** | BAX | CD8T | 0.6 |
| **HLA-DRA** | CALM2 | CD8T | 0.599 |
| **HLA-DRA** | ARPP19 | CD8T | 0.599 |
| **HLA-DRA** | PSMC4 | CD8T | 0.598 |
| **HLA-DRA** | RGS1 | CD8T | 0.598 |
| **HLA-DRA** | BIN1 | CD8T | 0.597 |
| **HLA-DRA** | CLIC3 | CD8T | 0.595 |
| **HLA-DRA** | NUDT5 | CD8T | 0.594 |
| **HLA-DRA** | IL2RG | CD8T | 0.594 |
| **HLA-DRA** | CARHSP1 | CD8T | 0.593 |
| **HLA-DRA** | PCMT1 | CD8T | 0.593 |
| **HLA-DRA** | PYCARD | CD8T | 0.592 |
| **HLA-DRA** | TNFRSF1B | CD8T | 0.592 |
| **HLA-DRA** | H2AFZ | CD8T | 0.588 |
| **HLA-DRA** | OAZ1 | CD8T | 0.588 |
| **HLA-DRA** | MCL1 | CD8T | 0.587 |
| **HLA-DRA** | SAMSN1 | CD8T | 0.586 |
| **HLA-DRA** | NELFCD | CD8T | 0.585 |
| **HLA-DRA** | SNRPD1 | CD8T | 0.585 |
| **HLA-DRA** | PSMA7 | CD8T | 0.583 |
| **HLA-DRA** | PPDPF | CD8T | 0.583 |
| **HLA-DRA** | B3GNT2 | CD8T | 0.583 |
| **HLA-DRA** | PRDX5 | CD8T | 0.583 |
| **HLA-DRA** | CCDC28B | CD8T | 0.582 |
| **HLA-DRA** | NUSAP1 | CD8T | 0.582 |
| **HLA-DRA** | SURF4 | CD8T | 0.581 |
| **HLA-DRA** | AP2S1 | CD8T | 0.58 |
| **HLA-DRA** | TMEM165 | CD8T | 0.58 |

| Gene | Gene 2 | Cell Type | Correlation value |
|---|---|---|---|
| **HLA-DRA** | AIP | CD8T | 0.578 |
| **HLA-DRA** | SKA2 | CD8T | 0.577 |
| **HLA-DRA** | ATP6V1E1 | CD8T | 0.576 |
| **HLA-DRA** | PSMA1 | CD8T | 0.575 |
| **HLA-DRA** | UBE2A | CD8T | 0.573 |
| **HLA-DRA** | EOMES | CD8T | 0.573 |
| **HLA-DRA** | PDIA3 | CD8T | 0.572 |
| **HLA-DRA** | TANK | CD8T | 0.571 |
| **HLA-DRA** | CD247 | CD8T | 0.57 |
| **HLA-DRA** | COX8A | CD8T | 0.57 |
| **HLA-DRA** | DNAJC8 | CD8T | 0.569 |
| **HLA-DRA** | ACTB | CD8T | 0.567 |
| **HLA-DRA** | CLEC2D | CD8T | 0.565 |
| **HLA-DRA** | UBE2L3 | CD8T | 0.564 |
| **HLA-DRA** | YWHAQ | CD8T | 0.561 |
| **HLA-DRA** | FERMT3 | CD8T | 0.56 |
| **HLA-DRA** | YWHAE | CD8T | 0.558 |
| **HLA-DRA** | SLBP | CD8T | 0.558 |
| **HLA-DRA** | EID1 | CD8T | 0.558 |
| **HLA-DRA** | CORO1A | CD8T | 0.557 |
| **HLA-DRA** | CD3E | CD8T | 0.556 |
| **HLA-DRA** | PRR13 | CD8T | 0.555 |
| **HLA-DRA** | RNF167 | CD8T | 0.554 |
| **HLA-DRA** | PFN1 | CD8T | 0.554 |
| **HLA-DRA** | TMCO1 | CD8T | 0.554 |
| **HLA-DRA** | BRK1 | CD8T | 0.554 |
| **HLA-DRA** | HSPB11 | CD8T | 0.554 |
| **HLA-DRA** | NAP1L4 | CD8T | 0.553 |
| **HLA-DRA** | SEC11A | CD8T | 0.552 |
| **HLA-DRA** | LARP7 | CD8T | 0.551 |
| **HLA-DRA** | THYN1 | CD8T | 0.551 |
| **HLA-DRA** | PDIA6 | CD8T | 0.551 |
| **HLA-DRA** | HM13 | CD8T | 0.55 |
| **HLA-DRA** | ARPC3 | CD8T | 0.55 |
| **HLA-DRA** | PMF1 | CD8T | 0.55 |
| **HLA-DRA** | DNPH1 | CD8T | 0.548 |
| **HLA-DRA** | CMTM3 | CD8T | 0.548 |

| Gene | Gene 2 | Cell Type | Correlation value |
|---|---|---|---|
| **HLA-DRA** | SRSF9 | CD8T | 0.547 |
| **HLA-DRA** | PSMB9 | CD8T | 0.547 |
| **HLA-DRA** | ITM2C | CD8T | 0.547 |
| **HLA-DRA** | RAB11B | CD8T | 0.546 |
| **HLA-DRA** | CTSD | CD8T | 0.546 |
| **HLA-DRA** | ARPC1B | CD8T | 0.545 |
| **HLA-DRA** | PPP4C | CD8T | 0.544 |
| **HLA-DRA** | UBL5 | CD8T | 0.544 |
| **HLA-DRA** | PRKAR1A | CD8T | 0.543 |
| **HLA-DRA** | LINC01480 | CD8T | 0.543 |
| **HLA-DRA** | LBH | CD8T | 0.542 |
| **HLA-DRA** | MRPS16 | CD8T | 0.541 |
| **HLA-DRA** | HMGN1 | CD8T | 0.541 |
| **HLA-DRA** | LAMTOR5 | CD8T | 0.54 |
| **HLA-DRA** | BLOC1S1 | CD8T | 0.539 |
| **HLA-DRA** | KXD1 | CD8T | 0.538 |
| **HLA-DRA** | GSTP1 | CD8T | 0.538 |
| **HLA-DRA** | TMEM179B | CD8T | 0.538 |
| **HLA-DRA** | AKR1B1 | CD8T | 0.538 |
| **HLA-DRA** | HSD17B10 | CD8T | 0.537 |
| **HLA-DRA** | SAP18 | CD8T | 0.537 |
| **HLA-DRA** | SLC1A5 | CD8T | 0.536 |
| **HLA-DRA** | PEF1 | CD8T | 0.536 |
| **HLA-DRA** | MT1F | CD8T | 0.534 |
| **HLA-DRA** | RHBDD2 | CD8T | 0.531 |
| **HLA-DRA** | SERPINB9 | CD8T | 0.53 |
| **HLA-DRA** | ACTG1 | CD8T | 0.53 |
| **HLA-DRA** | NDUFV2 | CD8T | 0.529 |
| **HLA-DRA** | ARHGEF1 | CD8T | 0.528 |
| **HLA-DRA** | HMGN2 | CD8T | 0.528 |
| **HLA-DRA** | MRPS34 | CD8T | 0.527 |
| **HLA-DRA** | EIF4E | CD8T | 0.527 |
| **HLA-DRA** | CYB5B | CD8T | 0.527 |
| **HLA-DRA** | PSMD4 | CD8T | 0.526 |
| **HLA-DRA** | ASNA1 | CD8T | 0.526 |
| **HLA-DRA** | CYCS | CD8T | 0.526 |
| **HLA-DRA** | PSMB1 | CD8T | 0.523 |

| Gene | Gene 2 | Cell Type | Correlation value |
|---|---|---|---|
| **HLA-DRA** | COX6A1 | CD8T | 0.522 |
| **HLA-DRA** | NDUFA13 | CD8T | 0.522 |
| **HLA-DRA** | SHKBP1 | CD8T | 0.522 |
| **HLA-DRA** | PTPN7 | CD8T | 0.522 |
| **HLA-DRA** | RASAL3 | CD8T | 0.521 |
| **HLA-DRA** | B2M | CD8T | 0.521 |
| **HLA-DRA** | AP2M1 | CD8T | 0.521 |
| **HLA-DRA** | TXN | CD8T | 0.521 |
| **HLA-DRA** | ARL6IP1 | CD8T | 0.52 |
| **HLA-DRA** | RNF181 | CD8T | 0.52 |
| **HLA-DRA** | PSMD8 | CD8T | 0.519 |
| **HLA-DRA** | FKBP8 | CD8T | 0.519 |
| **HLA-DRA** | SMC4 | CD8T | 0.519 |
| **HLA-DRA** | COMMD3 | CD8T | 0.518 |
| **HLA-DRA** | CALR | CD8T | 0.517 |
| **HLA-DRA** | PSMD7 | CD8T | 0.517 |
| **HLA-DRA** | COPE | CD8T | 0.516 |
| **HLA-DRA** | SNU13 | CD8T | 0.516 |
| **HLA-DRA** | TMX1 | CD8T | 0.516 |
| **HLA-DRA** | FDPS | CD8T | 0.514 |
| **HLA-DRA** | SEC61G | CD8T | 0.513 |
| **HLA-DRA** | PLEKHJ1 | CD8T | 0.513 |
| **HLA-DRA** | ITGB2 | CD8T | 0.513 |
| **HLA-DRA** | ITGB7 | CD8T | 0.513 |
| **HLA-DRA** | TMEM59 | CD8T | 0.513 |
| **HLA-DRA** | PHPT1 | CD8T | 0.513 |
| **HLA-DRA** | HNRNPC | CD8T | 0.513 |
| **HLA-DRA** | CAPZA1 | CD8T | 0.512 |
| **HLA-DRA** | TSPO | CD8T | 0.512 |
| **HLA-DRA** | CYBA | CD8T | 0.511 |

**Table S5 –** List of genes significantly correlated with HLA-DR from BRCA_GSE114727_10X database

| Gene | Gene 2 | Cell Type | Correlation value |
|---|---|---|---|
| **HLA-DRA** | GZMH | CD8T | 0.864 |
| **HLA-DRA** | CTSC | CD8T | 0.84 |
| **HLA-DRA** | ANXA5 | CD8T | 0.828 |
| **HLA-DRA** | FABP5 | CD8T | 0.82 |
| **HLA-DRA** | APOBEC3G | CD8T | 0.819 |
| **HLA-DRA** | SUB1 | CD8T | 0.817 |
| **HLA-DRA** | IDH2 | CD8T | 0.81 |
| **HLA-DRA** | LSP1 | CD8T | 0.808 |
| **HLA-DRA** | APOBEC3C | CD8T | 0.806 |
| **HLA-DRA** | PTTG1 | CD8T | 0.803 |
| **HLA-DRA** | CAP1 | CD8T | 0.794 |
| **HLA-DRA** | COTL1 | CD8T | 0.788 |
| **HLA-DRA** | PSMB9 | CD8T | 0.78 |
| **HLA-DRA** | LAG3 | CD8T | 0.777 |
| **HLA-DRA** | PTMS | CD8T | 0.773 |
| **HLA-DRA** | ARPC2 | CD8T | 0.765 |
| **HLA-DRA** | CAPZB | CD8T | 0.765 |
| **HLA-DRA** | PPP1CA | CD8T | 0.763 |
| **HLA-DRA** | CXCR6 | CD8T | 0.761 |
| **HLA-DRA** | CD74 | CD8T | 0.755 |
| **HLA-DRA** | EVL | CD8T | 0.753 |
| **HLA-DRA** | POMP | CD8T | 0.752 |
| **HLA-DRA** | ALOX5AP | CD8T | 0.751 |
| **HLA-DRA** | RHOA | CD8T | 0.75 |
| **HLA-DRA** | CLIC1 | CD8T | 0.744 |
| **HLA-DRA** | PDCD1 | CD8T | 0.744 |
| **HLA-DRA** | CARD16 | CD8T | 0.743 |
| **HLA-DRA** | CTSD | CD8T | 0.739 |
| **HLA-DRA** | IFNG | CD8T | 0.737 |
| **HLA-DRA** | TPM3 | CD8T | 0.737 |
| **HLA-DRA** | PRF1 | CD8T | 0.736 |
| **HLA-DRA** | HMGB1 | CD8T | 0.734 |
| **HLA-DRA** | COX5A | CD8T | 0.732 |
| **HLA-DRA** | set/07 | CD8T | 0.732 |
| **HLA-DRA** | LCP1 | CD8T | 0.73 |

| Gene | Gene 2 | Cell Type | Correlation value |
|---|---|---|---|
| **HLA-DRA** | GZMA | CD8T | 0.728 |
| **HLA-DRA** | PYCARD | CD8T | 0.726 |
| **HLA-DRA** | CD2 | CD8T | 0.725 |
| **HLA-DRA** | DUSP4 | CD8T | 0.725 |
| **HLA-DRA** | RBCK1 | CD8T | 0.722 |
| **HLA-DRA** | ZYX | CD8T | 0.722 |
| **HLA-DRA** | LCK | CD8T | 0.721 |
| **HLA-DRA** | GAPDH | CD8T | 0.721 |
| **HLA-DRA** | PSTPIP1 | CD8T | 0.72 |
| **HLA-DRA** | ANXA6 | CD8T | 0.718 |
| **HLA-DRA** | CCL3 | CD8T | 0.718 |
| **HLA-DRA** | PSME2 | CD8T | 0.717 |
| **HLA-DRA** | UBE2L3 | CD8T | 0.717 |
| **HLA-DRA** | ACTB | CD8T | 0.716 |
| **HLA-DRA** | RAB1B | CD8T | 0.714 |
| **HLA-DRA** | PARK7 | CD8T | 0.714 |
| **HLA-DRA** | CCR5 | CD8T | 0.714 |
| **HLA-DRA** | REEP5 | CD8T | 0.712 |
| **HLA-DRA** | IFI16 | CD8T | 0.71 |
| **HLA-DRA** | WDR1 | CD8T | 0.708 |
| **HLA-DRA** | MAP4K1 | CD8T | 0.707 |
| **HLA-DRA** | ARPC3 | CD8T | 0.706 |
| **HLA-DRA** | MSC | CD8T | 0.706 |
| **HLA-DRA** | CFL1 | CD8T | 0.705 |
| **HLA-DRA** | RABAC1 | CD8T | 0.704 |
| **HLA-DRA** | DBI | CD8T | 0.703 |
| **HLA-DRA** | BATF | CD8T | 0.702 |
| **HLA-DRA** | DYNLRB1 | CD8T | 0.702 |
| **HLA-DRA** | PSMB3 | CD8T | 0.702 |
| **HLA-DRA** | SHISA5 | CD8T | 0.702 |
| **HLA-DRA** | CDK2AP2 | CD8T | 0.702 |
| **HLA-DRA** | YWHAB | CD8T | 0.701 |
| **HLA-DRA** | MSN | CD8T | 0.7 |
| **HLA-DRA** | UBE2L6 | CD8T | 0.7 |
| **HLA-DRA** | GNG5 | CD8T | 0.7 |
| **HLA-DRA** | IFI27L2 | CD8T | 0.699 |
| **HLA-DRA** | BTG3 | CD8T | 0.699 |

| Gene | Gene 2 | Cell Type | Correlation value |
|---|---|---|---|
| **HLA-DRA** | ADGRE5 | CD8T | 0.698 |
| **HLA-DRA** | HSPB11 | CD8T | 0.697 |
| **HLA-DRA** | PDIA6 | CD8T | 0.697 |
| **HLA-DRA** | CD3D | CD8T | 0.695 |
| **HLA-DRA** | GZMB | CD8T | 0.694 |
| **HLA-DRA** | ITGAE | CD8T | 0.694 |
| **HLA-DRA** | FERMT3 | CD8T | 0.691 |
| **HLA-DRA** | LY6E | CD8T | 0.69 |
| **HLA-DRA** | COPE | CD8T | 0.69 |
| **HLA-DRA** | GNAS | CD8T | 0.689 |
| **HLA-DRA** | CALR | CD8T | 0.689 |
| **HLA-DRA** | MYO1G | CD8T | 0.688 |
| **HLA-DRA** | B2M | CD8T | 0.687 |
| **HLA-DRA** | WAS | CD8T | 0.686 |
| **HLA-DRA** | ENSA | CD8T | 0.686 |
| **HLA-DRA** | PSMB8 | CD8T | 0.686 |
| **HLA-DRA** | PSMB10 | CD8T | 0.685 |
| **HLA-DRA** | ACTR3 | CD8T | 0.685 |
| **HLA-DRA** | AKIRIN2 | CD8T | 0.685 |
| **HLA-DRA** | PSMD8 | CD8T | 0.684 |
| **HLA-DRA** | EZR | CD8T | 0.683 |
| **HLA-DRA** | JAML | CD8T | 0.683 |
| **HLA-DRA** | MIR155HG | CD8T | 0.683 |
| **HLA-DRA** | SH3BP1 | CD8T | 0.683 |
| **HLA-DRA** | PPIB | CD8T | 0.679 |
| **HLA-DRA** | LCP2 | CD8T | 0.679 |
| **HLA-DRA** | ITM2A | CD8T | 0.678 |
| **HLA-DRA** | CORO1A | CD8T | 0.678 |
| **HLA-DRA** | ANXA2 | CD8T | 0.678 |
| **HLA-DRA** | CSK | CD8T | 0.677 |
| **HLA-DRA** | DYNLL1 | CD8T | 0.677 |
| **HLA-DRA** | MYH9 | CD8T | 0.676 |
| **HLA-DRA** | BIN1 | CD8T | 0.676 |
| **HLA-DRA** | SEC11A | CD8T | 0.676 |
| **HLA-DRA** | GPI | CD8T | 0.675 |
| **HLA-DRA** | RALY | CD8T | 0.675 |
| **HLA-DRA** | MYL6 | CD8T | 0.674 |

| Gene | Gene 2 | Cell Type | Correlation value |
|---|---|---|---|
| **HLA-DRA** | COPZ1 | CD8T | 0.674 |
| **HLA-DRA** | TBCB | CD8T | 0.672 |
| **HLA-DRA** | CHST12 | CD8T | 0.672 |
| **HLA-DRA** | CYBA | CD8T | 0.671 |
| **HLA-DRA** | RAC1 | CD8T | 0.671 |
| **HLA-DRA** | TAP1 | CD8T | 0.67 |
| **HLA-DRA** | PKM | CD8T | 0.669 |
| **HLA-DRA** | PSMA7 | CD8T | 0.669 |
| **HLA-DRA** | PIN1 | CD8T | 0.669 |
| **HLA-DRA** | COMMD7 | CD8T | 0.669 |
| **HLA-DRA** | TWF2 | CD8T | 0.667 |
| **HLA-DRA** | ASNA1 | CD8T | 0.666 |
| **HLA-DRA** | HNRNPA3 | CD8T | 0.666 |
| **HLA-DRA** | PSMA6 | CD8T | 0.666 |
| **HLA-DRA** | MRPL28 | CD8T | 0.666 |
| **HLA-DRA** | TXN | CD8T | 0.665 |
| **HLA-DRA** | RBX1 | CD8T | 0.665 |
| **HLA-DRA** | IL2RG | CD8T | 0.665 |
| **HLA-DRA** | PPP4C | CD8T | 0.664 |
| **HLA-DRA** | CASP4 | CD8T | 0.664 |
| **HLA-DRA** | CCL4 | CD8T | 0.664 |
| **HLA-DRA** | YWHAZ | CD8T | 0.662 |
| **HLA-DRA** | UBE2A | CD8T | 0.662 |
| **HLA-DRA** | BSG | CD8T | 0.661 |
| **HLA-DRA** | APBB1IP | CD8T | 0.66 |
| **HLA-DRA** | YWHAQ | CD8T | 0.659 |
| **HLA-DRA** | TNFRSF1B | CD8T | 0.658 |
| **HLA-DRA** | RAB11B | CD8T | 0.657 |
| **HLA-DRA** | UBE2D3 | CD8T | 0.657 |
| **HLA-DRA** | ARPC5L | CD8T | 0.657 |
| **HLA-DRA** | PSMB1 | CD8T | 0.656 |
| **HLA-DRA** | PLEKHF1 | CD8T | 0.656 |
| **HLA-DRA** | MDH2 | CD8T | 0.656 |
| **HLA-DRA** | TLN1 | CD8T | 0.656 |
| **HLA-DRA** | HCLS1 | CD8T | 0.656 |
| **HLA-DRA** | CD53 | CD8T | 0.653 |
| **HLA-DRA** | CD3G | CD8T | 0.653 |

| Gene | Gene 2 | Cell Type | Correlation value |
|---|---|---|---|
| **HLA-DRA** | CDCA7 | CD8T | 0.652 |
| **HLA-DRA** | PSMB6 | CD8T | 0.652 |
| **HLA-DRA** | TXNDC17 | CD8T | 0.651 |
| **HLA-DRA** | PPM1G | CD8T | 0.651 |
| **HLA-DRA** | PDZD11 | CD8T | 0.651 |
| **HLA-DRA** | DEK | CD8T | 0.651 |
| **HLA-DRA** | ACP5 | CD8T | 0.65 |
| **HLA-DRA** | SMC4 | CD8T | 0.649 |
| **HLA-DRA** | PSMB2 | CD8T | 0.649 |
| **HLA-DRA** | ARF1 | CD8T | 0.649 |
| **HLA-DRA** | MT2A | CD8T | 0.649 |
| **HLA-DRA** | SRGN | CD8T | 0.648 |
| **HLA-DRA** | PGAM1 | CD8T | 0.645 |
| **HLA-DRA** | NKG7 | CD8T | 0.645 |
| **HLA-DRA** | GYG1 | CD8T | 0.644 |
| **HLA-DRA** | ABI3 | CD8T | 0.644 |
| **HLA-DRA** | PPM1M | CD8T | 0.643 |
| **HLA-DRA** | ARHGAP9 | CD8T | 0.643 |
| **HLA-DRA** | GBP5 | CD8T | 0.641 |
| **HLA-DRA** | ARHGDIA | CD8T | 0.641 |
| **HLA-DRA** | PSMA5 | CD8T | 0.641 |
| **HLA-DRA** | FASLG | CD8T | 0.641 |
| **HLA-DRA** | TRAPPC1 | CD8T | 0.639 |
| **HLA-DRA** | MT1F | CD8T | 0.639 |
| **HLA-DRA** | GNB2 | CD8T | 0.639 |
| **HLA-DRA** | ARF5 | CD8T | 0.639 |
| **HLA-DRA** | VAMP8 | CD8T | 0.638 |
| **HLA-DRA** | RNF167 | CD8T | 0.638 |
| **HLA-DRA** | CAPN2 | CD8T | 0.637 |
| **HLA-DRA** | AP2M1 | CD8T | 0.636 |
| **HLA-DRA** | CD63 | CD8T | 0.636 |
| **HLA-DRA** | CD99 | CD8T | 0.635 |
| **HLA-DRA** | PPP1R18 | CD8T | 0.634 |
| **HLA-DRA** | CALM3 | CD8T | 0.634 |
| **HLA-DRA** | PARP1 | CD8T | 0.634 |
| **HLA-DRA** | AC069363.1 | CD8T | 0.633 |
| **HLA-DRA** | TMEM120A | CD8T | 0.632 |

| Gene | Gene 2 | Cell Type | Correlation value |
|---|---|---|---|
| **HLA-DRA** | OTUB1 | CD8T | 0.632 |
| **HLA-DRA** | CLTB | CD8T | 0.632 |
| **HLA-DRA** | TMEM50A | CD8T | 0.632 |
| **HLA-DRA** | RAC2 | CD8T | 0.632 |
| **HLA-DRA** | MIEN1 | CD8T | 0.632 |
| **HLA-DRA** | NDUFS6 | CD8T | 0.631 |
| **HLA-DRA** | PDIA3 | CD8T | 0.631 |
| **HLA-DRA** | BLOC1S1 | CD8T | 0.63 |
| **HLA-DRA** | SSBP4 | CD8T | 0.629 |
| **HLA-DRA** | SUMO2 | CD8T | 0.629 |
| **HLA-DRA** | FKBP8 | CD8T | 0.629 |
| **HLA-DRA** | BUB3 | CD8T | 0.628 |
| **HLA-DRA** | PPP1R7 | CD8T | 0.627 |
| **HLA-DRA** | NDUFA6 | CD8T | 0.627 |
| **HLA-DRA** | RARRES3 | CD8T | 0.627 |
| **HLA-DRA** | RNF213 | CD8T | 0.626 |
| **HLA-DRA** | ZCRB1 | CD8T | 0.625 |
| **HLA-DRA** | CTSW | CD8T | 0.625 |
| **HLA-DRA** | TBC1D10C | CD8T | 0.624 |
| **HLA-DRA** | SLC7A5 | CD8T | 0.624 |
| **HLA-DRA** | CLEC2B | CD8T | 0.623 |
| **HLA-DRA** | IL32 | CD8T | 0.623 |
| **HLA-DRA** | EWSR1 | CD8T | 0.622 |
| **HLA-DRA** | GTF3C6 | CD8T | 0.622 |
| **HLA-DRA** | VDAC1 | CD8T | 0.619 |
| **HLA-DRA** | AP2S1 | CD8T | 0.619 |
| **HLA-DRA** | RPN1 | CD8T | 0.619 |
| **HLA-DRA** | SRI | CD8T | 0.617 |
| **HLA-DRA** | ARPP19 | CD8T | 0.616 |
| **HLA-DRA** | TMEM59 | CD8T | 0.616 |
| **HLA-DRA** | NDUFB3 | CD8T | 0.616 |
| **HLA-DRA** | SH3BGRL | CD8T | 0.616 |
| **HLA-DRA** | ACTG1 | CD8T | 0.615 |
| **HLA-DRA** | F2R | CD8T | 0.613 |
| **HLA-DRA** | RNF181 | CD8T | 0.612 |
| **HLA-DRA** | MFSD10 | CD8T | 0.611 |
| **HLA-DRA** | SPCS2 | CD8T | 0.61 |

| Gene | Gene 2 | Cell Type | Correlation value |
|---|---|---|---|
| **HLA-DRA** | PRDX1 | CD8T | 0.61 |
| **HLA-DRA** | SLC9A3R1 | CD8T | 0.609 |
| **HLA-DRA** | SURF4 | CD8T | 0.609 |
| **HLA-DRA** | UBE2N | CD8T | 0.609 |
| **HLA-DRA** | ITGB7 | CD8T | 0.609 |
| **HLA-DRA** | GLIPR1 | CD8T | 0.608 |
| **HLA-DRA** | ITGB2 | CD8T | 0.607 |
| **HLA-DRA** | SH3BGRL3 | CD8T | 0.607 |
| **HLA-DRA** | RGS1 | CD8T | 0.606 |
| **HLA-DRA** | CD52 | CD8T | 0.606 |
| **HLA-DRA** | RNF187 | CD8T | 0.606 |
| **HLA-DRA** | NUDT21 | CD8T | 0.605 |
| **HLA-DRA** | SP100 | CD8T | 0.605 |
| **HLA-DRA** | CALCOCO2 | CD8T | 0.605 |
| **HLA-DRA** | BRK1 | CD8T | 0.605 |
| **HLA-DRA** | RER1 | CD8T | 0.604 |
| **HLA-DRA** | CCDC167 | CD8T | 0.604 |
| **HLA-DRA** | FKBP1A | CD8T | 0.604 |
| **HLA-DRA** | EID1 | CD8T | 0.604 |
| **HLA-DRA** | ZBP1 | CD8T | 0.604 |
| **HLA-DRA** | LAMTOR5 | CD8T | 0.604 |
| **HLA-DRA** | SCAMP2 | CD8T | 0.603 |
| **HLA-DRA** | ORMDL2 | CD8T | 0.603 |
| **HLA-DRA** | DCTN3 | CD8T | 0.603 |
| **HLA-DRA** | ZNHIT1 | CD8T | 0.603 |
| **HLA-DRA** | AIP | CD8T | 0.603 |
| **HLA-DRA** | HSBP1 | CD8T | 0.602 |
| **HLA-DRA** | BUD31 | CD8T | 0.602 |
| **HLA-DRA** | GSDMD | CD8T | 0.602 |
| **HLA-DRA** | DRAP1 | CD8T | 0.602 |
| **HLA-DRA** | UBC | CD8T | 0.601 |
| **HLA-DRA** | SASH3 | CD8T | 0.601 |
| **HLA-DRA** | TPI1 | CD8T | 0.601 |
| **HLA-DRA** | HNRNPK | CD8T | 0.601 |
| **HLA-DRA** | UBB | CD8T | 0.601 |
| **HLA-DRA** | PSMC4 | CD8T | 0.601 |
| **HLA-DRA** | BST2 | CD8T | 0.601 |

| Gene | Gene 2 | Cell Type | Correlation value |
|---|---|---|---|
| **HLA-DRA** | MYL12B | CD8T | 0.6 |
| **HLA-DRA** | MDH1 | CD8T | 0.6 |
| **HLA-DRA** | CHMP2A | CD8T | 0.6 |
| **HLA-DRA** | CPNE7 | CD8T | 0.598 |
| **HLA-DRA** | RNF7 | CD8T | 0.597 |
| **HLA-DRA** | PPP2R1A | CD8T | 0.597 |
| **HLA-DRA** | SLA | CD8T | 0.596 |
| **HLA-DRA** | TBCD | CD8T | 0.595 |
| **HLA-DRA** | ARL6IP5 | CD8T | 0.595 |
| **HLA-DRA** | MAGOH | CD8T | 0.595 |
| **HLA-DRA** | SEC11C | CD8T | 0.594 |
| **HLA-DRA** | VASP | CD8T | 0.594 |
| **HLA-DRA** | MEA1 | CD8T | 0.594 |
| **HLA-DRA** | ADRM1 | CD8T | 0.594 |
| **HLA-DRA** | DCTN2 | CD8T | 0.593 |
| **HLA-DRA** | ANP32A | CD8T | 0.593 |
| **HLA-DRA** | SLC1A5 | CD8T | 0.593 |
| **HLA-DRA** | GNAI2 | CD8T | 0.593 |
| **HLA-DRA** | IAH1 | CD8T | 0.593 |
| **HLA-DRA** | LPXN | CD8T | 0.592 |
| **HLA-DRA** | SSNA1 | CD8T | 0.592 |
| **HLA-DRA** | GUK1 | CD8T | 0.592 |
| **HLA-DRA** | COX6A1 | CD8T | 0.592 |
| **HLA-DRA** | SERPINB1 | CD8T | 0.592 |
| **HLA-DRA** | GSTP1 | CD8T | 0.592 |
| **HLA-DRA** | PSMA4 | CD8T | 0.592 |
| **HLA-DRA** | UBE2I | CD8T | 0.591 |
| **HLA-DRA** | AURKAIP1 | CD8T | 0.591 |
| **HLA-DRA** | KIF5B | CD8T | 0.591 |
| **HLA-DRA** | NELFCD | CD8T | 0.591 |
| **HLA-DRA** | JOSD2 | CD8T | 0.59 |
| **HLA-DRA** | CHMP1A | CD8T | 0.59 |
| **HLA-DRA** | TSPO | CD8T | 0.59 |
| **HLA-DRA** | ANAPC11 | CD8T | 0.59 |
| **HLA-DRA** | SHKBP1 | CD8T | 0.59 |
| **HLA-DRA** | PSMC3 | CD8T | 0.59 |
| **HLA-DRA** | UCP2 | CD8T | 0.589 |

| Gene | Gene 2 | Cell Type | Correlation value |
|---|---|---|---|
| **HLA-DRA** | C11orf58 | CD8T | 0.589 |
| **HLA-DRA** | TMEM258 | CD8T | 0.589 |
| **HLA-DRA** | AP1S1 | CD8T | 0.589 |
| **HLA-DRA** | TNFRSF9 | CD8T | 0.588 |
| **HLA-DRA** | SRSF9 | CD8T | 0.588 |
| **HLA-DRA** | DR1 | CD8T | 0.588 |
| **HLA-DRA** | TYMP | CD8T | 0.587 |
| **HLA-DRA** | COMMD8 | CD8T | 0.587 |
| **HLA-DRA** | RASAL3 | CD8T | 0.586 |
| **HLA-DRA** | PSMD4 | CD8T | 0.586 |
| **HLA-DRA** | NDUFS7 | CD8T | 0.586 |
| **HLA-DRA** | PSME1 | CD8T | 0.586 |
| **HLA-DRA** | GBP2 | CD8T | 0.585 |
| **HLA-DRA** | OPTN | CD8T | 0.585 |
| **HLA-DRA** | SCAND1 | CD8T | 0.585 |
| **HLA-DRA** | POLD4 | CD8T | 0.584 |
| **HLA-DRA** | CCL5 | CD8T | 0.584 |
| **HLA-DRA** | ATP6V0E1 | CD8T | 0.584 |
| **HLA-DRA** | COX7A2 | CD8T | 0.584 |
| **HLA-DRA** | COX7B | CD8T | 0.583 |
| **HLA-DRA** | MPG | CD8T | 0.583 |
| **HLA-DRA** | CXCL13 | CD8T | 0.583 |
| **HLA-DRA** | MRPS36 | CD8T | 0.583 |
| **HLA-DRA** | RBM42 | CD8T | 0.583 |
| **HLA-DRA** | SNRPB | CD8T | 0.583 |
| **HLA-DRA** | GLUD1 | CD8T | 0.582 |
| **HLA-DRA** | CD2BP2 | CD8T | 0.582 |
| **HLA-DRA** | SNRPB2 | CD8T | 0.582 |
| **HLA-DRA** | PRR13 | CD8T | 0.582 |
| **HLA-DRA** | BAX | CD8T | 0.581 |
| **HLA-DRA** | VAMP5 | CD8T | 0.581 |
| **HLA-DRA** | CCL4L2 | CD8T | 0.58 |
| **HLA-DRA** | TSTA3 | CD8T | 0.58 |
| **HLA-DRA** | SIT1 | CD8T | 0.58 |
| **HLA-DRA** | TMEM179B | CD8T | 0.579 |
| **HLA-DRA** | FIBP | CD8T | 0.579 |
| **HLA-DRA** | TIGIT | CD8T | 0.579 |

| Gene | Gene 2 | Cell Type | Correlation value |
|---|---|---|---|
| **HLA-DRA** | NDUFA1 | CD8T | 0.578 |
| **HLA-DRA** | SNRPG | CD8T | 0.576 |
| **HLA-DRA** | PUF60 | CD8T | 0.575 |
| **HLA-DRA** | WDR83OS | CD8T | 0.575 |
| **HLA-DRA** | NDUFB10 | CD8T | 0.575 |
| **HLA-DRA** | PGK1 | CD8T | 0.574 |
| **HLA-DRA** | SUMO1 | CD8T | 0.574 |
| **HLA-DRA** | TALDO1 | CD8T | 0.574 |
| **HLA-DRA** | LAMTOR2 | CD8T | 0.574 |
| **HLA-DRA** | SNX6 | CD8T | 0.574 |
| **HLA-DRA** | GOLGA7 | CD8T | 0.573 |
| **HLA-DRA** | COX8A | CD8T | 0.573 |
| **HLA-DRA** | POLR2G | CD8T | 0.573 |
| **HLA-DRA** | LGALS1 | CD8T | 0.572 |
| **HLA-DRA** | SELPLG | CD8T | 0.572 |
| **HLA-DRA** | TIMM8B | CD8T | 0.572 |
| **HLA-DRA** | VPS28 | CD8T | 0.572 |
| **HLA-DRA** | CASP1 | CD8T | 0.571 |
| **HLA-DRA** | NAA38 | CD8T | 0.571 |
| **HLA-DRA** | SEC61B | CD8T | 0.571 |
| **HLA-DRA** | ITGB1BP1 | CD8T | 0.57 |
| **HLA-DRA** | CMTM3 | CD8T | 0.57 |
| **HLA-DRA** | PFN1 | CD8T | 0.57 |
| **HLA-DRA** | TCIRG1 | CD8T | 0.569 |
| **HLA-DRA** | MIIP | CD8T | 0.569 |
| **HLA-DRA** | UBL5 | CD8T | 0.569 |
| **HLA-DRA** | PET100 | CD8T | 0.568 |
| **HLA-DRA** | IL2RB | CD8T | 0.567 |
| **HLA-DRA** | YWHAH | CD8T | 0.567 |
| **HLA-DRA** | SF3B6 | CD8T | 0.567 |
| **HLA-DRA** | TMEM14A | CD8T | 0.566 |
| **HLA-DRA** | XRCC5 | CD8T | 0.566 |
| **HLA-DRA** | CTLA4 | CD8T | 0.566 |
| **HLA-DRA** | C4orf48 | CD8T | 0.566 |
| **HLA-DRA** | CALM2 | CD8T | 0.566 |
| **HLA-DRA** | ANXA11 | CD8T | 0.564 |
| **HLA-DRA** | PDCD6 | CD8T | 0.564 |

| Gene | Gene 2 | Cell Type | Correlation value |
|---|---|---|---|
| **HLA-DRA** | GABARAPL1 | CD8T | 0.564 |
| **HLA-DRA** | TADA3 | CD8T | 0.563 |
| **HLA-DRA** | TPM4 | CD8T | 0.563 |
| **HLA-DRA** | LSM2 | CD8T | 0.562 |
| **HLA-DRA** | ILK | CD8T | 0.562 |
| **HLA-DRA** | TRAC | CD8T | 0.562 |
| **HLA-DRA** | HCST | CD8T | 0.562 |
| **HLA-DRA** | ARRDC1 | CD8T | 0.562 |
| **HLA-DRA** | TMX4 | CD8T | 0.562 |
| **HLA-DRA** | HMGN2 | CD8T | 0.561 |
| **HLA-DRA** | CDC42 | CD8T | 0.56 |
| **HLA-DRA** | CKS2 | CD8T | 0.56 |
| **HLA-DRA** | SEC61G | CD8T | 0.56 |
| **HLA-DRA** | HSD17B10 | CD8T | 0.56 |
| **HLA-DRA** | MOB1A | CD8T | 0.559 |
| **HLA-DRA** | CST7 | CD8T | 0.559 |
| **HLA-DRA** | OASL | CD8T | 0.558 |
| **HLA-DRA** | LSM8 | CD8T | 0.558 |
| **HLA-DRA** | SAMD9L | CD8T | 0.557 |
| **HLA-DRA** | MPC2 | CD8T | 0.556 |
| **HLA-DRA** | CD8A | CD8T | 0.556 |
| **HLA-DRA** | ROMO1 | CD8T | 0.556 |
| **HLA-DRA** | MAD2L2 | CD8T | 0.556 |
| **HLA-DRA** | PAIP2 | CD8T | 0.555 |
| **HLA-DRA** | RAB7A | CD8T | 0.555 |
| **HLA-DRA** | IFI35 | CD8T | 0.555 |
| **HLA-DRA** | COX6B1 | CD8T | 0.555 |
| **HLA-DRA** | MAP1LC3B | CD8T | 0.555 |
| **HLA-DRA** | ISG15 | CD8T | 0.554 |
| **HLA-DRA** | SAP18 | CD8T | 0.554 |
| **HLA-DRA** | ITGB1 | CD8T | 0.554 |
| **HLA-DRA** | CFLAR | CD8T | 0.553 |
| **HLA-DRA** | ARHGAP30 | CD8T | 0.553 |
| **HLA-DRA** | SF3B2 | CD8T | 0.553 |
| **HLA-DRA** | DUSP23 | CD8T | 0.552 |
| **HLA-DRA** | NABP2 | CD8T | 0.552 |
| **HLA-DRA** | SDF2L1 | CD8T | 0.552 |

| Gene | Gene 2 | Cell Type | Correlation value |
|---|---|---|---|
| **HLA-DRA** | NDUFAB1 | CD8T | 0.551 |
| **HLA-DRA** | SH2D2A | CD8T | 0.551 |
| **HLA-DRA** | RAP1B | CD8T | 0.551 |
| **HLA-DRA** | CAPZA1 | CD8T | 0.55 |
| **HLA-DRA** | CD3E | CD8T | 0.55 |
| **HLA-DRA** | APOBEC3H | CD8T | 0.55 |
| **HLA-DRA** | PTPN7 | CD8T | 0.549 |
| **HLA-DRA** | EIF6 | CD8T | 0.549 |
| **HLA-DRA** | LIME1 | CD8T | 0.549 |
| **HLA-DRA** | NDUFS3 | CD8T | 0.549 |
| **HLA-DRA** | TRIM22 | CD8T | 0.548 |
| **HLA-DRA** | AC133644.2 | CD8T | 0.548 |
| **HLA-DRA** | WIPF1 | CD8T | 0.547 |
| **HLA-DRA** | PSMC1 | CD8T | 0.546 |
| **HLA-DRA** | COX17 | CD8T | 0.546 |
| **HLA-DRA** | NUDT5 | CD8T | 0.545 |
| **HLA-DRA** | SLTM | CD8T | 0.545 |
| **HLA-DRA** | SLC25A5 | CD8T | 0.545 |
| **HLA-DRA** | CHCHD2 | CD8T | 0.545 |
| **HLA-DRA** | NDUFB7 | CD8T | 0.544 |
| **HLA-DRA** | ENO1 | CD8T | 0.544 |
| **HLA-DRA** | S100A6 | CD8T | 0.544 |
| **HLA-DRA** | GSTK1 | CD8T | 0.543 |
| **HLA-DRA** | PHPT1 | CD8T | 0.543 |
| **HLA-DRA** | PRDX5 | CD8T | 0.543 |
| **HLA-DRA** | CCNDBP1 | CD8T | 0.543 |
| **HLA-DRA** | ARPC5 | CD8T | 0.543 |
| **HLA-DRA** | LDHA | CD8T | 0.542 |
| **HLA-DRA** | KDELR2 | CD8T | 0.542 |
| **HLA-DRA** | KRTCAP2 | CD8T | 0.542 |
| **HLA-DRA** | NDUFS8 | CD8T | 0.542 |
| **HLA-DRA** | ACTR2 | CD8T | 0.541 |
| **HLA-DRA** | NDUFA2 | CD8T | 0.541 |
| **HLA-DRA** | BTN3A2 | CD8T | 0.541 |
| **HLA-DRA** | C9orf16 | CD8T | 0.541 |
| **HLA-DRA** | BCL2L1 | CD8T | 0.541 |
| **HLA-DRA** | NDUFB2 | CD8T | 0.541 |

| Gene | Gene 2 | Cell Type | Correlation value |
|---|---|---|---|
| **HLA-DRA** | TCEA1 | CD8T | 0.541 |
| **HLA-DRA** | GSTO1 | CD8T | 0.541 |
| **HLA-DRA** | PLEKHJ1 | CD8T | 0.54 |
| **HLA-DRA** | TMX1 | CD8T | 0.539 |
| **HLA-DRA** | NDUFC1 | CD8T | 0.539 |
| **HLA-DRA** | ITPA | CD8T | 0.539 |
| **HLA-DRA** | RHOG | CD8T | 0.539 |
| **HLA-DRA** | PSMA3 | CD8T | 0.538 |
| **HLA-DRA** | RAD21 | CD8T | 0.538 |
| **HLA-DRA** | DBNL | CD8T | 0.538 |
| **HLA-DRA** | ATP6V1F | CD8T | 0.538 |
| **HLA-DRA** | UQCR11 | CD8T | 0.538 |
| **HLA-DRA** | COX5B | CD8T | 0.538 |
| **HLA-DRA** | HNRNPF | CD8T | 0.538 |
| **HLA-DRA** | ATP6V0B | CD8T | 0.537 |
| **HLA-DRA** | DAD1 | CD8T | 0.536 |
| **HLA-DRA** | ATP6V0C | CD8T | 0.536 |
| **HLA-DRA** | UBE2D2 | CD8T | 0.536 |
| **HLA-DRA** | CHCHD5 | CD8T | 0.535 |
| **HLA-DRA** | FMNL1 | CD8T | 0.535 |
| **HLA-DRA** | ECHS1 | CD8T | 0.535 |
| **HLA-DRA** | GABARAP | CD8T | 0.534 |
| **HLA-DRA** | SYNGR2 | CD8T | 0.533 |
| **HLA-DRA** | LGALS3 | CD8T | 0.533 |
| **HLA-DRA** | CARHSP1 | CD8T | 0.533 |
| **HLA-DRA** | THRAP3 | CD8T | 0.533 |
| **HLA-DRA** | P4HB | CD8T | 0.533 |
| **HLA-DRA** | PLP2 | CD8T | 0.533 |
| **HLA-DRA** | YIPF3 | CD8T | 0.533 |
| **HLA-DRA** | CCND2 | CD8T | 0.532 |
| **HLA-DRA** | SIRPG | CD8T | 0.532 |
| **HLA-DRA** | HERPUD1 | CD8T | 0.532 |
| **HLA-DRA** | DEF6 | CD8T | 0.532 |
| **HLA-DRA** | MRPL41 | CD8T | 0.532 |
| **HLA-DRA** | FLNA | CD8T | 0.532 |
| **HLA-DRA** | VIM | CD8T | 0.532 |
| **HLA-DRA** | PRDX6 | CD8T | 0.532 |

| Gene | Gene 2 | Cell Type | Correlation value |
|---|---|---|---|
| **HLA-DRA** | HNRNPLL | CD8T | 0.532 |
| **HLA-DRA** | EMC7 | CD8T | 0.532 |
| **HLA-DRA** | CISD3 | CD8T | 0.531 |
| **HLA-DRA** | H2AFZ | CD8T | 0.53 |
| **HLA-DRA** | CYB561D2 | CD8T | 0.53 |
| **HLA-DRA** | OAZ1 | CD8T | 0.529 |
| **HLA-DRA** | IDH3G | CD8T | 0.529 |
| **HLA-DRA** | H3F3B | CD8T | 0.529 |
| **HLA-DRA** | TMEM160 | CD8T | 0.528 |
| **HLA-DRA** | LAMTOR1 | CD8T | 0.528 |
| **HLA-DRA** | ELOVL1 | CD8T | 0.528 |
| **HLA-DRA** | ISG20 | CD8T | 0.528 |
| **HLA-DRA** | NUDT1 | CD8T | 0.528 |
| **HLA-DRA** | SNRPD3 | CD8T | 0.527 |
| **HLA-DRA** | TEX264 | CD8T | 0.527 |
| **HLA-DRA** | APOL6 | CD8T | 0.526 |
| **HLA-DRA** | CBX3 | CD8T | 0.526 |
| **HLA-DRA** | METTL9 | CD8T | 0.526 |
| **HLA-DRA** | FGFR1OP2 | CD8T | 0.526 |
| **HLA-DRA** | RHOF | CD8T | 0.526 |

**Table S6 –** List of genes significantly correlated with HLA-DR from BRCA_GSE161529 database

| Gene | Gene 2 | Cell Type | Correlation value |
|---|---|---|---|
| **HLA-DRA** | CD74 | CD8T | 0.961 |
| **HLA-DRA** | LAPTM5 | CD8T | 0.876 |
| **HLA-DRA** | CD53 | CD8T | 0.851 |
| **HLA-DRA** | EVI2B | CD8T | 0.832 |
| **HLA-DRA** | GPSM3 | CD8T | 0.823 |
| **HLA-DRA** | HCLS1 | CD8T | 0.813 |
| **HLA-DRA** | CTSS | CD8T | 0.808 |
| **HLA-DRA** | RGS1 | CD8T | 0.801 |
| **HLA-DRA** | NCF1 | CD8T | 0.796 |
| **HLA-DRA** | CYBB | CD8T | 0.787 |
| **HLA-DRA** | CD83 | CD8T | 0.786 |
| **HLA-DRA** | CD37 | CD8T | 0.786 |
| **HLA-DRA** | ITGB2 | CD8T | 0.783 |
| **HLA-DRA** | CTSH | CD8T | 0.78 |
| **HLA-DRA** | PLEK | CD8T | 0.778 |
| **HLA-DRA** | RNASE6 | CD8T | 0.777 |
| **HLA-DRA** | TYROBP | CD8T | 0.776 |
| **HLA-DRA** | AIF1 | CD8T | 0.773 |
| **HLA-DRA** | GMFG | CD8T | 0.772 |
| **HLA-DRA** | GPR183 | CD8T | 0.772 |
| **HLA-DRA** | C1orf162 | CD8T | 0.77 |
| **HLA-DRA** | FERMT3 | CD8T | 0.77 |
| **HLA-DRA** | LCP1 | CD8T | 0.768 |
| **HLA-DRA** | SPI1 | CD8T | 0.768 |
| **HLA-DRA** | LGALS9 | CD8T | 0.765 |
| **HLA-DRA** | HCST | CD8T | 0.761 |
| **HLA-DRA** | LST1 | CD8T | 0.761 |
| **HLA-DRA** | PTPN6 | CD8T | 0.76 |
| **HLA-DRA** | TAGAP | CD8T | 0.758 |
| **HLA-DRA** | CARD16 | CD8T | 0.756 |

| Gene | Gene 2 | Cell Type | Correlation value |
|---|---|---|---|
| **HLA-DRA** | ARHGDIB | CD8T | 0.755 |
| **HLA-DRA** | PTPRC | CD8T | 0.755 |
| **HLA-DRA** | FCER1G | CD8T | 0.75 |
| **HLA-DRA** | LYZ | CD8T | 0.749 |
| **HLA-DRA** | MS4A6A | CD8T | 0.747 |
| **HLA-DRA** | IL10RA | CD8T | 0.744 |
| **HLA-DRA** | CD84 | CD8T | 0.744 |
| **HLA-DRA** | MNDA | CD8T | 0.744 |
| **HLA-DRA** | SRGN | CD8T | 0.742 |
| **HLA-DRA** | CD48 | CD8T | 0.74 |
| **HLA-DRA** | LAT2 | CD8T | 0.74 |
| **HLA-DRA** | FYB1 | CD8T | 0.738 |
| **HLA-DRA** | LPXN | CD8T | 0.736 |
| **HLA-DRA** | LAIR1 | CD8T | 0.733 |
| **HLA-DRA** | FCGR2B | CD8T | 0.732 |
| **HLA-DRA** | C1QA | CD8T | 0.732 |
| **HLA-DRA** | GPR65 | CD8T | 0.732 |
| **HLA-DRA** | THEMIS2 | CD8T | 0.731 |
| **HLA-DRA** | LILRB1 | CD8T | 0.729 |
| **HLA-DRA** | RNASET2 | CD8T | 0.726 |
| **HLA-DRA** | PSMB9 | CD8T | 0.726 |
| **HLA-DRA** | C1QB | CD8T | 0.722 |
| **HLA-DRA** | CD4 | CD8T | 0.722 |
| **HLA-DRA** | LILRB4 | CD8T | 0.72 |
| **HLA-DRA** | IGSF6 | CD8T | 0.72 |
| **HLA-DRA** | ARRB2 | CD8T | 0.719 |
| **HLA-DRA** | LCP2 | CD8T | 0.715 |
| **HLA-DRA** | CASP1 | CD8T | 0.713 |
| **HLA-DRA** | C1QC | CD8T | 0.712 |
| **HLA-DRA** | CCL3 | CD8T | 0.71 |
| **HLA-DRA** | CYTH4 | CD8T | 0.71 |
| **HLA-DRA** | EVI2A | CD8T | 0.707 |

| Gene | Gene 2 | Cell Type | Correlation value |
|---|---|---|---|
| **HLA-DRA** | ABI3 | CD8T | 0.705 |
| **HLA-DRA** | TNFSF13B | CD8T | 0.705 |
| **HLA-DRA** | LYL1 | CD8T | 0.705 |
| **HLA-DRA** | CTSZ | CD8T | 0.703 |
| **HLA-DRA** | SLC15A3 | CD8T | 0.703 |
| **HLA-DRA** | CD68 | CD8T | 0.701 |
| **HLA-DRA** | CD14 | CD8T | 0.7 |
| **HLA-DRA** | CCL4 | CD8T | 0.699 |
| **HLA-DRA** | FCGR3A | CD8T | 0.698 |
| **HLA-DRA** | FCGR2A | CD8T | 0.698 |
| **HLA-DRA** | ALOX5AP | CD8T | 0.697 |
| **HLA-DRA** | GNAI2 | CD8T | 0.693 |
| **HLA-DRA** | CCR1 | CD8T | 0.693 |
| **HLA-DRA** | CORO1A | CD8T | 0.692 |
| **HLA-DRA** | RGS19 | CD8T | 0.692 |
| **HLA-DRA** | CXCR4 | CD8T | 0.691 |
| **HLA-DRA** | FCGR1A | CD8T | 0.69 |
| **HLA-DRA** | RASSF4 | CD8T | 0.688 |
| **HLA-DRA** | CSF1R | CD8T | 0.686 |
| **HLA-DRA** | SNX10 | CD8T | 0.686 |
| **HLA-DRA** | TPP1 | CD8T | 0.684 |
| **HLA-DRA** | CD300A | CD8T | 0.681 |
| **HLA-DRA** | MPEG1 | CD8T | 0.681 |
| **HLA-DRA** | FPR3 | CD8T | 0.68 |
| **HLA-DRA** | ACP5 | CD8T | 0.68 |
| **HLA-DRA** | DOK2 | CD8T | 0.68 |
| **HLA-DRA** | C3AR1 | CD8T | 0.678 |
| **HLA-DRA** | AP1S2 | CD8T | 0.678 |
| **HLA-DRA** | SAMHD1 | CD8T | 0.678 |
| **HLA-DRA** | CCL4L2 | CD8T | 0.678 |
| **HLA-DRA** | ITGAX | CD8T | 0.676 |
| **HLA-DRA** | LY96 | CD8T | 0.676 |

| Gene | Gene 2 | Cell Type | Correlation value |
|---|---|---|---|
| **HLA-DRA** | IL18 | CD8T | 0.675 |
| **HLA-DRA** | GIMAP4 | CD8T | 0.675 |
| **HLA-DRA** | NFKBID | CD8T | 0.674 |
| **HLA-DRA** | OLR1 | CD8T | 0.672 |
| **HLA-DRA** | BCL2A1 | CD8T | 0.67 |
| **HLA-DRA** | MPP1 | CD8T | 0.669 |
| **HLA-DRA** | TNFAIP8L2 | CD8T | 0.667 |
| **HLA-DRA** | CSF2RA | CD8T | 0.667 |
| **HLA-DRA** | SELPLG | CD8T | 0.662 |
| **HLA-DRA** | LRRC25 | CD8T | 0.662 |
| **HLA-DRA** | CLEC4A | CD8T | 0.66 |
| **HLA-DRA** | CYBA | CD8T | 0.659 |
| **HLA-DRA** | FAM49B | CD8T | 0.659 |
| **HLA-DRA** | GBP4 | CD8T | 0.657 |
| **HLA-DRA** | SAMD9L | CD8T | 0.657 |
| **HLA-DRA** | IL2RG | CD8T | 0.655 |
| **HLA-DRA** | RHOG | CD8T | 0.655 |
| **HLA-DRA** | KCTD12 | CD8T | 0.655 |
| **HLA-DRA** | FGR | CD8T | 0.653 |
| **HLA-DRA** | LILRB2 | CD8T | 0.653 |
| **HLA-DRA** | SIGLEC10 | CD8T | 0.652 |
| **HLA-DRA** | NCF2 | CD8T | 0.652 |
| **HLA-DRA** | APOC1 | CD8T | 0.651 |
| **HLA-DRA** | IFI30 | CD8T | 0.65 |
| **HLA-DRA** | UNC93B1 | CD8T | 0.65 |
| **HLA-DRA** | CSK | CD8T | 0.649 |
| **HLA-DRA** | IRF8 | CD8T | 0.648 |
| **HLA-DRA** | TREM2 | CD8T | 0.648 |
| **HLA-DRA** | FGL2 | CD8T | 0.646 |
| **HLA-DRA** | PARP14 | CD8T | 0.646 |
| **HLA-DRA** | SLAMF8 | CD8T | 0.644 |
| **HLA-DRA** | STAT1 | CD8T | 0.644 |

| Gene | Gene 2 | Cell Type | Correlation value |
|---|---|---|---|
| **HLA-DRA** | CNPY3 | CD8T | 0.641 |
| **HLA-DRA** | SLA | CD8T | 0.64 |
| **HLA-DRA** | MS4A4A | CD8T | 0.639 |
| **HLA-DRA** | CXorf21 | CD8T | 0.638 |
| **HLA-DRA** | CD52 | CD8T | 0.637 |
| **HLA-DRA** | IL4I1 | CD8T | 0.637 |
| **HLA-DRA** | SERPINB9 | CD8T | 0.633 |
| **HLA-DRA** | VSIG4 | CD8T | 0.632 |
| **HLA-DRA** | SP110 | CD8T | 0.632 |
| **HLA-DRA** | EFHD2 | CD8T | 0.632 |
| **HLA-DRA** | LIMD2 | CD8T | 0.631 |
| **HLA-DRA** | RNASE1 | CD8T | 0.629 |
| **HLA-DRA** | TGFB1 | CD8T | 0.629 |
| **HLA-DRA** | CYTIP | CD8T | 0.626 |
| **HLA-DRA** | CD163 | CD8T | 0.625 |
| **HLA-DRA** | ADAM8 | CD8T | 0.625 |
| **HLA-DRA** | GRN | CD8T | 0.624 |
| **HLA-DRA** | SH2B3 | CD8T | 0.623 |
| **HLA-DRA** | FCGR1B | CD8T | 0.622 |
| **HLA-DRA** | TCIRG1 | CD8T | 0.62 |
| **HLA-DRA** | CTSD | CD8T | 0.617 |
| **HLA-DRA** | SPP1 | CD8T | 0.617 |
| **HLA-DRA** | CALHM6 | CD8T | 0.616 |
| **HLA-DRA** | APOBR | CD8T | 0.616 |
| **HLA-DRA** | TNFRSF14 | CD8T | 0.615 |
| **HLA-DRA** | OSCAR | CD8T | 0.614 |
| **HLA-DRA** | IFI44L | CD8T | 0.614 |
| **HLA-DRA** | IGKC | CD8T | 0.614 |
| **HLA-DRA** | LAP3 | CD8T | 0.613 |
| **HLA-DRA** | MYO1G | CD8T | 0.613 |
| **HLA-DRA** | CCRL2 | CD8T | 0.612 |
| **HLA-DRA** | C5AR1 | CD8T | 0.612 |

| Gene | Gene 2 | Cell Type | Correlation value |
|---|---|---|---|
| **HLA-DRA** | STK4 | CD8T | 0.611 |
| **HLA-DRA** | ARHGAP9 | CD8T | 0.61 |
| **HLA-DRA** | XAF1 | CD8T | 0.608 |
| **HLA-DRA** | SGK1 | CD8T | 0.605 |
| **HLA-DRA** | LSP1 | CD8T | 0.603 |
| **HLA-DRA** | CSF2RB | CD8T | 0.601 |
| **HLA-DRA** | HHEX | CD8T | 0.6 |
| **HLA-DRA** | SIGLEC1 | CD8T | 0.597 |
| **HLA-DRA** | MOB1A | CD8T | 0.597 |
| **HLA-DRA** | ACTR2 | CD8T | 0.597 |
| **HLA-DRA** | GPR34 | CD8T | 0.596 |
| **HLA-DRA** | CLEC7A | CD8T | 0.596 |
| **HLA-DRA** | SAMSN1 | CD8T | 0.594 |
| **HLA-DRA** | TLR7 | CD8T | 0.592 |
| **HLA-DRA** | SLC16A3 | CD8T | 0.59 |
| **HLA-DRA** | IL1B | CD8T | 0.59 |
| **HLA-DRA** | PPP1R18 | CD8T | 0.589 |
| **HLA-DRA** | CXCL10 | CD8T | 0.587 |
| **HLA-DRA** | VSIR | CD8T | 0.587 |
| **HLA-DRA** | LILRB3 | CD8T | 0.586 |
| **HLA-DRA** | SLAMF7 | CD8T | 0.585 |
| **HLA-DRA** | CCL3L1 | CD8T | 0.583 |
| **HLA-DRA** | OTUD1 | CD8T | 0.583 |
| **HLA-DRA** | ADGRE5 | CD8T | 0.582 |
| **HLA-DRA** | PLA2G7 | CD8T | 0.582 |
| **HLA-DRA** | GBP5 | CD8T | 0.581 |
| **HLA-DRA** | CD40 | CD8T | 0.581 |
| **HLA-DRA** | TNFRSF1B | CD8T | 0.58 |
| **HLA-DRA** | GPR84 | CD8T | 0.579 |
| **HLA-DRA** | PTGER4 | CD8T | 0.578 |
| **HLA-DRA** | IL10 | CD8T | 0.576 |
| **HLA-DRA** | SMIM25 | CD8T | 0.576 |

| Gene | Gene 2 | Cell Type | Correlation value |
|---|---|---|---|
| **HLA-DRA** | CLEC4E | CD8T | 0.575 |
| **HLA-DRA** | PYCARD | CD8T | 0.574 |
| **HLA-DRA** | B2M | CD8T | 0.574 |
| **HLA-DRA** | IFI35 | CD8T | 0.572 |
| **HLA-DRA** | APOBEC3C | CD8T | 0.572 |
| **HLA-DRA** | set/06 | CD8T | 0.571 |
| **HLA-DRA** | RAC2 | CD8T | 0.57 |
| **HLA-DRA** | CTSC | CD8T | 0.57 |
| **HLA-DRA** | NR4A2 | CD8T | 0.567 |
| **HLA-DRA** | REL | CD8T | 0.567 |
| **HLA-DRA** | RNF213 | CD8T | 0.566 |
| **HLA-DRA** | OSM | CD8T | 0.565 |
| **HLA-DRA** | IL18BP | CD8T | 0.564 |
| **HLA-DRA** | STAB1 | CD8T | 0.563 |
| **HLA-DRA** | P2RY13 | CD8T | 0.563 |
| **HLA-DRA** | CCR5 | CD8T | 0.563 |
| **HLA-DRA** | GIMAP1 | CD8T | 0.562 |
| **HLA-DRA** | IFIT3 | CD8T | 0.56 |
| **HLA-DRA** | ADORA3 | CD8T | 0.56 |
| **HLA-DRA** | JAML | CD8T | 0.558 |
| **HLA-DRA** | RGS2 | CD8T | 0.558 |
| **HLA-DRA** | STK17B | CD8T | 0.558 |
| **HLA-DRA** | CAPG | CD8T | 0.557 |
| **HLA-DRA** | SLC37A2 | CD8T | 0.557 |
| **HLA-DRA** | RENBP | CD8T | 0.555 |
| **HLA-DRA** | MRC1 | CD8T | 0.554 |
| **HLA-DRA** | OAS1 | CD8T | 0.553 |
| **HLA-DRA** | VMO1 | CD8T | 0.552 |
| **HLA-DRA** | TMEM273 | CD8T | 0.551 |
| **HLA-DRA** | LINC01857 | CD8T | 0.55 |
| **HLA-DRA** | SUCNR1 | CD8T | 0.549 |
| **HLA-DRA** | GAPT | CD8T | 0.549 |

| Gene | Gene 2 | Cell Type | Correlation value |
|---|---|---|---|
| **HLA-DRA** | NPC2 | CD8T | 0.548 |
| **HLA-DRA** | ADAMDEC1 | CD8T | 0.546 |
| **HLA-DRA** | SLC11A1 | CD8T | 0.545 |
| **HLA-DRA** | LGMN | CD8T | 0.541 |
| **HLA-DRA** | RILPL2 | CD8T | 0.541 |
| **HLA-DRA** | POU2F2 | CD8T | 0.539 |
| **HLA-DRA** | GBP1 | CD8T | 0.538 |
| **HLA-DRA** | MX1 | CD8T | 0.535 |
| **HLA-DRA** | LMO2 | CD8T | 0.535 |
| **HLA-DRA** | SPN | CD8T | 0.535 |
| **HLA-DRA** | GAL3ST4 | CD8T | 0.534 |
| **HLA-DRA** | AKR1B1 | CD8T | 0.533 |
| **HLA-DRA** | IGLC2 | CD8T | 0.533 |
| **HLA-DRA** | TXNIP | CD8T | 0.533 |
| **HLA-DRA** | CD69 | CD8T | 0.533 |
| **HLA-DRA** | NLRP3 | CD8T | 0.532 |
| **HLA-DRA** | CXCL9 | CD8T | 0.53 |
| **HLA-DRA** | CTSB | CD8T | 0.53 |
| **HLA-DRA** | CLEC2B | CD8T | 0.53 |
| **HLA-DRA** | GLRX | CD8T | 0.528 |
| **HLA-DRA** | RNASE2 | CD8T | 0.528 |
| **HLA-DRA** | TGFBI | CD8T | 0.528 |
| **HLA-DRA** | ARPC1B | CD8T | 0.527 |
| **HLA-DRA** | HAMP | CD8T | 0.526 |
| **HLA-DRA** | PARP9 | CD8T | 0.525 |
| **HLA-DRA** | C2 | CD8T | 0.525 |
| **HLA-DRA** | IFIT2 | CD8T | 0.524 |
| **HLA-DRA** | IGLC3 | CD8T | 0.524 |
| **HLA-DRA** | SDS | CD8T | 0.523 |
| **HLA-DRA** | CSF3R | CD8T | 0.521 |
| **HLA-DRA** | UCP2 | CD8T | 0.52 |
| **HLA-DRA** | APOE | CD8T | 0.52 |

| Gene | Gene 2 | Cell Type | Correlation value |
|---|---|---|---|
| **HLA-DRA** | MIR155HG | CD8T | 0.52 |
| **HLA-DRA** | PPT1 | CD8T | 0.52 |
| **HLA-DRA** | COTL1 | CD8T | 0.519 |
| **HLA-DRA** | MMP9 | CD8T | 0.519 |
| **HLA-DRA** | ANKRD22 | CD8T | 0.518 |
| **HLA-DRA** | NR1H3 | CD8T | 0.517 |

**Table S7** – List of genes significantly correlated with HLA-DR from BRCA_EMTAB8107 database

| Gene | Gene 2 | Cell Type | Correlation value |
|---|---|---|---|
| **HLA-DRA** | CD74 | CD8T | 0.794 |
| **HLA-DRA** | LAG3 | CD8T | 0.763 |
| **HLA-DRA** | FASLG | CD8T | 0.742 |
| **HLA-DRA** | IFNG | CD8T | 0.727 |
| **HLA-DRA** | PTMS | CD8T | 0.717 |
| **HLA-DRA** | ALOX5AP | CD8T | 0.701 |
| **HLA-DRA** | APOBEC3C | CD8T | 0.695 |
| **HLA-DRA** | CCL3 | CD8T | 0.69 |
| **HLA-DRA** | GZMH | CD8T | 0.684 |
| **HLA-DRA** | FABP5 | CD8T | 0.679 |
| **HLA-DRA** | OASL | CD8T | 0.678 |
| **HLA-DRA** | AC069363.1 | CD8T | 0.677 |
| **HLA-DRA** | GZMB | CD8T | 0.673 |
| **HLA-DRA** | CXCR6 | CD8T | 0.665 |
| **HLA-DRA** | CTSC | CD8T | 0.661 |
| **HLA-DRA** | IDH2 | CD8T | 0.647 |
| **HLA-DRA** | MT2A | CD8T | 0.644 |
| **HLA-DRA** | GBP5 | CD8T | 0.643 |
| **HLA-DRA** | ANXA5 | CD8T | 0.643 |
| **HLA-DRA** | COTL1 | CD8T | 0.636 |
| **HLA-DRA** | PDCD1 | CD8T | 0.631 |
| **HLA-DRA** | PRF1 | CD8T | 0.629 |
| **HLA-DRA** | CCL4 | CD8T | 0.624 |
| **HLA-DRA** | PSMB9 | CD8T | 0.622 |
| **HLA-DRA** | APOBEC3G | CD8T | 0.622 |
| **HLA-DRA** | ARPC2 | CD8T | 0.618 |
| **HLA-DRA** | MT1E | CD8T | 0.613 |
| **HLA-DRA** | ID2 | CD8T | 0.608 |
| **HLA-DRA** | TAP1 | CD8T | 0.607 |
| **HLA-DRA** | MT1F | CD8T | 0.605 |
| **HLA-DRA** | JAML | CD8T | 0.604 |
| **HLA-DRA** | GAPDH | CD8T | 0.603 |
| **HLA-DRA** | TIGIT | CD8T | 0.603 |
| **HLA-DRA** | MIR155HG | CD8T | 0.598 |
| **HLA-DRA** | VCAM1 | CD8T | 0.595 |

| Gene | Gene 2 | Cell Type | Correlation value |
|---|---|---|---|
| **HLA-DRA** | CD2 | CD8T | 0.591 |
| **HLA-DRA** | LGALS1 | CD8T | 0.59 |
| **HLA-DRA** | PTPN7 | CD8T | 0.586 |
| **HLA-DRA** | GZMA | CD8T | 0.583 |
| **HLA-DRA** | KRT86 | CD8T | 0.58 |
| **HLA-DRA** | PLPP1 | CD8T | 0.578 |
| **HLA-DRA** | IFI16 | CD8T | 0.578 |
| **HLA-DRA** | LY6E | CD8T | 0.575 |
| **HLA-DRA** | IFI35 | CD8T | 0.572 |
| **HLA-DRA** | CCND2 | CD8T | 0.571 |
| **HLA-DRA** | PLSCR1 | CD8T | 0.567 |
| **HLA-DRA** | CCR5 | CD8T | 0.566 |
| **HLA-DRA** | ACP5 | CD8T | 0.565 |
| **HLA-DRA** | SUB1 | CD8T | 0.562 |
| **HLA-DRA** | CDKN2A | CD8T | 0.558 |
| **HLA-DRA** | IFI27L2 | CD8T | 0.556 |
| **HLA-DRA** | DUSP4 | CD8T | 0.555 |
| **HLA-DRA** | ABI3 | CD8T | 0.555 |
| **HLA-DRA** | SAMD9L | CD8T | 0.555 |
| **HLA-DRA** | CTSD | CD8T | 0.551 |
| **HLA-DRA** | TNFRSF9 | CD8T | 0.549 |
| **HLA-DRA** | RBCK1 | CD8T | 0.547 |
| **HLA-DRA** | CXCL13 | CD8T | 0.544 |
| **HLA-DRA** | ISG15 | CD8T | 0.543 |
| **HLA-DRA** | PLEKHF1 | CD8T | 0.542 |
| **HLA-DRA** | ADGRE5 | CD8T | 0.542 |
| **HLA-DRA** | ACTN4 | CD8T | 0.541 |
| **HLA-DRA** | IFITM3 | CD8T | 0.54 |
| **HLA-DRA** | CCL4L2 | CD8T | 0.537 |
| **HLA-DRA** | PTTG1 | CD8T | 0.537 |
| **HLA-DRA** | CAPZB | CD8T | 0.536 |
| **HLA-DRA** | UBE2L6 | CD8T | 0.536 |
| **HLA-DRA** | POMP | CD8T | 0.535 |
| **HLA-DRA** | set/07 | CD8T | 0.53 |
| **HLA-DRA** | RHOA | CD8T | 0.528 |
| **HLA-DRA** | LSP1 | CD8T | 0.528 |
| **HLA-DRA** | TYMP | CD8T | 0.526 |

| Gene | Gene 2 | Cell Type | Correlation value |
|---|---|---|---|
| **HLA-DRA** | ITM2A | CD8T | 0.526 |
| **HLA-DRA** | MTHFD2 | CD8T | 0.525 |
| **HLA-DRA** | CST7 | CD8T | 0.523 |
| **HLA-DRA** | GBP1 | CD8T | 0.521 |
| **HLA-DRA** | ANXA2 | CD8T | 0.521 |
| **HLA-DRA** | BST2 | CD8T | 0.521 |
| **HLA-DRA** | PARK7 | CD8T | 0.52 |
| **HLA-DRA** | LCP2 | CD8T | 0.52 |
| **HLA-DRA** | CAPZA1 | CD8T | 0.519 |
| **HLA-DRA** | SRGN | CD8T | 0.519 |
| **HLA-DRA** | CSF1 | CD8T | 0.519 |
| **HLA-DRA** | CAPN2 | CD8T | 0.517 |
| **HLA-DRA** | SHISA5 | CD8T | 0.515 |
| **HLA-DRA** | DRAP1 | CD8T | 0.515 |
| **HLA-DRA** | CLEC2D | CD8T | 0.514 |
| **HLA-DRA** | PDIA6 | CD8T | 0.513 |
| **HLA-DRA** | LGALS3 | CD8T | 0.512 |
| **HLA-DRA** | ACTB | CD8T | 0.511 |
| **HLA-DRA** | DYNLL1 | CD8T | 0.51 |

**Table S8 –** List of genes significantly correlated with PD1 from BRCA_GSE110686 database

| Gene | Gene 2 | Cell Type | Correlation value |
|---|---|---|---|
| **PDCD1** | TIGIT | CD8T | 0.889 |
| **PDCD1** | PHLDA1 | CD8T | 0.881 |
| **PDCD1** | CCL3 | CD8T | 0.879 |
| **PDCD1** | TNFRSF9 | CD8T | 0.865 |
| **PDCD1** | FABP5 | CD8T | 0.854 |
| **PDCD1** | GAPDH | CD8T | 0.845 |
| **PDCD1** | CXCL13 | CD8T | 0.843 |
| **PDCD1** | VCAM1 | CD8T | 0.838 |
| **PDCD1** | HAVCR2 | CD8T | 0.834 |
| **PDCD1** | BHLHE40 | CD8T | 0.83 |
| **PDCD1** | GZMB | CD8T | 0.822 |
| **PDCD1** | DUSP4 | CD8T | 0.821 |
| **PDCD1** | IFNG | CD8T | 0.82 |
| **PDCD1** | BATF | CD8T | 0.815 |
| **PDCD1** | RGS2 | CD8T | 0.813 |
| **PDCD1** | SYNGR2 | CD8T | 0.812 |
| **PDCD1** | YWHAH | CD8T | 0.809 |
| **PDCD1** | PKM | CD8T | 0.806 |
| **PDCD1** | PTMS | CD8T | 0.805 |
| **PDCD1** | LAG3 | CD8T | 0.805 |
| **PDCD1** | MIR155HG | CD8T | 0.799 |
| **PDCD1** | ANXA5 | CD8T | 0.798 |
| **PDCD1** | SH2D2A | CD8T | 0.797 |
| **PDCD1** | AC069363.1 | CD8T | 0.797 |
| **PDCD1** | ACP5 | CD8T | 0.795 |
| **PDCD1** | SIRPG | CD8T | 0.793 |
| **PDCD1** | FASLG | CD8T | 0.793 |
| **PDCD1** | ID2 | CD8T | 0.792 |
| **PDCD1** | FKBP1A | CD8T | 0.791 |
| **PDCD1** | CXCR6 | CD8T | 0.787 |
| **PDCD1** | AC133644.2 | CD8T | 0.786 |
| **PDCD1** | CCND2 | CD8T | 0.776 |
| **PDCD1** | RHOA | CD8T | 0.775 |
| **PDCD1** | LSP1 | CD8T | 0.773 |
| **PDCD1** | CD27 | CD8T | 0.771 |

| Gene | Gene 2 | Cell Type | Correlation value |
|---|---|---|---|
| **PDCD1** | ITM2A | CD8T | 0.767 |
| **PDCD1** | SIT1 | CD8T | 0.766 |
| **PDCD1** | CD82 | CD8T | 0.764 |
| **PDCD1** | PARK7 | CD8T | 0.761 |
| **PDCD1** | ID3 | CD8T | 0.76 |
| **PDCD1** | TNFRSF1B | CD8T | 0.75 |
| **PDCD1** | CCL4 | CD8T | 0.748 |
| **PDCD1** | HSPB1 | CD8T | 0.747 |
| **PDCD1** | DYNLL1 | CD8T | 0.746 |
| **PDCD1** | CD74 | CD8T | 0.738 |
| **PDCD1** | SCAMP2 | CD8T | 0.735 |
| **PDCD1** | PGAM1 | CD8T | 0.733 |
| **PDCD1** | CKS2 | CD8T | 0.73 |
| **PDCD1** | HSBP1 | CD8T | 0.727 |
| **PDCD1** | CTLA4 | CD8T | 0.726 |
| **PDCD1** | DNPH1 | CD8T | 0.723 |
| **PDCD1** | PRDX6 | CD8T | 0.721 |
| **PDCD1** | COPZ1 | CD8T | 0.721 |
| **PDCD1** | LITAF | CD8T | 0.72 |
| **PDCD1** | NR4A2 | CD8T | 0.717 |
| **PDCD1** | PRF1 | CD8T | 0.71 |
| **PDCD1** | KLRC1 | CD8T | 0.709 |
| **PDCD1** | GNLY | CD8T | 0.708 |
| **PDCD1** | SRI | CD8T | 0.706 |
| **PDCD1** | ALOX5AP | CD8T | 0.706 |
| **PDCD1** | SOX4 | CD8T | 0.705 |
| **PDCD1** | SNRPD1 | CD8T | 0.704 |
| **PDCD1** | ZBED2 | CD8T | 0.703 |
| **PDCD1** | NDUFA13 | CD8T | 0.698 |
| **PDCD1** | SLA | CD8T | 0.696 |
| **PDCD1** | SUMO2 | CD8T | 0.696 |
| **PDCD1** | CD63 | CD8T | 0.694 |
| **PDCD1** | CSF1 | CD8T | 0.693 |
| **PDCD1** | DBI | CD8T | 0.689 |
| **PDCD1** | TRAPPC1 | CD8T | 0.688 |
| **PDCD1** | CD2 | CD8T | 0.685 |
| **PDCD1** | LAYN | CD8T | 0.684 |

| Gene | Gene 2 | Cell Type | Correlation value |
|---|---|---|---|
| **PDCD1** | GALM | CD8T | 0.684 |
| **PDCD1** | SAMSN1 | CD8T | 0.681 |
| **PDCD1** | RABAC1 | CD8T | 0.68 |
| **PDCD1** | GZMH | CD8T | 0.68 |
| **PDCD1** | PTTG1 | CD8T | 0.679 |
| **PDCD1** | SOD1 | CD8T | 0.677 |
| **PDCD1** | VCP | CD8T | 0.676 |
| **PDCD1** | COX8A | CD8T | 0.673 |
| **PDCD1** | TRAC | CD8T | 0.673 |
| **PDCD1** | CDK2AP2 | CD8T | 0.671 |
| **PDCD1** | IFI27L2 | CD8T | 0.671 |
| **PDCD1** | CALM3 | CD8T | 0.669 |
| **PDCD1** | EZR | CD8T | 0.667 |
| **PDCD1** | PRDM1 | CD8T | 0.667 |
| **PDCD1** | CHST12 | CD8T | 0.667 |
| **PDCD1** | CLEC2D | CD8T | 0.66 |
| **PDCD1** | PTPN6 | CD8T | 0.659 |
| **PDCD1** | IGFLR1 | CD8T | 0.658 |
| **PDCD1** | CCR5 | CD8T | 0.655 |
| **PDCD1** | APOBEC3C | CD8T | 0.655 |
| **PDCD1** | SUB1 | CD8T | 0.653 |
| **PDCD1** | SRGN | CD8T | 0.651 |
| **PDCD1** | PSMB3 | CD8T | 0.65 |
| **PDCD1** | CD8A | CD8T | 0.649 |
| **PDCD1** | CD79A | CD8T | 0.648 |
| **PDCD1** | CARHSP1 | CD8T | 0.647 |
| **PDCD1** | CLEC2B | CD8T | 0.646 |
| **PDCD1** | SDC4 | CD8T | 0.645 |
| **PDCD1** | MYL6 | CD8T | 0.643 |
| **PDCD1** | CAP1 | CD8T | 0.642 |
| **PDCD1** | ADGRE5 | CD8T | 0.642 |
| **PDCD1** | PDIA6 | CD8T | 0.641 |
| **PDCD1** | TPI1 | CD8T | 0.638 |
| **PDCD1** | IL2RG | CD8T | 0.637 |
| **PDCD1** | TSPO | CD8T | 0.635 |
| **PDCD1** | TXNDC17 | CD8T | 0.632 |
| **PDCD1** | HNRNPK | CD8T | 0.631 |

| Gene | Gene 2 | Cell Type | Correlation value |
|---|---|---|---|
| **PDCD1** | CYCS | CD8T | 0.63 |
| **PDCD1** | MCM5 | CD8T | 0.628 |
| **PDCD1** | CD2BP2 | CD8T | 0.628 |
| **PDCD1** | PRDX5 | CD8T | 0.628 |
| **PDCD1** | SERPINB9 | CD8T | 0.627 |
| **PDCD1** | ARPP19 | CD8T | 0.627 |
| **PDCD1** | KLRC2 | CD8T | 0.627 |
| **PDCD1** | NRBP1 | CD8T | 0.626 |
| **PDCD1** | SRP14 | CD8T | 0.626 |
| **PDCD1** | UCP2 | CD8T | 0.623 |
| **PDCD1** | IFI27 | CD8T | 0.622 |
| **PDCD1** | CLIC1 | CD8T | 0.622 |
| **PDCD1** | EID1 | CD8T | 0.621 |
| **PDCD1** | MCL1 | CD8T | 0.62 |
| **PDCD1** | MT1E | CD8T | 0.619 |
| **PDCD1** | HERPUD1 | CD8T | 0.618 |
| **PDCD1** | CCR1 | CD8T | 0.618 |
| **PDCD1** | PRDX1 | CD8T | 0.617 |
| **PDCD1** | TRBC1 | CD8T | 0.617 |
| **PDCD1** | HNRNPLL | CD8T | 0.616 |
| **PDCD1** | UBB | CD8T | 0.616 |
| **PDCD1** | SLC1A5 | CD8T | 0.615 |
| **PDCD1** | TMED9 | CD8T | 0.615 |
| **PDCD1** | ARPC2 | CD8T | 0.615 |
| **PDCD1** | NKG7 | CD8T | 0.613 |
| **PDCD1** | RGS1 | CD8T | 0.613 |
| **PDCD1** | PHPT1 | CD8T | 0.613 |
| **PDCD1** | SNU13 | CD8T | 0.612 |
| **PDCD1** | APOBEC3G | CD8T | 0.61 |
| **PDCD1** | NDUFB3 | CD8T | 0.609 |
| **PDCD1** | NUSAP1 | CD8T | 0.607 |
| **PDCD1** | COTL1 | CD8T | 0.607 |
| **PDCD1** | LGALS1 | CD8T | 0.606 |
| **PDCD1** | LINC01480 | CD8T | 0.606 |
| **PDCD1** | NDUFC1 | CD8T | 0.605 |
| **PDCD1** | KRT86 | CD8T | 0.604 |
| **PDCD1** | SURF4 | CD8T | 0.601 |

| Gene | Gene 2 | Cell Type | Correlation value |
|---|---|---|---|
| **PDCD1** | FDPS | CD8T | 0.6 |
| **PDCD1** | PMF1 | CD8T | 0.6 |
| **PDCD1** | CTSC | CD8T | 0.599 |
| **PDCD1** | LAMTOR5 | CD8T | 0.599 |
| **PDCD1** | NUDT5 | CD8T | 0.598 |
| **PDCD1** | KIF20B | CD8T | 0.598 |
| **PDCD1** | HMGN3 | CD8T | 0.597 |
| **PDCD1** | CD3D | CD8T | 0.596 |
| **PDCD1** | GZMA | CD8T | 0.595 |
| **PDCD1** | BUB3 | CD8T | 0.593 |
| **PDCD1** | HNRNPA3 | CD8T | 0.591 |
| **PDCD1** | DCTN3 | CD8T | 0.59 |
| **PDCD1** | MOB1A | CD8T | 0.589 |
| **PDCD1** | SLC25A3 | CD8T | 0.588 |
| **PDCD1** | SLC25A11 | CD8T | 0.588 |
| **PDCD1** | HMGN1 | CD8T | 0.588 |
| **PDCD1** | ITGB1 | CD8T | 0.586 |
| **PDCD1** | DEF6 | CD8T | 0.585 |
| **PDCD1** | RHOC | CD8T | 0.584 |
| **PDCD1** | CCDC28B | CD8T | 0.584 |
| **PDCD1** | COX5A | CD8T | 0.583 |
| **PDCD1** | U2AF1L4 | CD8T | 0.583 |
| **PDCD1** | MRPS34 | CD8T | 0.581 |
| **PDCD1** | MT1X | CD8T | 0.58 |
| **PDCD1** | IVNS1ABP | CD8T | 0.579 |
| **PDCD1** | GTF3C6 | CD8T | 0.578 |
| **PDCD1** | PSMC4 | CD8T | 0.575 |
| **PDCD1** | ANP32A | CD8T | 0.575 |
| **PDCD1** | SH2D1A | CD8T | 0.575 |
| **PDCD1** | SDF2L1 | CD8T | 0.574 |
| **PDCD1** | SKA2 | CD8T | 0.574 |
| **PDCD1** | CCNDBP1 | CD8T | 0.573 |
| **PDCD1** | AIF1 | CD8T | 0.57 |
| **PDCD1** | DNAJC8 | CD8T | 0.569 |
| **PDCD1** | COMMD7 | CD8T | 0.569 |
| **PDCD1** | PSMC3 | CD8T | 0.568 |
| **PDCD1** | ABI3 | CD8T | 0.568 |

| Gene | Gene 2 | Cell Type | Correlation value |
|---|---|---|---|
| **PDCD1** | CFL1 | CD8T | 0.567 |
| **PDCD1** | CYB5B | CD8T | 0.567 |
| **PDCD1** | PFDN2 | CD8T | 0.567 |
| **PDCD1** | CARD16 | CD8T | 0.567 |
| **PDCD1** | CAPZB | CD8T | 0.566 |
| **PDCD1** | SAP18 | CD8T | 0.566 |
| **PDCD1** | AP2S1 | CD8T | 0.565 |
| **PDCD1** | CST7 | CD8T | 0.565 |
| **PDCD1** | SERF2 | CD8T | 0.564 |
| **PDCD1** | GTF2A2 | CD8T | 0.564 |
| **PDCD1** | HSD17B10 | CD8T | 0.564 |
| **PDCD1** | ARL6IP1 | CD8T | 0.562 |
| **PDCD1** | COX6C | CD8T | 0.562 |
| **PDCD1** | B3GNT2 | CD8T | 0.561 |
| **PDCD1** | MSN | CD8T | 0.559 |
| **PDCD1** | ARHGEF1 | CD8T | 0.559 |
| **PDCD1** | TMCO1 | CD8T | 0.558 |
| **PDCD1** | MRPS11 | CD8T | 0.558 |
| **PDCD1** | EIF4E2 | CD8T | 0.558 |
| **PDCD1** | COX7A2 | CD8T | 0.557 |
| **PDCD1** | TRAPPC4 | CD8T | 0.557 |
| **PDCD1** | TXN | CD8T | 0.556 |
| **PDCD1** | EIF4H | CD8T | 0.554 |
| **PDCD1** | ENSA | CD8T | 0.554 |
| **PDCD1** | TMEM179B | CD8T | 0.553 |
| **PDCD1** | MRPL51 | CD8T | 0.552 |
| **PDCD1** | ITGB7 | CD8T | 0.552 |
| **PDCD1** | PSMA7 | CD8T | 0.552 |
| **PDCD1** | NDUFS6 | CD8T | 0.552 |
| **PDCD1** | UBE2A | CD8T | 0.551 |
| **PDCD1** | KPNA2 | CD8T | 0.55 |
| **PDCD1** | YIF1A | CD8T | 0.55 |
| **PDCD1** | PTPN7 | CD8T | 0.548 |
| **PDCD1** | CD8B | CD8T | 0.547 |
| **PDCD1** | RBX1 | CD8T | 0.547 |
| **PDCD1** | MLF2 | CD8T | 0.547 |
| **PDCD1** | KXD1 | CD8T | 0.546 |

| Gene | Gene 2 | Cell Type | Correlation value |
|---|---|---|---|
| **PDCD1** | GSTO1 | CD8T | 0.545 |
| **PDCD1** | HSPA5 | CD8T | 0.544 |
| **PDCD1** | ROMO1 | CD8T | 0.544 |
| **PDCD1** | UQCR10 | CD8T | 0.544 |
| **PDCD1** | PYCARD | CD8T | 0.543 |
| **PDCD1** | OAZ1 | CD8T | 0.542 |
| **PDCD1** | CMTM6 | CD8T | 0.541 |
| **PDCD1** | RNF181 | CD8T | 0.541 |
| **PDCD1** | DDA1 | CD8T | 0.539 |
| **PDCD1** | LSM8 | CD8T | 0.537 |
| **PDCD1** | RHBDD2 | CD8T | 0.537 |
| **PDCD1** | LCK | CD8T | 0.536 |
| **PDCD1** | PPP4C | CD8T | 0.535 |
| **PDCD1** | PSME2 | CD8T | 0.535 |
| **PDCD1** | H2AFZ | CD8T | 0.534 |
| **PDCD1** | MAP2K3 | CD8T | 0.534 |
| **PDCD1** | TALDO1 | CD8T | 0.534 |
| **PDCD1** | GGCT | CD8T | 0.534 |
| **PDCD1** | LBH | CD8T | 0.534 |
| **PDCD1** | ARPC1B | CD8T | 0.534 |
| **PDCD1** | CAPZA1 | CD8T | 0.534 |
| **PDCD1** | EDF1 | CD8T | 0.533 |
| **PDCD1** | SMC4 | CD8T | 0.532 |
| **PDCD1** | SLC16A3 | CD8T | 0.532 |
| **PDCD1** | DNAJB11 | CD8T | 0.53 |
| **PDCD1** | YWHAQ | CD8T | 0.53 |
| **PDCD1** | IDH2 | CD8T | 0.53 |
| **PDCD1** | BAX | CD8T | 0.53 |
| **PDCD1** | HSP90B1 | CD8T | 0.53 |
| **PDCD1** | COX6A1 | CD8T | 0.529 |
| **PDCD1** | PSMA2 | CD8T | 0.527 |
| **PDCD1** | KCNN4 | CD8T | 0.527 |
| **PDCD1** | SEC11A | CD8T | 0.526 |
| **PDCD1** | PSMA1 | CD8T | 0.526 |
| **PDCD1** | TANK | CD8T | 0.526 |
| **PDCD1** | RPS26 | CD8T | 0.525 |
| **PDCD1** | AP2M1 | CD8T | 0.525 |

| Gene | Gene 2 | Cell Type | Correlation value |
|---|---|---|---|
| **PDCD1** | GTF2H5 | CD8T | 0.525 |
| **PDCD1** | PEF1 | CD8T | 0.525 |
| **PDCD1** | set/07 | CD8T | 0.524 |
| **PDCD1** | TMX1 | CD8T | 0.523 |
| **PDCD1** | UQCRFS1 | CD8T | 0.522 |
| **PDCD1** | MPC2 | CD8T | 0.52 |
| **PDCD1** | SFT2D1 | CD8T | 0.519 |
| **PDCD1** | GLOD4 | CD8T | 0.519 |
| **PDCD1** | PDIA3 | CD8T | 0.518 |
| **PDCD1** | SEC61G | CD8T | 0.518 |
| **PDCD1** | CTSW | CD8T | 0.517 |
| **PDCD1** | HNRNPC | CD8T | 0.516 |
| **PDCD1** | LSM1 | CD8T | 0.516 |

**Table S9 –** List of genes significantly correlated with PD1 from BRCA_GSE114727_10X database

| Gene | Gene 2 | Cell Type | Correlation value |
|---|---|---|---|
| **PDCD1** | MIR155HG | CD8T | 0.851 |
| **PDCD1** | CXCL13 | CD8T | 0.835 |
| **PDCD1** | CTLA4 | CD8T | 0.827 |
| **PDCD1** | FABP5 | CD8T | 0.813 |
| **PDCD1** | ITGAE | CD8T | 0.798 |
| **PDCD1** | CCL3 | CD8T | 0.796 |
| **PDCD1** | ANXA5 | CD8T | 0.795 |
| **PDCD1** | AC069363.1 | CD8T | 0.788 |
| **PDCD1** | ACP5 | CD8T | 0.764 |
| **PDCD1** | SIRPG | CD8T | 0.763 |
| **PDCD1** | JAML | CD8T | 0.762 |
| **PDCD1** | PTTG1 | CD8T | 0.76 |
| **PDCD1** | RGS1 | CD8T | 0.747 |
| **PDCD1** | AC133644.2 | CD8T | 0.747 |
| **PDCD1** | AKAP5 | CD8T | 0.742 |
| **PDCD1** | CXCR6 | CD8T | 0.728 |
| **PDCD1** | LAYN | CD8T | 0.719 |
| **PDCD1** | IFNG | CD8T | 0.71 |
| **PDCD1** | TNFRSF1B | CD8T | 0.709 |
| **PDCD1** | PARK7 | CD8T | 0.709 |
| **PDCD1** | CTSD | CD8T | 0.707 |
| **PDCD1** | TIGIT | CD8T | 0.706 |
| **PDCD1** | COMMD7 | CD8T | 0.703 |
| **PDCD1** | LSP1 | CD8T | 0.703 |
| **PDCD1** | IFI27L2 | CD8T | 0.698 |
| **PDCD1** | PTMS | CD8T | 0.695 |
| **PDCD1** | GZMB | CD8T | 0.692 |
| **PDCD1** | CAPZB | CD8T | 0.689 |
| **PDCD1** | TNFRSF9 | CD8T | 0.689 |
| **PDCD1** | CARD16 | CD8T | 0.689 |
| **PDCD1** | WDR1 | CD8T | 0.684 |
| **PDCD1** | ALOX5AP | CD8T | 0.681 |
| **PDCD1** | SUB1 | CD8T | 0.681 |
| **PDCD1** | ITM2A | CD8T | 0.68 |
| **PDCD1** | RALY | CD8T | 0.679 |

| Gene | Gene 2 | Cell Type | Correlation value |
|---|---|---|---|
| **PDCD1** | AC017002.1 | CD8T | 0.678 |
| **PDCD1** | PPM1M | CD8T | 0.677 |
| **PDCD1** | BATF | CD8T | 0.677 |
| **PDCD1** | ITGB1 | CD8T | 0.677 |
| **PDCD1** | CD200 | CD8T | 0.672 |
| **PDCD1** | LAG3 | CD8T | 0.669 |
| **PDCD1** | CLEC2D | CD8T | 0.666 |
| **PDCD1** | BTG3 | CD8T | 0.665 |
| **PDCD1** | TXNDC17 | CD8T | 0.661 |
| **PDCD1** | SEC11A | CD8T | 0.66 |
| **PDCD1** | PKM | CD8T | 0.66 |
| **PDCD1** | SMC4 | CD8T | 0.66 |
| **PDCD1** | CD63 | CD8T | 0.659 |
| **PDCD1** | FKBP1A | CD8T | 0.658 |
| **PDCD1** | EVL | CD8T | 0.658 |
| **PDCD1** | IFI16 | CD8T | 0.658 |
| **PDCD1** | APOBEC3C | CD8T | 0.657 |
| **PDCD1** | HMGB1 | CD8T | 0.654 |
| **PDCD1** | ENSA | CD8T | 0.653 |
| **PDCD1** | KRT86 | CD8T | 0.653 |
| **PDCD1** | BLOC1S1 | CD8T | 0.65 |
| **PDCD1** | MSN | CD8T | 0.649 |
| **PDCD1** | CTSC | CD8T | 0.647 |
| **PDCD1** | PPM1G | CD8T | 0.647 |
| **PDCD1** | PSMD8 | CD8T | 0.646 |
| **PDCD1** | LCP2 | CD8T | 0.645 |
| **PDCD1** | NR3C1 | CD8T | 0.644 |
| **PDCD1** | GAPDH | CD8T | 0.643 |
| **PDCD1** | RHOA | CD8T | 0.643 |
| **PDCD1** | DUSP4 | CD8T | 0.639 |
| **PDCD1** | PDIA6 | CD8T | 0.638 |
| **PDCD1** | ARPC2 | CD8T | 0.633 |
| **PDCD1** | CARHSP1 | CD8T | 0.633 |
| **PDCD1** | APBB1IP | CD8T | 0.632 |
| **PDCD1** | TMX4 | CD8T | 0.631 |
| **PDCD1** | SLA | CD8T | 0.631 |
| **PDCD1** | TMEM120A | CD8T | 0.628 |

| Gene | Gene 2 | Cell Type | Correlation value |
|---|---|---|---|
| **PDCD1** | CD2BP2 | CD8T | 0.627 |
| **PDCD1** | POMP | CD8T | 0.626 |
| **PDCD1** | COX5A | CD8T | 0.626 |
| **PDCD1** | PYCARD | CD8T | 0.625 |
| **PDCD1** | DYNLL1 | CD8T | 0.624 |
| **PDCD1** | set/07 | CD8T | 0.623 |
| **PDCD1** | FIBP | CD8T | 0.622 |
| **PDCD1** | GZMH | CD8T | 0.622 |
| **PDCD1** | GBP2 | CD8T | 0.62 |
| **PDCD1** | LCK | CD8T | 0.618 |
| **PDCD1** | MZB1 | CD8T | 0.617 |
| **PDCD1** | GNAS | CD8T | 0.617 |
| **PDCD1** | ARHGAP9 | CD8T | 0.615 |
| **PDCD1** | CDCA7 | CD8T | 0.614 |
| **PDCD1** | CSK | CD8T | 0.614 |
| **PDCD1** | RAC1 | CD8T | 0.613 |
| **PDCD1** | MAP4K1 | CD8T | 0.612 |
| **PDCD1** | LINC01480 | CD8T | 0.611 |
| **PDCD1** | CAP1 | CD8T | 0.609 |
| **PDCD1** | PHYKPL | CD8T | 0.608 |
| **PDCD1** | TLN1 | CD8T | 0.608 |
| **PDCD1** | CD2 | CD8T | 0.608 |
| **PDCD1** | MYL6B | CD8T | 0.608 |
| **PDCD1** | CD3D | CD8T | 0.607 |
| **PDCD1** | PPP1CA | CD8T | 0.607 |
| **PDCD1** | GNAI2 | CD8T | 0.606 |
| **PDCD1** | HSBP1 | CD8T | 0.606 |
| **PDCD1** | CCND2 | CD8T | 0.605 |
| **PDCD1** | EID1 | CD8T | 0.604 |
| **PDCD1** | TADA3 | CD8T | 0.603 |
| **PDCD1** | DEK | CD8T | 0.603 |
| **PDCD1** | TWF2 | CD8T | 0.602 |
| **PDCD1** | PSMB8 | CD8T | 0.6 |
| **PDCD1** | CDK2AP2 | CD8T | 0.599 |
| **PDCD1** | COPZ1 | CD8T | 0.598 |
| **PDCD1** | COTL1 | CD8T | 0.597 |
| **PDCD1** | PDZD11 | CD8T | 0.597 |

| Gene | Gene 2 | Cell Type | Correlation value |
|---|---|---|---|
| **PDCD1** | BUB3 | CD8T | 0.597 |
| **PDCD1** | TSPO | CD8T | 0.596 |
| **PDCD1** | PARP1 | CD8T | 0.595 |
| **PDCD1** | PRF1 | CD8T | 0.593 |
| **PDCD1** | CHST12 | CD8T | 0.593 |
| **PDCD1** | PLEKHF1 | CD8T | 0.593 |
| **PDCD1** | ACTR3 | CD8T | 0.592 |
| **PDCD1** | SLC7A5 | CD8T | 0.592 |
| **PDCD1** | GNG5 | CD8T | 0.591 |
| **PDCD1** | AP2M1 | CD8T | 0.59 |
| **PDCD1** | SH3BP1 | CD8T | 0.589 |
| **PDCD1** | DYNLRB1 | CD8T | 0.589 |
| **PDCD1** | ARF5 | CD8T | 0.588 |
| **PDCD1** | UBE2L3 | CD8T | 0.586 |
| **PDCD1** | CSF1 | CD8T | 0.585 |
| **PDCD1** | CD3G | CD8T | 0.584 |
| **PDCD1** | UCP2 | CD8T | 0.584 |
| **PDCD1** | PAQR4 | CD8T | 0.581 |
| **PDCD1** | UBE2A | CD8T | 0.58 |
| **PDCD1** | RAB1B | CD8T | 0.579 |
| **PDCD1** | HNRNPA3 | CD8T | 0.578 |
| **PDCD1** | GEM | CD8T | 0.578 |
| **PDCD1** | AKIRIN2 | CD8T | 0.578 |
| **PDCD1** | PRDX3 | CD8T | 0.578 |
| **PDCD1** | CD82 | CD8T | 0.577 |
| **PDCD1** | ZCRB1 | CD8T | 0.576 |
| **PDCD1** | LAMTOR5 | CD8T | 0.575 |
| **PDCD1** | RNF187 | CD8T | 0.574 |
| **PDCD1** | PGAM1 | CD8T | 0.574 |
| **PDCD1** | WAS | CD8T | 0.574 |
| **PDCD1** | IDH2 | CD8T | 0.574 |
| **PDCD1** | MRPS36 | CD8T | 0.573 |
| **PDCD1** | CMTM6 | CD8T | 0.573 |
| **PDCD1** | ASNA1 | CD8T | 0.572 |
| **PDCD1** | RABAC1 | CD8T | 0.569 |
| **PDCD1** | TCIRG1 | CD8T | 0.569 |
| **PDCD1** | PPP4C | CD8T | 0.569 |

| Gene | Gene 2 | Cell Type | Correlation value |
|---|---|---|---|
| **PDCD1** | APOBEC3G | CD8T | 0.569 |
| **PDCD1** | EWSR1 | CD8T | 0.568 |
| **PDCD1** | SYNGR2 | CD8T | 0.568 |
| **PDCD1** | SERPINB1 | CD8T | 0.568 |
| **PDCD1** | OTUB1 | CD8T | 0.567 |
| **PDCD1** | TPM3 | CD8T | 0.567 |
| **PDCD1** | PSMB2 | CD8T | 0.567 |
| **PDCD1** | COPE | CD8T | 0.567 |
| **PDCD1** | ACTB | CD8T | 0.565 |
| **PDCD1** | ETFB | CD8T | 0.564 |
| **PDCD1** | NUDT21 | CD8T | 0.564 |
| **PDCD1** | MSC | CD8T | 0.563 |
| **PDCD1** | MRPL28 | CD8T | 0.563 |
| **PDCD1** | MPG | CD8T | 0.562 |
| **PDCD1** | NDUFB3 | CD8T | 0.562 |
| **PDCD1** | GBP5 | CD8T | 0.562 |
| **PDCD1** | TAP1 | CD8T | 0.562 |
| **PDCD1** | IL2RG | CD8T | 0.562 |
| **PDCD1** | RAB11B | CD8T | 0.561 |
| **PDCD1** | PSMB3 | CD8T | 0.561 |
| **PDCD1** | PDCD6 | CD8T | 0.56 |
| **PDCD1** | SUMO2 | CD8T | 0.559 |
| **PDCD1** | PSMB9 | CD8T | 0.558 |
| **PDCD1** | SIT1 | CD8T | 0.557 |
| **PDCD1** | HCLS1 | CD8T | 0.556 |
| **PDCD1** | SDF4 | CD8T | 0.555 |
| **PDCD1** | YWHAB | CD8T | 0.554 |
| **PDCD1** | AP2S1 | CD8T | 0.554 |
| **PDCD1** | BST2 | CD8T | 0.554 |
| **PDCD1** | AP1S1 | CD8T | 0.554 |
| **PDCD1** | NUDT5 | CD8T | 0.553 |
| **PDCD1** | VDAC1 | CD8T | 0.552 |
| **PDCD1** | IAH1 | CD8T | 0.551 |
| **PDCD1** | DBI | CD8T | 0.551 |
| **PDCD1** | MTHFD2 | CD8T | 0.55 |
| **PDCD1** | HNRNPLL | CD8T | 0.55 |
| **PDCD1** | DUSP23 | CD8T | 0.549 |

| Gene | Gene 2 | Cell Type | Correlation value |
|---|---|---|---|
| **PDCD1** | MAD2L2 | CD8T | 0.548 |
| **PDCD1** | LSM2 | CD8T | 0.548 |
| **PDCD1** | CD79B | CD8T | 0.548 |
| **PDCD1** | SRSF9 | CD8T | 0.547 |
| **PDCD1** | ZYX | CD8T | 0.547 |
| **PDCD1** | PRDM1 | CD8T | 0.547 |
| **PDCD1** | LY6E | CD8T | 0.547 |
| **PDCD1** | RBX1 | CD8T | 0.546 |
| **PDCD1** | NAP1L4 | CD8T | 0.546 |
| **PDCD1** | MDH2 | CD8T | 0.545 |
| **PDCD1** | BIN1 | CD8T | 0.544 |
| **PDCD1** | CHMP2A | CD8T | 0.544 |
| **PDCD1** | PTPN7 | CD8T | 0.543 |
| **PDCD1** | UBE2D2 | CD8T | 0.543 |
| **PDCD1** | SRGN | CD8T | 0.543 |
| **PDCD1** | FASLG | CD8T | 0.542 |
| **PDCD1** | GGCT | CD8T | 0.542 |
| **PDCD1** | TPI1 | CD8T | 0.541 |
| **PDCD1** | CTSB | CD8T | 0.541 |
| **PDCD1** | VAMP5 | CD8T | 0.541 |
| **PDCD1** | ID2 | CD8T | 0.541 |
| **PDCD1** | MIEN1 | CD8T | 0.54 |
| **PDCD1** | NDUFA6 | CD8T | 0.54 |
| **PDCD1** | ZBED2 | CD8T | 0.54 |
| **PDCD1** | CFL1 | CD8T | 0.54 |
| **PDCD1** | IL2RB | CD8T | 0.54 |
| **PDCD1** | GSTO1 | CD8T | 0.54 |
| **PDCD1** | LGALS3 | CD8T | 0.538 |
| **PDCD1** | TMEM179B | CD8T | 0.537 |
| **PDCD1** | POLR2G | CD8T | 0.537 |
| **PDCD1** | HSD17B10 | CD8T | 0.536 |
| **PDCD1** | NELFCD | CD8T | 0.536 |
| **PDCD1** | MKI67 | CD8T | 0.536 |
| **PDCD1** | NDUFS8 | CD8T | 0.535 |
| **PDCD1** | MIIP | CD8T | 0.534 |
| **PDCD1** | TYMP | CD8T | 0.534 |
| **PDCD1** | SLC1A5 | CD8T | 0.534 |

| Gene | Gene 2 | Cell Type | Correlation value |
|---|---|---|---|
| **PDCD1** | NABP2 | CD8T | 0.533 |
| **PDCD1** | CKS2 | CD8T | 0.533 |
| **PDCD1** | FLII | CD8T | 0.533 |
| **PDCD1** | SUMO1 | CD8T | 0.533 |
| **PDCD1** | UBE2L6 | CD8T | 0.533 |
| **PDCD1** | EZR | CD8T | 0.533 |
| **PDCD1** | CLIC1 | CD8T | 0.532 |
| **PDCD1** | PIN1 | CD8T | 0.532 |
| **PDCD1** | TIMM8B | CD8T | 0.532 |
| **PDCD1** | BSG | CD8T | 0.531 |
| **PDCD1** | ADGRE5 | CD8T | 0.531 |
| **PDCD1** | PRKAR1A | CD8T | 0.531 |
| **PDCD1** | GTF3C6 | CD8T | 0.531 |
| **PDCD1** | ARPC3 | CD8T | 0.531 |
| **PDCD1** | B2M | CD8T | 0.53 |
| **PDCD1** | ZNHIT1 | CD8T | 0.53 |
| **PDCD1** | CALM3 | CD8T | 0.529 |
| **PDCD1** | NDUFC1 | CD8T | 0.529 |
| **PDCD1** | MCM5 | CD8T | 0.528 |
| **PDCD1** | PSMA4 | CD8T | 0.528 |
| **PDCD1** | YWHAQ | CD8T | 0.528 |
| **PDCD1** | CCDC167 | CD8T | 0.528 |
| **PDCD1** | UBE2N | CD8T | 0.527 |
| **PDCD1** | RBCK1 | CD8T | 0.527 |
| **PDCD1** | NDUFS6 | CD8T | 0.527 |
| **PDCD1** | NCF4 | CD8T | 0.526 |
| **PDCD1** | MYL6 | CD8T | 0.526 |
| **PDCD1** | CALR | CD8T | 0.526 |
| **PDCD1** | TPM4 | CD8T | 0.526 |
| **PDCD1** | GPI | CD8T | 0.526 |
| **PDCD1** | VASP | CD8T | 0.525 |
| **PDCD1** | PSMB6 | CD8T | 0.525 |
| **PDCD1** | AIP | CD8T | 0.524 |
| **PDCD1** | PPP1R7 | CD8T | 0.524 |
| **PDCD1** | TBCB | CD8T | 0.523 |
| **PDCD1** | CD151 | CD8T | 0.523 |
| **PDCD1** | SCAND1 | CD8T | 0.522 |

| Gene | Gene 2 | Cell Type | Correlation value |
|---|---|---|---|
| **PDCD1** | COMMD8 | CD8T | 0.522 |
| **PDCD1** | LAMTOR1 | CD8T | 0.522 |
| **PDCD1** | TMX1 | CD8T | 0.522 |
| **PDCD1** | CCR5 | CD8T | 0.522 |
| **PDCD1** | PRDX6 | CD8T | 0.521 |
| **PDCD1** | FERMT3 | CD8T | 0.52 |
| **PDCD1** | LPXN | CD8T | 0.52 |
| **PDCD1** | REEP5 | CD8T | 0.519 |
| **PDCD1** | PSMD4 | CD8T | 0.519 |
| **PDCD1** | CLTB | CD8T | 0.519 |
| **PDCD1** | SLTM | CD8T | 0.518 |
| **PDCD1** | SNRPB2 | CD8T | 0.518 |

**Table S10 –** List of genes significantly correlated with PD1 from BRCA_EMTAB8107 database

| Gene | Gene 2 | Cell Type | Correlation value |
|---|---|---|---|
| **PDCD1** | CXCL13 | CD8T | 0.869 |
| **PDCD1** | TIGIT | CD8T | 0.821 |
| **PDCD1** | CXCR6 | CD8T | 0.806 |
| **PDCD1** | LAG3 | CD8T | 0.798 |
| **PDCD1** | PTMS | CD8T | 0.796 |
| **PDCD1** | IFNG | CD8T | 0.784 |
| **PDCD1** | DUSP4 | CD8T | 0.775 |
| **PDCD1** | MIR155HG | CD8T | 0.765 |
| **PDCD1** | PHLDA1 | CD8T | 0.76 |
| **PDCD1** | AC069363.1 | CD8T | 0.745 |
| **PDCD1** | CSF1 | CD8T | 0.737 |
| **PDCD1** | RGS1 | CD8T | 0.737 |
| **PDCD1** | PLPP1 | CD8T | 0.723 |
| **PDCD1** | SIRPG | CD8T | 0.721 |
| **PDCD1** | CAMK1 | CD8T | 0.719 |
| **PDCD1** | ACP5 | CD8T | 0.717 |
| **PDCD1** | LINC01480 | CD8T | 0.704 |
| **PDCD1** | MTHFD2 | CD8T | 0.704 |
| **PDCD1** | ZBED2 | CD8T | 0.699 |
| **PDCD1** | PTPN7 | CD8T | 0.696 |
| **PDCD1** | PTTG1 | CD8T | 0.696 |
| **PDCD1** | CD200 | CD8T | 0.693 |
| **PDCD1** | TYMP | CD8T | 0.691 |
| **PDCD1** | GAPDH | CD8T | 0.688 |
| **PDCD1** | SUB1 | CD8T | 0.68 |
| **PDCD1** | CCND2 | CD8T | 0.676 |
| **PDCD1** | FABP5 | CD8T | 0.676 |
| **PDCD1** | CD2 | CD8T | 0.663 |
| **PDCD1** | ANXA5 | CD8T | 0.663 |
| **PDCD1** | TNFRSF1B | CD8T | 0.659 |
| **PDCD1** | AC133644.2 | CD8T | 0.659 |
| **PDCD1** | IGFLR1 | CD8T | 0.654 |
| **PDCD1** | TNFRSF9 | CD8T | 0.651 |
| **PDCD1** | CLEC2D | CD8T | 0.649 |
| **PDCD1** | PGAM1 | CD8T | 0.645 |

| Gene | Gene 2 | Cell Type | Correlation value |
|---|---|---|---|
| **PDCD1** | PDIA6 | CD8T | 0.642 |
| **PDCD1** | ALOX5AP | CD8T | 0.641 |
| **PDCD1** | FKBP1A | CD8T | 0.641 |
| **PDCD1** | CD74 | CD8T | 0.638 |
| **PDCD1** | KRT86 | CD8T | 0.637 |
| **PDCD1** | LY6E | CD8T | 0.636 |
| **PDCD1** | ITM2A | CD8T | 0.635 |
| **PDCD1** | CTLA4 | CD8T | 0.633 |
| **PDCD1** | TAP1 | CD8T | 0.632 |
| **PDCD1** | CLTA | CD8T | 0.629 |
| **PDCD1** | PKM | CD8T | 0.627 |
| **PDCD1** | GEM | CD8T | 0.627 |
| **PDCD1** | BST2 | CD8T | 0.622 |
| **PDCD1** | HMOX1 | CD8T | 0.617 |
| **PDCD1** | CCL3 | CD8T | 0.617 |
| **PDCD1** | APOBEC3C | CD8T | 0.614 |
| **PDCD1** | LAP3 | CD8T | 0.614 |
| **PDCD1** | GNG5 | CD8T | 0.612 |
| **PDCD1** | IFI16 | CD8T | 0.612 |
| **PDCD1** | DYNLL1 | CD8T | 0.611 |
| **PDCD1** | MYL6B | CD8T | 0.608 |
| **PDCD1** | CD82 | CD8T | 0.607 |
| **PDCD1** | OAS1 | CD8T | 0.606 |
| **PDCD1** | IFI27L2 | CD8T | 0.606 |
| **PDCD1** | IFI6 | CD8T | 0.602 |
| **PDCD1** | SH2D2A | CD8T | 0.602 |
| **PDCD1** | BTG3 | CD8T | 0.601 |
| **PDCD1** | TPI1 | CD8T | 0.6 |
| **PDCD1** | CAPZB | CD8T | 0.599 |
| **PDCD1** | BATF | CD8T | 0.596 |
| **PDCD1** | SRGN | CD8T | 0.595 |
| **PDCD1** | CCDC50 | CD8T | 0.594 |
| **PDCD1** | ISG15 | CD8T | 0.591 |
| **PDCD1** | MT1E | CD8T | 0.591 |
| **PDCD1** | ARPC2 | CD8T | 0.589 |
| **PDCD1** | PARK7 | CD8T | 0.589 |
| **PDCD1** | PSMB9 | CD8T | 0.585 |

| Gene | Gene 2 | Cell Type | Correlation value |
|---|---|---|---|
| **PDCD1** | ITGB1 | CD8T | 0.584 |
| **PDCD1** | MT2A | CD8T | 0.584 |
| **PDCD1** | ID2 | CD8T | 0.584 |
| **PDCD1** | COX5A | CD8T | 0.583 |
| **PDCD1** | CTSC | CD8T | 0.583 |
| **PDCD1** | GBP5 | CD8T | 0.581 |
| **PDCD1** | PSMB8 | CD8T | 0.58 |
| **PDCD1** | ARPC1B | CD8T | 0.579 |
| **PDCD1** | CORO1B | CD8T | 0.578 |
| **PDCD1** | RHOA | CD8T | 0.575 |
| **PDCD1** | PPM1M | CD8T | 0.569 |
| **PDCD1** | CKS2 | CD8T | 0.569 |
| **PDCD1** | MX1 | CD8T | 0.569 |
| **PDCD1** | RANBP1 | CD8T | 0.564 |
| **PDCD1** | RHOB | CD8T | 0.562 |
| **PDCD1** | IFI44 | CD8T | 0.56 |
| **PDCD1** | FAM166B | CD8T | 0.56 |
| **PDCD1** | POMP | CD8T | 0.559 |
| **PDCD1** | CD2BP2 | CD8T | 0.558 |
| **PDCD1** | VAMP5 | CD8T | 0.558 |
| **PDCD1** | PHPT1 | CD8T | 0.557 |
| **PDCD1** | PLSCR1 | CD8T | 0.557 |
| **PDCD1** | TXNDC17 | CD8T | 0.555 |
| **PDCD1** | MAP1LC3A | CD8T | 0.554 |
| **PDCD1** | IFI35 | CD8T | 0.553 |
| **PDCD1** | IRF7 | CD8T | 0.553 |
| **PDCD1** | UBE2L6 | CD8T | 0.552 |
| **PDCD1** | CALM3 | CD8T | 0.549 |
| **PDCD1** | NBL1 | CD8T | 0.548 |
| **PDCD1** | OASL | CD8T | 0.546 |
| **PDCD1** | LCP2 | CD8T | 0.544 |
| **PDCD1** | JAML | CD8T | 0.544 |
| **PDCD1** | YWHAH | CD8T | 0.541 |
| **PDCD1** | ZFP36L1 | CD8T | 0.539 |
| **PDCD1** | SH3BGRL3 | CD8T | 0.539 |
| **PDCD1** | RBCK1 | CD8T | 0.538 |
| **PDCD1** | SUMO2 | CD8T | 0.538 |

| Gene | Gene 2 | Cell Type | Correlation value |
|---|---|---|---|
| **PDCD1** | HSBP1 | CD8T | 0.538 |
| **PDCD1** | FASLG | CD8T | 0.538 |
| **PDCD1** | TMEM14A | CD8T | 0.537 |
| **PDCD1** | SHISA5 | CD8T | 0.537 |
| **PDCD1** | PDLIM7 | CD8T | 0.536 |
| **PDCD1** | NDUFV2 | CD8T | 0.535 |
| **PDCD1** | COTL1 | CD8T | 0.534 |
| **PDCD1** | NCF4 | CD8T | 0.534 |
| **PDCD1** | SAMD9L | CD8T | 0.534 |
| **PDCD1** | ICOS | CD8T | 0.534 |
| **PDCD1** | CDKN2A | CD8T | 0.533 |
| **PDCD1** | CARD16 | CD8T | 0.533 |
| **PDCD1** | SDC4 | CD8T | 0.53 |
| **PDCD1** | CSF2 | CD8T | 0.53 |
| **PDCD1** | VDAC1 | CD8T | 0.528 |
| **PDCD1** | SURF4 | CD8T | 0.527 |
| **PDCD1** | CD3D | CD8T | 0.527 |
| **PDCD1** | SERF2 | CD8T | 0.526 |
| **PDCD1** | COMMD7 | CD8T | 0.525 |
| **PDCD1** | TPM4 | CD8T | 0.525 |
| **PDCD1** | C4orf48 | CD8T | 0.523 |
| **PDCD1** | TNFRSF18 | CD8T | 0.522 |
| **PDCD1** | CCR1 | CD8T | 0.522 |
| **PDCD1** | LBH | CD8T | 0.522 |
| **PDCD1** | RBX1 | CD8T | 0.521 |
| **PDCD1** | DEK | CD8T | 0.521 |
| **PDCD1** | NPDC1 | CD8T | 0.52 |
| **PDCD1** | DRAP1 | CD8T | 0.52 |
| **PDCD1** | KLHDC7B | CD8T | 0.519 |
| **PDCD1** | B2M | CD8T | 0.518 |
| **PDCD1** | MZB1 | CD8T | 0.518 |
| **PDCD1** | WDR1 | CD8T | 0.518 |
| **PDCD1** | C3orf14 | CD8T | 0.517 |
| **PDCD1** | SCAMP2 | CD8T | 0.517 |
| **PDCD1** | PPP1CC | CD8T | 0.516 |
| **PDCD1** | SNRPG | CD8T | 0.516 |
| **PDCD1** | PLEKHF1 | CD8T | 0.514 |

**Table S11 –** List of genes significantly correlated with PD1 from BRCA_GSE161529 database

| Gene | Gene 2 | Cell Type | Correlation value |
|---|---|---|---|
| **PDCD1** | TIGIT | CD8T | 0.758 |
| **PDCD1** | CD3D | CD8T | 0.756 |
| **PDCD1** | CD2 | CD8T | 0.749 |
| **PDCD1** | TRBC1 | CD8T | 0.741 |
| **PDCD1** | TRAC | CD8T | 0.721 |
| **PDCD1** | LCK | CD8T | 0.712 |
| **PDCD1** | CTLA4 | CD8T | 0.698 |
| **PDCD1** | IFNG | CD8T | 0.696 |
| **PDCD1** | TRBC2 | CD8T | 0.693 |
| **PDCD1** | CD3G | CD8T | 0.683 |
| **PDCD1** | ZBED2 | CD8T | 0.682 |
| **PDCD1** | CD247 | CD8T | 0.679 |
| **PDCD1** | GPR171 | CD8T | 0.665 |
| **PDCD1** | LAIR2 | CD8T | 0.65 |
| **PDCD1** | CXCR6 | CD8T | 0.648 |
| **PDCD1** | CD7 | CD8T | 0.644 |
| **PDCD1** | SPOCK2 | CD8T | 0.636 |
| **PDCD1** | CD3E | CD8T | 0.636 |
| **PDCD1** | SH2D2A | CD8T | 0.635 |
| **PDCD1** | IL2RB | CD8T | 0.623 |
| **PDCD1** | ICOS | CD8T | 0.621 |
| **PDCD1** | PTPN7 | CD8T | 0.62 |
| **PDCD1** | CLEC2D | CD8T | 0.609 |
| **PDCD1** | APOBEC3G | CD8T | 0.607 |
| **PDCD1** | SIT1 | CD8T | 0.607 |
| **PDCD1** | KLRB1 | CD8T | 0.6 |
| **PDCD1** | TBC1D10C | CD8T | 0.591 |
| **PDCD1** | LINC01943 | CD8T | 0.568 |
| **PDCD1** | GZMM | CD8T | 0.566 |
| **PDCD1** | S1PR4 | CD8T | 0.564 |

| Gene | Gene 2 | Cell Type | Correlation value |
|---|---|---|---|
| **PDCD1** | CD8A | CD8T | 0.564 |
| **PDCD1** | TNFRSF9 | CD8T | 0.557 |
| **PDCD1** | CD27 | CD8T | 0.555 |
| **PDCD1** | CD8B | CD8T | 0.553 |
| **PDCD1** | CXCR3 | CD8T | 0.552 |
| **PDCD1** | GZMA | CD8T | 0.549 |
| **PDCD1** | IL2RG | CD8T | 0.549 |
| **PDCD1** | RAC2 | CD8T | 0.543 |
| **PDCD1** | CTSW | CD8T | 0.537 |
| **PDCD1** | CORO1A | CD8T | 0.536 |
| **PDCD1** | LAT | CD8T | 0.523 |
